# Prdm16 amplifies Notch signaling and suppresses venous lineage specification to prevent arteriovenous malformations during vascular development

**DOI:** 10.1101/2021.12.05.471275

**Authors:** Jore Van Wauwe, Sander Craps, KC Ashmita, Lisabel Asuncion, Pieter Vrancaert, Margo Daems, Chris Finck, Shane Wright, Christian Müller, Nicholas D. Leigh, Maike Frye, Calum A. MacRae, Aernout Luttun, Manu Beerens

## Abstract

Proper arterial versus venous endothelial lineage identity is required to establish a hierarchical network of arteries and veins and prevent the occurrence of life-threatening arteriovenous malformations. The molecular mechanisms that govern arteriovenous lineage specification remain however incompletely understood. Here, we demonstrate that the transcription factor Prdm16 is expressed in arterial but not venous endothelial cells (ECs) from the earliest stages of development, where it actively promotes arterial EC identity by enhancing canonical Notch activity, while simultaneously suppressing the alternative venous cell fate. Concordantly, our results in zebrafish show that Prdm16 coordinates proper arterial development and arteriovenous lineage specification together with canonical Notch signaling, as combined loss of prdm16 and notch in zebrafish invariably leads to arteriovenous malformations (AVMs). Although the arterializing effect of Prdm16 in human ECs is dependent on the absolute levels of the cleaved intracellular domain of the Notch receptors (NICD), Prdm16 does not increase NICD levels per se. Rather, Prdm16 physically and functionally interacts with NICD to potentiate its effect. Prdm16 further finetunes Notch signaling and arterial development by complexing with the Notch downstream effector Hey2, which regulates arterial lineage specification and development across species. Together, our data demonstrate that Prdm16 act as a rheostat for endothelial Notch activity and suggest that Prdm16 signaling may constitute a novel therapeutic target for AVMs.

## Introduction

Developing a distinctive network of arteries and veins is essential to ensure arterial supply of oxygen and nutrient-rich blood to all tissues and venous return of oxygen-deprived blood back to the heart. Endothelial cells (ECs) lining the inside of all blood vessels are critical for both blood vessel development and homeostasis. Their further specification into cells of the arterial or venous lineage is essential for proper vascular development and function and is regulated by diverse signaling cascades that converge on transcription factors (TFs). Venous identity is promoted by chicken ovalbumin upstream TF 2 (COUP-TFII) expressed in venous but not arterial ECs, where it actively suppresses canonical NOTCH signaling.^1–3^ NOTCH is initiated by binding of Delta-like ligand 4 (DLL4) to endothelial NOTCH receptors, which triggers the release and nuclear translocation of the NOTCH receptor intracellular domain (NICD). Within the nucleus, NICD associates with DNA-bound RBPJ_*k*_ to induce canonical NOTCH activity which drives arterial EC specification in zebrafish,^4–9^ mice,^10–12^ and human cells.^13,14^ This is largely mediated via NOTCH downstream effector TFs, Hes-Related Family basic helix-loop-helix (bHLH) Transcription Factor with YRPW Motif (HEY)1 and HEY2, as evident by the reduced caliber of arterial vessels and failure to establish proper arterial cell fate in compound *Hey1/Hey2* knockout mice.^10^ Importantly, perturbed arteriovenous (AV) differentiation can lead to life-threatening AV malformations (AVMs) or shunts.^15^ Gaining more insight into the molecular mechanisms that govern AV endothelial specification could thus identify new druggable targets to treat AVMs, and inform tissue engineering strategies to more faithfully recapitulate physiological vessel phenotypes.

Positive regulatory domain-containing protein (PRDM) 16 belongs to a TF family which coordinates binary cell fate decisions in nematodes, fruit flies, and mammals.^16^ In adipose tissue, Prdm16 is exclusively expressed by and regulates the transcriptional program of brown/beige adipocytes, while concomitantly suppressing white adipose-specific gene expression.^17,18^ *Vice versa*, COUP-TFII drives white adipogenesis with *Coup-TFII^+/-^*mice displaying a marked upregulation of brown adipose-selective genes in white fat deposits.^19^ This mutually exclusive expression pattern of PRDM16 and COUP-TFII is mirrored in the heart, where they are expressed by and control lineage specification of ventricular^20–22^ and atrial^23^ cardiomyocytes, respectively. Accordingly, a wide range of cardiac anomalies have been observed in mice with deletion of *Prdm16* in cardiomyocytes or their progenitors,^20,22,24,25^ and in zebrafish with perturbed prdm16 signaling.^21^ Prdm16’s role in the circulatory system is however not restricted to the heart, as others^27–29^ and we^26^ have documented pronounced Prdm16 expression in arterial smooth muscle cells, where its decreased expression has been linked to atherogenesis and aneurysm formation.^27,30^ Additionally, genomic variants in the *PRDM16* locus have been associated with coronary artery disease (CAD),^28,31^ further supporting an important role for PRDM16 during cardiovascular development and function, and illustrating that impaired PRDM16 levels may predispose to cardiovascular disorders.

Recent evidence indicated that Prdm16 controls angiogenesis, i.e. the growth of new blood vessels from existing ones. Knockdown (KD) of *Prdm16* impairs angiogenesis in zebrafish and reduces migration of human induced pluripotent stem cell-derived ECs,^32^ while EC-specific deletion of *Prdm16* in neonatal mice perturbs retinal angiogenesis.^29^ Additionally, we previously showed that Prdm16 is expressed in arterial but not venous ECs of the adult mouse and human vasculature.^26,33^ Concordantly, we documented impaired perfusion recovery and endothelial dysfunction in mice with postnatal deletion of endothelial *Prdm16* in a hindlimb ischemia model,^26^ which faithfully recapitulates several aspects of human thrombotic disease. Together, these studies illustrate the importance of endothelial Prdm16 for vascular sprouting and homeostasis and suggest that the association of PRDM16 with CAD^31^ might be at least partly due to its endothelial function. Thompson *et al*. further revealed that the developing arteries of EC-specific *Prdm16* knockout mice display slightly increased protein levels of the venous marker Endomucin (EMCN) and reduced smooth muscle cell coating,^29^ the latter echoing our findings in adult mice with EC-specific deletion of *Prdm16.*^26^ Although these studies suggest a potential role for endothelial Prdm16 during vascular development and homeostasis, the precise function of PRDM16 and its downstream molecular mechanisms within ECs remains incompletely understood.

Here, we show that *Prdm16* is expressed in arterial ECs from zebrafish and mouse embryos from the earliest developmental stages onwards. Consistent with a role for Prdm16 during AV endothelial lineage specification, we found that Prdm16 suppresses the venous transcriptional program while boosting the arterial gene signature in human endothelial (progenitor) cells via increasing canonical Notch activity. *Prdm16* KD in zebrafish resulted in AVMs which were rescued by simultaneous overexpression of mouse Prdm16 in the endothelium, indicating that they are the result of reduced prdm16-mediated signaling within ECs. Partial inhibition of canonical notch sensitized *prdm16* KD embryos to develop AVMs, while incomplete *prdm16* KD resulted in shunt formation in otherwise phenotypically normal zebrafish embryos heterozygous for the *grl^m145^*(hey2) hypomorph,^7^ indicating that prdm16 and notch cooperatively regulate vascular development. Mechanistically, PRDM16 amplified canonical NOTCH signaling via its physical interaction with both NICD and HEY2, in line with the strong functional interplay between Prdm16 and Notch^20,34,35^ and the physical interaction between PRDM and bHLH TFs observed in non-vascular tissues.^36,37,38^ Together, our results unravel a critical role for Prdm16 during AV endothelial lineage specification, mediated via its modulatory effect on canonical Notch activity in ECs, and highlight that PRDM16 signaling may be a novel therapeutic target to treat AVMs.

## Results

### Prdm16 is part of a conserved arterial-specific endothelial gene signature

To identify TFs that regulate arterial EC specification and development, we performed an intersectional analysis on our transcriptomic dataset of freshly isolated human arterial and venous ECs^33^ and a newly collected dataset of fluorescence activated cell sorting (FACS) isolated adult mouse arterial and venous ECs (**Fig.S1A**). This analysis revealed a common set of 52 genes (FDR ≤ 0.05 and logFC ≥ |1| in both species) marking the AV identity of mature ECs (**TableS1**). These genes represented key physiological processes implicated in vascular development including ‘circulatory system development’, ‘blood vessel morphogenesis’, ‘artery development’, ‘aorta development’, ‘EC migration’, and ‘cell fate commitment’ (**TableS2**). Our list contained key drivers of arterial (Hey2) and venous (Coup-TFII) EC lineage specification, suggesting enrichment for candidate genes implicated in AV EC differentiation processes. Among the additional 21 and 29 arterial- and venous-specific genes, respectively, we identified the TF Prdm16 as a genuine marker for arterial ECs (**Fig.S1A**). We confirmed its unique arterial expression pattern on independently sorted mouse and human arterial and venous ECs (**Fig.1A,B**), and by PRDM16 immunohistochemistry in the human umbilical cord (**Fig.1C**). β-galactosidase expression in *Prdm16^lacZ/+^* reporter mouse embryos^39^ further revealed that *Prdm16* is already robustly expressed by ECs of the paired dorsal aorta (DA) at embryonic day (E) 9.5 (**Fig.1D,E**). Prdm16 levels were also markedly higher in the umbilical and vitelline artery compared to their venous counterparts, illustrating that the arterial enrichment of Prdm16 is not restricted to intra-embryonic vessels (**Fig.1F,G**). At E10.5, Prdm16 expression remained present in the DA, the intersomitic vessels (ISVs) sprouting from it, and in the intracranial arteries (**Fig.1H-K**). Arterial ECs from the E14.5 jugular and intercostal artery also showed robust Prdm16 immunoreactivity, which was absent in ECs from the jugular and intercostal vein (**Fig.1L,M**). Finally, Prdm16 was readily detected in ECs from the aorta and the basilar artery at E14.5 (**Fig.1N,O**), and in ECs of the coronary artery but not vein at E17.5 (**Fig.1P,Q**). In agreement with these findings, publicly available (single-cell) RNA-sequencing datasets (Gtex portal) highlighted prominent *Prdm16* mRNA levels in arterial tissue and blood but not lymphatic ECs (**Fig.S1B,C**). Together, these data indicate that – within the endothelium – Prdm16 is uniquely expressed by arterial ECs independent of species, hemodynamic forces, oxygen tension, anatomical location, or developmental stage.

**Figure 1.**
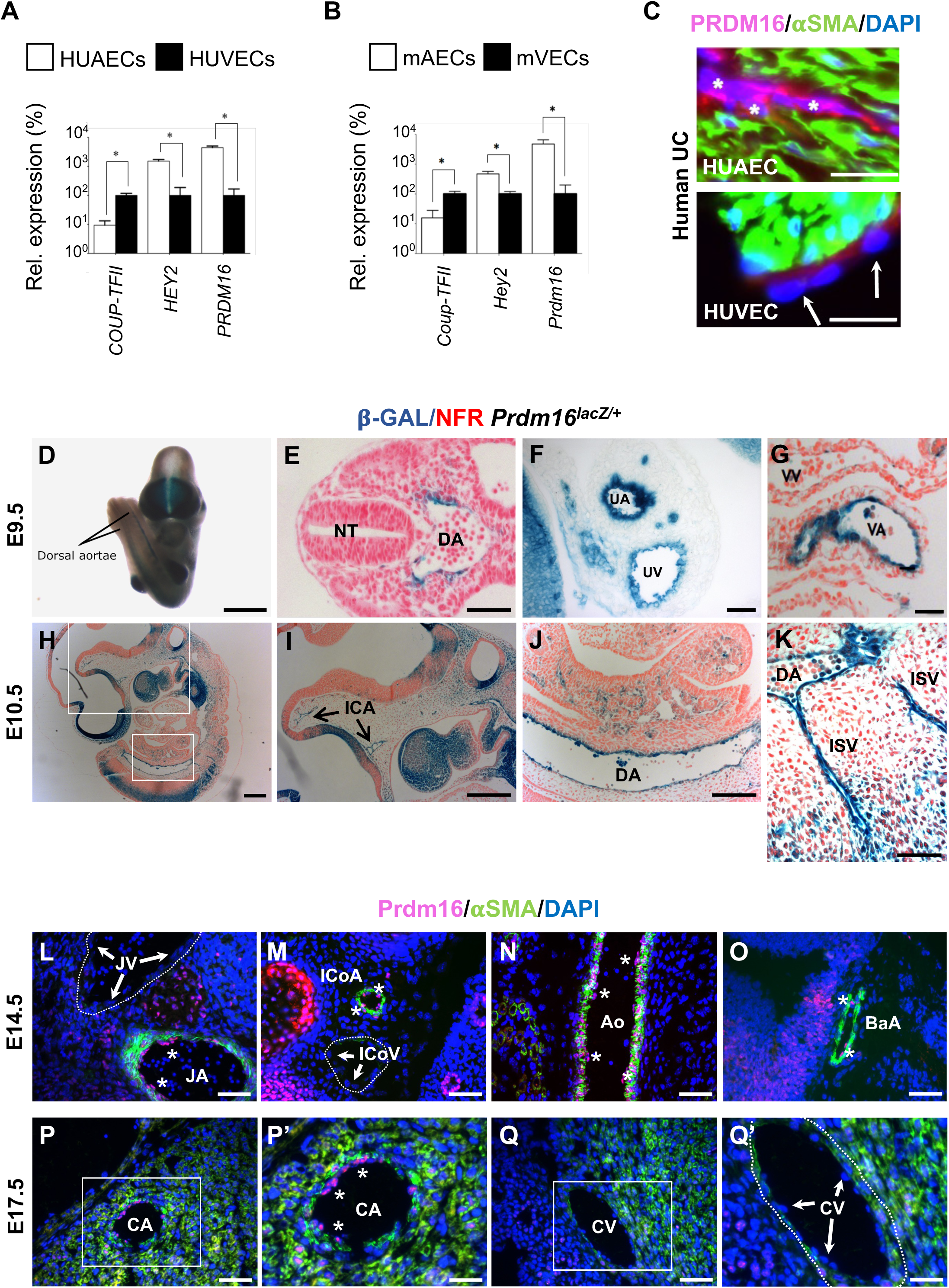
Endothelial *prdm16* is required for proper vascular development in zebrafish. **(A,B)** *Hey2*, *Coup-TFII*, and *Prdm16* mRNA levels on freshly isolated arterial (white bars) and venous (black bars) from human (A) and mouse (B) adult ECs, demonstrating that *Hey2* and *Prdm16* are enriched in arterial versus venous ECs, while the opposite is true for *Coup-TFII*. \**P*<0.05 by paired Student’s *t*-test**. (C)** Immunofluorescence (IF) staining for Prdm16 (magenta), α-smooth muscle actin (αSMA; green) and cell nuclei (DAPI; blue) on cross-sections from the human umbilical cord, indicating the presence of PRDM16 on HUAECs (white asterisks) but not HUVECs (white arrows). **(D-K)** Whole embryo view (D), cross-sections (E-G) or sagittal sections (H-K) of β-galactosidase (β-GAL) activity staining on embryonic day (E)9.5 (D-G) or E10.5 (H-K) embryos of the Prdm16 reporter mouse strain (*Prdm16^Gt683Lex^*). NFR counterstaining was used in E,G, H-K. NT: neural tube; DA: dorsal aorta; UA: umbilical artery; UV: umbilical vein; VA: vitelline artery; VV: vitelline vein; ICA: intracranial artery; ISV: intersomitic vessel. **(L-Q)** IF staining for Prdm16 (magenta), α-smooth muscle actin (αSMA; green) and cell nuclei (DAPI; blue) on cross-sections of E14.5 (L-O) or E17.5 (P-Q’) FvB mouse embryos. Asterisks indicate Prdm16-expressing arterial ECs, white arrows Prdm16-negative venous ECs. Veins are lined with white dotted lines in L,M, Q’. JA: jugular artery; JV: jugular vein; ICoA: intercostal artery; ICoV: intercostal vein; Ao: aorta; BaA: basilar artery; CA: coronary artery; CV: coronary vein. Scale bars: 1mm in D; 100μm in E,H-K,M,N; 50μm in L-Q; 25μm in P’,Q’.

### Prdm16 controls early arterial development in zebrafish

We then analyzed the vascular expression pattern of *prdm16* in zebrafish using whole mount *in situ* hybridization (WISH) and the *tud28Gt prdm16* reporter.^40^ The arterial-specific expression of Prdm16 was conserved in non-mammalian vertebrates, as *prdm16* was present in the DA, but not the posterior cardinal vein (PCV) of one-, 2-, and 6-day-old zebrafish embryos, respectively (**Fig.2A-D**). FACS of endothelial *versus* non-endothelial cell fractions confirmed EC enrichment for *prdm16* at 24 hours post-fertilization (hpf), which was subsequently lost at later developmental stages when the neuronal expression of *prdm16* becomes more prominent (**Fig.S1D**). Nonetheless, prdm16 still marked the DA in 6-day-old zebrafish embryos (**Fig.2D**). Finally, additional WISH experiments revealed that, parallel to our findings in mice, *prdm16* was present in the arterial endothelium at the earliest stages of vessel formation (**Fig.S2**), suggesting a key role for this TF in early arterial development.

**Figure 2.**
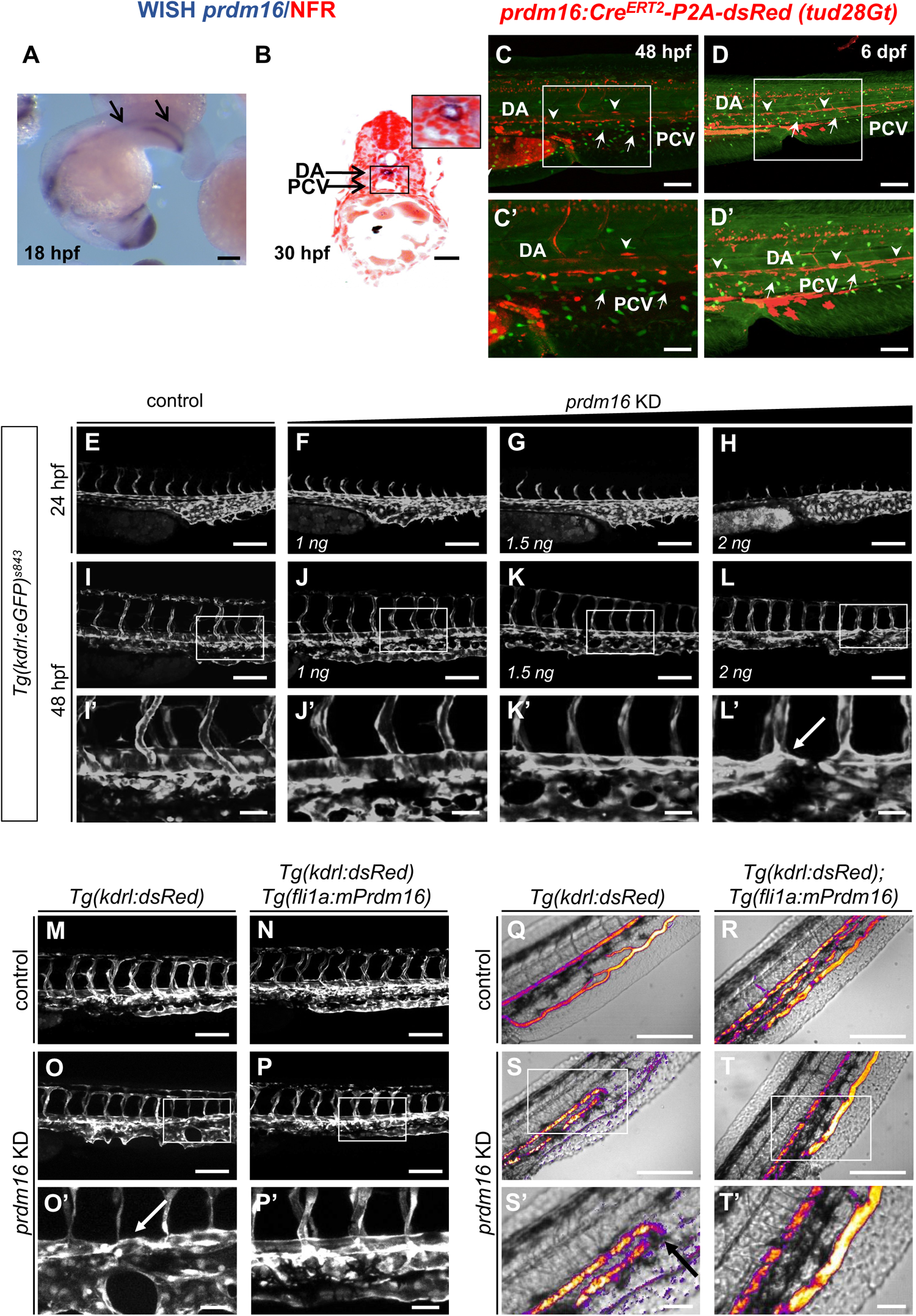
Endothelial *prdm16* is required for proper vascular development in zebrafish. **(A)** WISH for *prdm16* in an 18 hpf zebrafish embryo. Black arrows indicate *prdm16* expression in the DA. **(B)** Cross-section of a 30 hpf zebrafish embryo after WISH for *prdm16* counterstained with NFR, indicating *prdm16* expression in the DA, but not the posterior cardinal vein (PCV). **(C,D)** Confocal images of *tud28Gt* zebrafish embryos, reporting prdm16 activity (red), illustrating the presence of *prdm16* in the DA (white arrowheads), but not the PCV (white arrows) of zebrafish embryos at 48 hpf (C,C’) and 6 dpf (D,D’). **(E-L)** Confocal images of *Tg(kdrl:eGFP)^s843^* control embryos pr embryos with increasing KD of *prdm16* at 24 (E-H) and 48 hpf (I-L’), indicating angiogenic and aortic defects in *prdm16* KD but not control embryos during early vascular development. **(M-P)** Confocal images of *Tg(kdrl:dsRed)* and *Tg((kdrl:dsRed;fli1a:mPrdm16))* zebrafish embryos at 48 hpf, demonstrating that EC-specific overexpression of mPrdm16 rescues the vascular defects observed in *prdm16* KD *Tg(kdrl:dsRed)* embryos (white arrow). **(Q-T)** Angiograms generated by DIS of control and *prdm16* KD embryos with or without EC-specific overexpression of mPrdm16, illustrating the presence of an AVM (AV shunt) in a *prdm16* KD *Tg(kdrl:dsRed)* zebrafish embryos (S, black arrow in S’), which is no longer seen when mPrdm16 is simultaneously overexpressed in the endothelium (T,T’). Note the absence of vascular abnormalities in control *Tg(kdrl:dsRed)* and *Tg((kdrl:dsRed;fli1a:mPrdm16))* embryos (Q,R). Scale bars: 200μm in A; 100μm in B,E-L’,M-P,Q-T’; 50μm in C,D; 25μm in O’,P’. Panels labeled with*’* represent insets.

To determine if Prdm16’s function during vascular development extended beyond its reported role in angiogenesis^29,32^ we performed morpholino (Mo)-mediated *prdm16* KD in *Tg(kdrl:eGFP)^s843^*zebrafish embryos, in which all ECs are labeled by the eGFP fluorophore. This revealed dose-dependent vascular defects hallmarked by abnormal arterial ISV development and aortic hypoplasia at 24 hpf compared to control embryos injected with a scrambled Mo (**Fig.2E-H**). These vascular malformations persisted throughout development as we – in line with previous findings,^32^ observed reduced ISV length in *prdm16* morphants at 48 hpf (**Fig.S3A**). We additionally noted reduced aortic diameters, aortic coarctation (**Fig.2I-L**), and abnormal blood flow patterns in *prdm16* KD but not control embryos. Pairwise digital image subtraction (DIS) to visualize these aberrant flow patterns indicated that *prdm16* KD resulted in AVMs, i. e. abnormal connections between the axial vessels resulting in premature venous return of blood (**MoviesS1,S2**). The AVMs in *prdm16* KD embryos were not secondary to altered hemodynamics or cardiac performance, as they were still observed in *Tg(cmlc2:PRDM16^WT^)* zebrafish which overexpress human PRDM16 in cardiomyocytes and better preserve cardiac function upon *prdm16* KD (**Fig.S3B-D**). We then generated zebrafish specifically overexpressing mouse Prdm16 throughout their endothelium and crossed these *Tg(fli1a:mPrdm16)* zebrafish (which also carried a cmlc2:eGFP reporter to indicate presence of the transgene) with *Tg(kdrl:dsRed)* transgenics to analyze vascular development under baseline conditions and after *prdm16* KD. *Tg(fli1a:mPrdm16;kdrl:dsRed)* zebrafish did not display fusions between the DA and the PCV or other vascular abnormalities, indicating that endothelial overexpression of Prdm16 by itself does not interfere with proper vascular development. However, EC-specific overexpression of mPrdm16 did rescue *prdm16* KD-induced aortic defects (**Fig.2M-P**) and AVM occurrence (**Fig.2Q-T; Fig.S3E**). Together, these results illustrate an important, cell-autonomous role for endothelial Prdm16 during arterial development.

### Prdm16 activates Notch expression and signaling in human ECs and zebrafish

To unravel the molecular mechanisms behind these *in vivo* phenotypes, we performed whole genome transcriptomics on human endothelial progenitor cells (blood outgrowth endothelial cells; BOECs) overexpressing mouse Prdm16 (mPrdm16+) or the inert fluorophore Cherry (control). This approach was chosen as others and we showed that Prdm16 expression is very rapidly (within 48 hours) silenced in cultured arterial ECs,^41,33^ preventing us from performing siRNA studies; and because Prdm16 is known to regulate binary cell fate decisions in progenitor cells.^17,18,34^ We detected 441 differentially expressed genes (DEGs, FDR ≤ 0.05 and logFC ≥ |1|) between mPrdm16+ and control BOECs of which 284 and 157 were up- or downregulated, respectively (**TableS3**). In line with its expression pattern and the arterial defects observed in *prdm16* KD zebrafish embryos, gene ontology analysis revealed that Prdm16 controlled key processes such as ‘circulatory system development’ and ‘vascular development’, while more arterial-specific terms such as ‘artery morphogenesis’, and ‘artery development’ represented DEGs more robustly upregulated by Prdm16 (logFC ≥ |2|;**TableS4**).

Consistent with its known role as a negative regulator of TGFβ signaling,^39,42,43^ KEGG pathway analysis indicated that Prdm16 controlled TGFβ signaling in BOECs. Additionally, Prdm16 also modulated NOTCH signaling (**Fig.S4A; TableS5**), which is essential for arterial lineage specification and arterial development, as perturbations in Notch signaling cause aberrant arterial development and AVMs.^3,4,7,10,44–46^ Specifically, we found that Prdm16 predominantly induced the expression of NOTCH pathway components with a known arterial-restricted expression profile and implicated in arterial EC lineage specification and development, including *DLL4, HEY1, HEY2* and *Ephrin-B2 (EFNB2)* (**Fig.3A**). *DLL4, HEY1/2* and *EFNB2* mRNA levels were also significantly increased in mPrdm16+ human umbilical vein ECs (HUVECs) compared to control cells (**Fig.3B**). Enhanced expression of *HEY1/2* corresponded with increased *Hey1* and *Hey2* promoter activity in HUVECs (**Fig.S4B,C**), and with considerably higher levels of canonical NOTCH activity in both HUVECs (**Fig.3C**) and BOECs (**Fig.S4D**). We then investigated if NICD release was necessary to induce the NOTCH pathway downstream Prdm16 signaling, by incubating mPrdm16+ and control HUVECs/BOECs with N-[N-(3,5-Difluorophenacetyl)-L-alanyl]-S-phenylglycine t-butyl ester (DAPT) or vehicle (DMSO). DAPT attenuated the enhanced canonical NOTCH activity observed in mPrdm16+ HUVECs (**Fig.S4E**). Consistent with these findings, Prdm16 failed to induce *DLL4, HEY1, HEY2,* and *EFNB2* mRNA levels in HUVECs (**Fig.3D**) and BOECs (**Fig.S4F**) treated with DAPT, indicating that the effect of Prdm16 on NOTCH is dependent on the cleavage and subsequent release of NICD, which is blocked by DAPT. Prdm16 further induced the expression of the arterial-specific markers Gap junction protein alpha 5 (GJA5)^47,48^ and Dickkopf WNT signaling pathway inhibitor 2 (DKK2) in a NICD-dependent manner in both HUVECs and BOECs (**Fig.S4G,H**).

**Figure 3.**
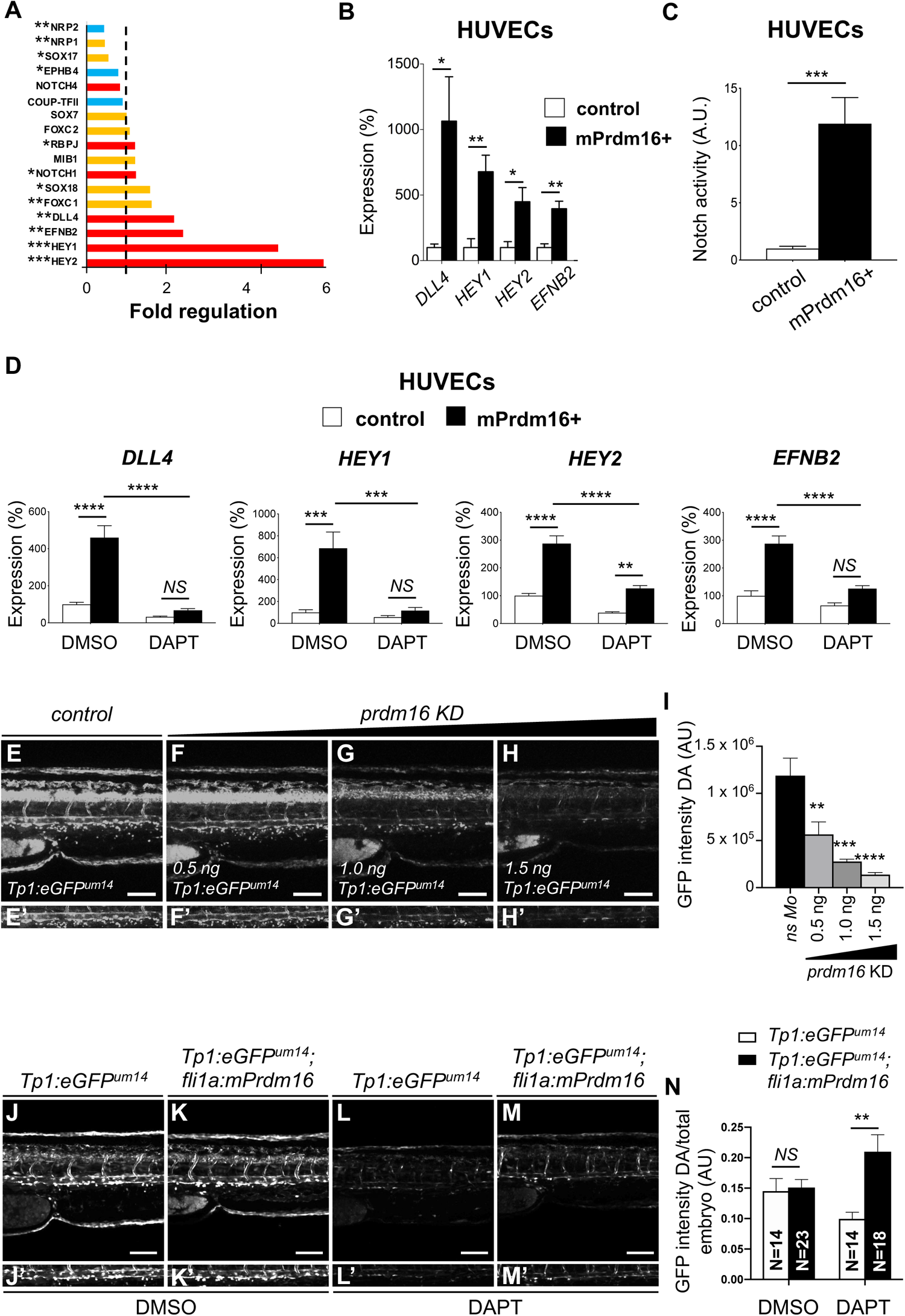
Prdm16 controls endothelial Notch signaling *in vitro* and *in vivo*. **(A)** Diagram showing the average fold regulation of select NOTCH pathway members known to be important for arterial development and of venous markers in mPrdm16+ versus control BOECs as determined by microarray analysis (N=3). Red and orange mark classical canonical Notch pathway members or Notch modifiers, respectively, while blue marks venous-specific genes. **(B)** qRT-PCR mRNA analysis (N=5) for *DLL4*, *HEY1*, *HEY2* and *EFNB2* in control (white bars) *versus* mPrdm16+ (black bars) HUVECs. (C) Canonical Notch activity in control (white bar) and mPrdm16+ (black bar) HUVECs transfected with Tp1:luciferase construct, illustrating the positive effect of Prdm16 on canonical Notch signaling. **(F)** mRNA levels of *DLL4*, *HEY1, HEY2* and *EFNB2* in control (white bars) and mPrdm16+ (black bars) HUVECs (N=6) treated with DMSO or the canonical Notch inhibitor DAPT, indicating that NICD is required for Prdm16-mediated stimulation of Notch signaling. **(E-I)** Confocal images of 48 hpf *Tg(Tp1:eGFP)^um14^* zebrafish control (E) embryos or embryos with increasing KD of *prdm16* (F-H) and corresponding quantification (I) of canonical notch activity (GFP intensity, N=5-6) in the dorsal aorta (DA; shown separately in E’-H’) of control (black bar) and *prdm16* KD *Tg(Tp1:eGFP)^um14^* embryos (gray bars). **(J-N)** Confocal images of 48 hpf *Tg(Tp1:eGFP)^um14^* (J,L) and *Tg((Tp1:eGFP)^um14^;fli1a:mPrdm16)* (K,M) embryos treated with DMSO (J,K) or DAPT (L,M) and corresponding ratiometric quantification (N) of canonical notch activity (GFP intensity, N=14-23) in the DA (shown separately in J’-M’) relative to the total embryo of *Tg(Tp1:eGFP)^um14^* (white bars) and *Tg((Tp1:eGFP)^um14^;fli1a:mPrdm16)* (black bars) embryos treated with DMSO or DAPT. Note that DAPT caused a similar reduction of GFP signal in non-endothelial tissues. \**P*<0.05; \*\**P*<0.01; \*\*\**P*<0.001; \*\*\*\**P*<0.0001; NS: not significant by paired Student’s *t*-test (B,C), by one-way (I) or 2-way ANOVA (D,N). Scale bars: 100μm.

To substantiate our *in vitro* findings and further analyze the impact of Prdm16 on *in vivo* Notch activity in the DA, we utilized the *Tg(Tp1:eGFP)^um14^* canonical Notch reporter zebrafish line.^49^ Endothelial Notch activity was also determined by Prdm16 *in vivo*, as evidenced by the dose-dependent loss of eGFP+ signal in the DA of *Tg(Tp1:eGFP)^um14^* zebrafish embryos injected with increasing concentrations of *prdm16* Mo (**Fig.3E-I**). We then crossed *Tg(fli1a:mPrdm16)* with *Tg(Tp1:eGFP)^um14^* zebrafish to analyze the effect of ectopic Prdm16 on vascular Notch signaling. Although EC-specific overexpression of mPrdm16 was insufficient to induce ectopic Notch activity in the PCV (**Fig.3J,K**), *Tg(fli1a:mPrdm16)* zebrafish embryos retained higher Notch activity levels in the DA when challenged with DAPT compared to their respective controls (**Fig.3J-N**). Importantly, this effect was specific to the vasculature, as DAPT-treated *Tg(Tp1:eGFP)^um14^;Tg(fli1a:mPrdm16)* embryos failed to preserve canonical Notch activity in non-endothelial tissues. Similarly, more residual canonical Notch activity was observed in the DA of *Tg(fli1a:mPrdm16) versus* control embryos when we knocked down *prdm16* (**Fig.S5A-E**), illustrating that Prdm16 defines endothelial Notch activity both *in vitro* and *in vivo*. Together, these data support an EC-autonomous role for Prdm16 during AV cell-fate specification and arterial development and indicate that this is largely mediated via its effect on canonical Notch activity in arterial ECs.

### Prdm16 suppresses the venous transcriptional program of ECs *in vitro* and *in vivo*

As Prdm16 is known to suppress alternate cell fates, we hypothesized that it favors arterial endothelial specification at the expense of venous endothelial lineage identity. Both mPrdm16+ HUVECs (**Fig.4A**) and mPrdm16+ BOECs (**Fig.S5F**) indeed showed reduced expression of several venous markers compared to their respective controls. Analogous to our observations for arterial-restricted genes, Prdm16 was unable to suppress Neuropilin-2 (*NRP2)* but not Endomucin (*EMCN*) mRNA levels in the presence of DAPT (**Fig.4B**), illustrating that Prdm16 at least partly suppresses venous lineage specification via modulating Notch activity.

**Figure 4.**
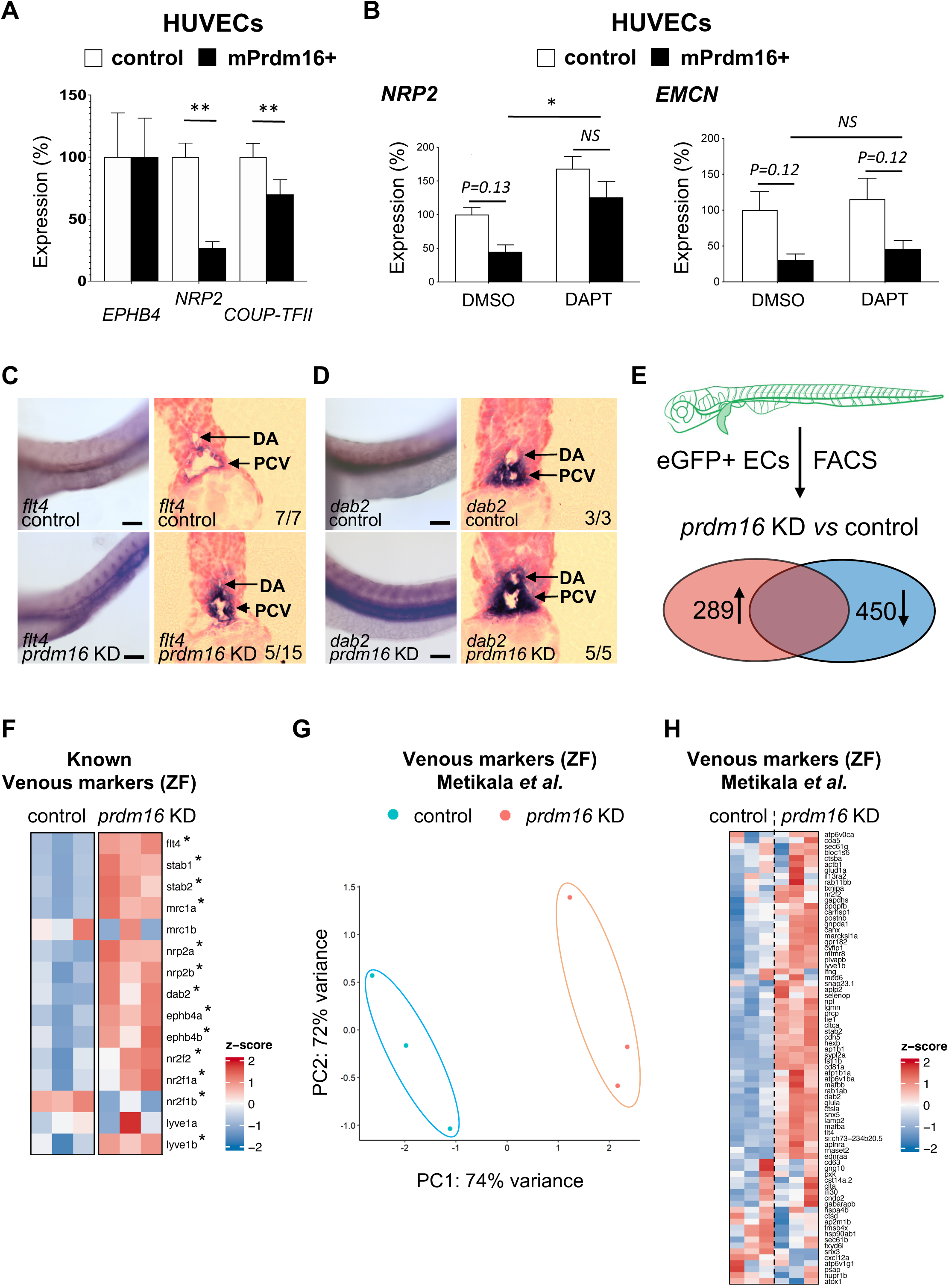
Prdm16 suppresses the venous transcriptional program in ECs. **(A)** qRT-PCR indicating reduced expression of the venous markers *NRP2* and *COUP-TFII* in mPrdm16+ *versus* control HUVECs. **(B)** mRNA levels of the venous markers *NRP2* and *EMCN* in control and mPrdm16+ HUVECs treated with vehicle (DMSO) or DAPT, demonstrating that Prdm16 inhibits venous lineage identity via both Notch-dependent and -independent mechanisms. **(C,D)** WISH for *flt4* (C) and *dab2* (D) in control and *prdm16* KD embryos at 24 hpf, indicating ectopic expression of typically venous-restricted *flt4* and *dab2* in *prdm16* KD embryos. **(E)** Schematic overview of the strategy for bulk RNA-sequencing from control and *prdm16* KD embryos, and the amount of DEGs. **(F)** Detailed analysis of the transcript levels of known venous markers in control and *prdm16* KD zebrafish embryos, illustrating the repressive effect of *prdm16* on these previously described venous markers. **(G)** Sample clustering of control and *prdm16* KD ECs based on the gene signature of venous ECs in the developing zebrafish as described by Metikala and colleagues. **(H)** Detailed analysis of endothelial expression of the venous endothelial-specific genes reported by Metikala *et al.* in control and *prdm16* KD embryos, demonstrating that *prdm16* suppresses the venous transcriptional program in ECs as indicated by the apparent upregulation of many genes that identified the venous EC population.

Prdm16 also repressed venous identity *in vivo,* as loss of arterial Notch activity in *prdm16* KD embryos (**Fig.3H**) was accompanied by ectopic expression of venous markers *flt4* and *dab2* in the DA at 28 hpf (**Fig.4C,D**). To better understand the effect of prdm16 on the entire endothelial trancriptome, we performed bulk RNA sequencing on FACS-isolated ECs from 28 hpf control or *prdm16* KD zebrafish (**Fig.S5G,H**), which identified 739 DEGs (FDR ≤ 0.05; log2FC ≥ |1|), of which 289 and 450 were up- or downregulated, respectively (**Fig.4E**). While canonical notch components implicated in arterial EC lineage specification (*dll4*, *hey1*, *hey2*, and *efnb2a*) were not differentially expressed in ECs from control and *prdm16* KD embryos, many previously described genes that specifically mark venous ECs in zebrafish (*flt4*, *stab1*, *stab2*, *mrc1a, nrp2a*, *nrp2b*, *nr2f2*, *nr2f1a*, and *dab2*) were significantly upregulated in *prdm16* KD embryos (**Fig.4F**). As their upregulation was modest and typically ranged between 1>log2FC >0.5, we reasoned that our transcriptomic dataset lacked single cell resolution to comprehensively analyze the effect of *prdm16* KD on the arterial and venous endothelial subpopulations specifically. We therefore took advantage of a previously published single cell RNA-seq dataset which identified an arterial and venous EC population in zebrafish at nearly the same developmental stage (30 hpf) as we performed our bulk RNAseq analysis,^50^ to analyze the transcriptional profile of control and *prdm16* KD ECs in more detail. Principal component analysis (PCA) based on the published arterial or venous gene signature separately clustered our control and *prdm16* KD samples (**Fig.4G**; **Fig.S5I**), emphasizing that prdm16 modulates the AV endothelial transcriptome beyond classical arterial and venous markers in zebrafish. ECs from *prdm16* KD zebrafish embryos adopted a more venous-like transcriptional program, as indicated by the apparent upregulation of many genes that identified the venous EC population (**Fig.4H**). Consistent with our previous results, arterial-specific genes were modestly affected with some being downregulated, while others displayed increased mRNA levels in ECs of *prdm16* KD embryos compared to controls (**Fig.S5J**). In summary, these data highlight that Prdm16’s primary *in vivo* function in ECs is to suppress the venous transcriptional program. This is likely also true in mammals, as others^29^ and we (**Fig.S5K**) observed ectopic EMCN in embryonic and postnatal arteries of EC-specific *Prdm16* knockout mice compared to their littermate controls.

### Prdm16 and Notch co-determine early vascular development in zebrafish

As Prdm16 interacts with Notch signaling in various non-vascular tissues^20,34,35^ and the vascular defects in *prdm16* KD embryos resembled those observed in notch-deficient zebrafish,^4,5,7,9^ we hypothesized that Prdm16 and Notch cooperatively orchestrate arterial development. To test this, we injected *Tg(kdrl:eGFP)^s843^* embryos with a subthreshold dose of the *prdm16* Mo that did not interfere with development, angiogenesis, or DA formation; while simultaneously lowering notch activity through inhibition of the dll4-nicd-rbpj_*k*_ signaling cascade at different levels (**Fig.5A,E,J,O**). Importantly, this approach circumvented the profound cardiac defects associated with higher doses of *prdm16* Mo or those associated with notch inhibition, allowing us to quantify AVMs in the absence of severe cardiac dysfunction (**Fig.S6A-C**).

**Figure 5.**
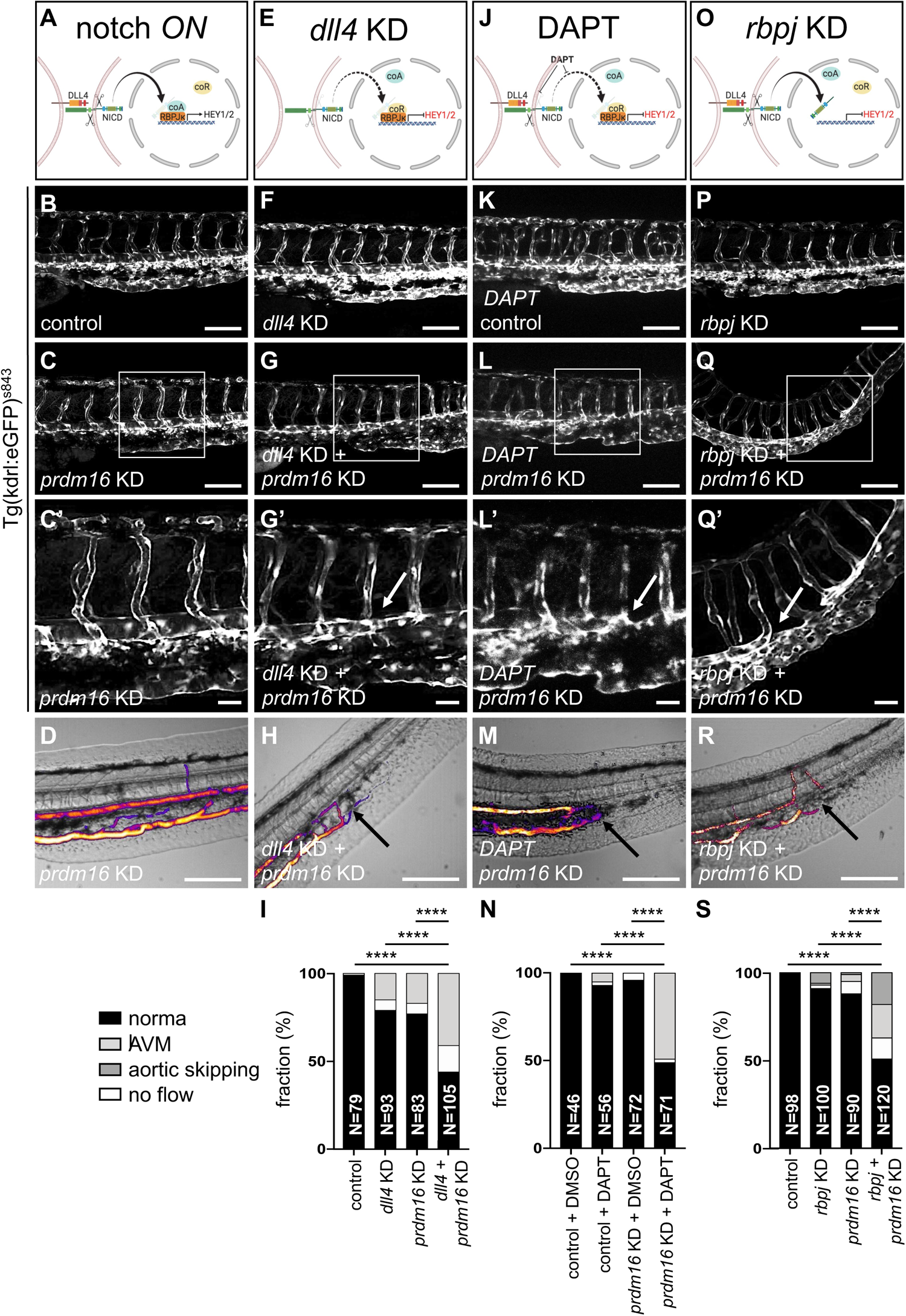
Genetic interaction between prdm16 and the notch signaling pathway during zebrafish vascular development. **(A)** Schematic overview of normal canonical Notch signaling. **(B-C’)** Confocal images of 48 hpf *Tg(kdrl:eGFP)^s843^* control (B) or subthreshold prdm16 KD (C,C’) embryos when notch is ON. **(D)** Angiogram of a 48 hpf *Tg(kdrl:eGFP)^s843^* embryo injected with a suboptimal *prdm16* Mo dose. **(E)** Schematic overview of canonical Notch signaling in the absence of the Notch ligand DLL4. **(F-G′)** Confocal images of 48 hpf *Tg(kdrl:eGFP)^s843^* embryos with subthreshold *dll4* KD alone (F) or KD of both *dll4* and *prdm16* (G,G’), demonstrating aortic defects in compound *dll4*/*prdm16* (white arrow in *g’*), but not single *dll4* or *prdm16* morphants. **(H)** Angiogram of *dll4*/*prdm16* KD embryos indicating the presence of an AVM (black arrow). **(I)** Quantification of the vascular defects in *dll4*/*prdm16* KD and control embryos. (**J**) Schematic overview of canonical Notch signaling in the presence of DAPT. **(K-L’)** Confocal images of 48 hpf *Tg(kdrl:eGFP)^s843^* control (K) or *prdm16* KD (I,I’) embryos treated with DAPT, indicating aortic defects (white arrow in I’) in *prdm16* KD embryos in the presence of DAPT. **(M)** Angiogram of DAPT-treated embryos with partial *prdm16* KD highlighting an AVM (black arrow). **(N)** Quantification of the vascular defects in DAPT or DMSO-treated control or partial *prdm16* KD embryos. **(O)** Schematic overview of canonical Notch signaling in the absence of RBPJ_*k*_. **(P-Q’)** Confocal images of *Tg(kdrl:eGFP)^s843^* embryos with incomplete *rbpj* KD (P) or subthreshold KD of both *rbpj* and *prdm16* (Q,Q’), the latter resulting in ‘aortic skipping’ and aortic coarctation (white arrow in Q’). **(R)** Angiogram of *rbpj*/*prdm16* double KD embryos illustrating the presence of an AV shunt (black arrow). **(S)** Quantification of the vascular defects in *rbpj*/*prdm16* double KD *vs.* control embryos. Panels labeled with*’* represent insets. \*\*\*\**P*<0.0001 by Chi square test. Scale bars: 100μm in B-D,F-H,K-M,P-R; 25μm in C’,G’,I’,Q’.

First, we combined subthreshold KD of both *prdm16* and *dll4.* While KD of *dll4* (or *prdm16*) alone did not result in overt vascular defects (**Fig.5B-D,F**), > 40% of compound *prdm16*/*dll4* morphants displayed aortic coarctation of the DA or AVMs (**Fig.5G-I**; **MoviesS4-6**). Similar results were obtained when notch was blocked with DAPT (25 µM) instead of *dll4* KD, as DAPT induced aortic coarctation and AVMs in ∼50% of embryos with partial *prdm16* KD, while DMSO-treated embryos or DAPT-treated controls remained unaffected (**Fig.5K-N**; **MoviesS7,8**). This synergistic effect between prdm16 and nicd release was further confirmed by the use of a second *prdm16* Mo (‘KD2’) (**Fig.S7A-F**) and another γ-secretase inhibitor (DBZ; **Fig.S7G-L**). Accordingly, the DA of *prdm16* KD embryos dosed with DAPT was completely devoid of arterial Notch activity at 24 and 48 hpf, while residual Notch activity could be detected in embryos challenged with *prdm16* KD or DAPT alone (**Fig.S7M-T)**, suggesting that complete loss of Notch is required to trigger AVMs. Finally, we knocked down *rbpjκ* using a previously validated *rbpja/b* Mo^51^ as in the absence of RBPJκ, ECs fail to induce *Hey1/2* promoter activity and hence, correct arterial differentiation (**Fig.5O**). AVMs were not observed upon partial KD of *rbpj* in zebrafish embryos (**Fig.5P,S**; **MovieS9**), but combined KD of *prdm16* and *rbpj* phenocopied the vascular deformations observed in *prdm16*/*dll4* KD and DAPT-treated *prdm16* KD embryos (**Fig.5Q-S**). Additionally, *prdm16*/*rbpj* morphants occasionally exhibited partial loss of the DA, with blood flow bypassing the missing arterial segment through sequential ISVs (‘aortic skipping’; **Fig.5R,S**; **MovieS10**). Altogether, these *in vivo* experiments strongly suggest that *prdm16* and *notch* work together to establish proper arteriovenous lineage identity and prevent AVMs.

### Impaired endothelial Prdm16 signaling facilitates DAPT-induced AVMs in zebrafish

We then adopted an acute CRISPR knockout (KO) system able to generate near complete null phenotypes in F0 zebrafish,^52^ which successfully targeted the entire *prdm16* locus (**Fig.S8A**). We found that >40% (16/36) of 48 hpf-old DAPT-treated *prdm16* CRISPant embryos displayed arterial defects reminiscent of those observed in our KD studies, while AVMs were not observed in DAPT-treated control or DMSO-treated *prdm16* CRISPants (**Fig.6A-D; Fig.S8B**). Thus, both Mo- and CRISPR/Cas9-mediated interference with *prdm16* evoked AVMs in zebrafish embryos challenged with DAPT. We also noted that DAPT administration resulted in abnormal, hyperbranched ISVs in *prdm16* CRISPants but not controls, without affecting ISV development in control embryos (**Fig.6A-D; Fig.S8C,D**). Hence, loss of *prdm16* in the presence of a subthreshold dose of DAPT results in hypersprouting phenotypes reminiscent of those observed in various notch morphants and mutants or embryos treated with higher doses of DAPT.^53,54^ Together, these data highlight that prdm16 and notch work together to warrant proper vascular development in zebrafish. To investigate if this interaction is mediated by endothelial Prdm16, we generated zebrafish overexpressing the dominant-negative, pathogenic PRDM16^K702Q^ variant^21^ in ECs using the UAS-Gal4 driver system. As expected, we did not observe significant vascular defects in *Tg(kdrl:Gal4)* or *Tg(UAS:PRDM16^K702Q^)* control embryos treated with vehicle or a low dose of DAPT (**Fig.6E,F,I**). Likewise, *Tg(kdrl:Gal4;UAS:PRDM16^K702Q^)* embryos did not display overt vascular anomalies under baseline conditions (**Fig.6G,I**; **MovieS11**), most likely due to the presence of endogenous zebrafish prdm16. Strikingly however, >50% of DAPT-treated *Tg(kdrl:Gal4;UAS:PRDM16^K702Q^)* embryos displayed AVMs (**Fig.6H,I**; **MovieS12**), which were observed in the absence of notable differences in cardiac performance (heart rate, stroke volume and cardiac output) between *Tg(kdrl:Gal4;UAS:PRDM16^K702Q^)* and control embryos in the presence or absence of DAPT (**Fig.6J-L**). Hence, interference with endothelial-specific prdm16 signaling predisposed zebrafish embryos to DAPT-induced AVMs.

**Figure 6.**
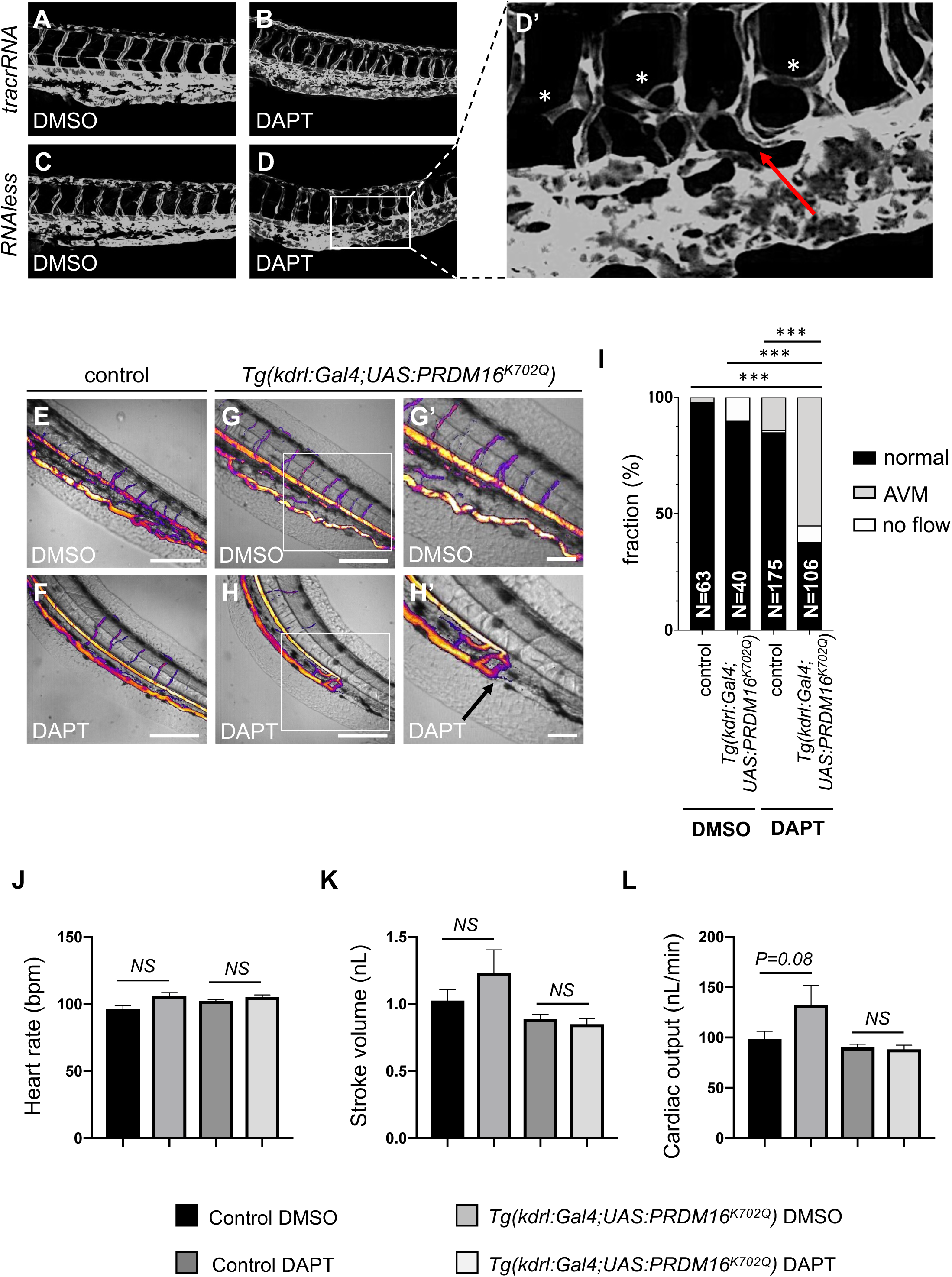
EC-specific overexpression of the dominant-negative PRDM16^K702Q^ variant results in AV shunting in the presence of DAPT. **(A-D’)** Confocal images of DMSO or DAPT-treated control (tracRNA) and *prdm16* CRISPant zebrafish *Tg(kdrl:eGFP)^s843^* embryos at 48 hpf, indicating the presence of aortic defects (red arrow) and vascular hypersprouting (white asterisks) in *prdm16* CRISPants challenged with DAPT, but not in DMSO-treated CRISpants or control embryos. **(E-H’)** Angiograms of 48 hpf control (E,F) and *Tg(kdrl:Gal4;UAS:PRDM16^K702Q^)* (G,G’,H,H’) zebrafish larvae treated with DMSO (E,G,G’) or DAPT (F,H,H’), indicating AVMs in DAPT-treated *Tg(kdrl:Gal4;UAS:PRDM16^K702Q^*) embryos (black arrow in H’). **(I)** Quantification of AVM occurrence in DMSO and DAPT-treated control and *Tg(kdrl:Gal4;UAS:PRDM16^K702Q^)* embryos. **(J-L)** Quantification of heart rate (J), stroke volume (K) and cardiac output (L) of control and *Tg(kdrl:Gal4;UAS:PRDM16^K702Q^*) embryos treated with DMSO or DAPT. \*\*\**P*<0.001; NS: not significant by Chi square (I) or 2-way ANOVA (J-L). Panels labeled with*’* represent insets. Scale bars: 100μm.

### Prdm16 amplifies the endothelial response downstream of NICD input

We next performed reactome pathway analysis of the genes upregulated in mPrdm16+ BOECs to gain more insights into the precise mechanisms via which Prdm16 regulates Notch. This analysis again emphasized the impact of Prdm16 on Notch signaling, but specifically highlighted its effect on signaling via de NOTCH1 receptor (**Fig.7A**). Surprisingly however, immunoblot analysis revealed similar levels of cleaved NOTCH1 ICD (N1ICD) in mPrdm16+ *versus* control HUVECs (**Fig.7B**), despite the inability of Prdm16 to boost Notch activity in DAPT-challenged HUVECs (**Fig.3D**) and BOECs (**Fig.S4E**). To further analyze the effect of Prdm16 on NICD-mediated signaling, we stimulated mPrdm16+ and control HUVECs/BOECs with recombinant (r)Dll4 or vehicle (0.1% BSA) and measured the effect on Notch via qRT-PCR. These experiments revealed that Prdm16 amplified the endothelial response to rDll4, as indicated by the synergistic effect of rDll4 and ectopic mPrdm16 on the mRNA levels of *DLL4*, *HEY1, HEY2* and *EFNB2* in HUVECs (**Fig.7C**) and BOECs (**Fig.S9A**). Additionally, mRNA levels of the venous markers *COUP-TFII*, *NRP2*, *EMCN* and *Fibronectin Leucine Rich Transmembrane Protein 2* (*FLRT2)* trended downwards in rDll4+mPrdm16 HUVECs (**Fig.S9B**) and BOECs (**Fig.S9C**) compared to either BSA-stimulated mPrdm16+ or Dll4-stimulated Cherry+ control cells. Concordantly, we observed higher DLL4 and lower FLRT2 protein levels in rDll4+mPrdm16+ compared to BSA-mPrdm16+ and rDll4+mPrdm16+ HUVECs despite similar levels of mPrdm16-overexpression (**Fig.S9D**). Altogether, these data suggested that the impact of Prdm16 on the endothelial transcriptome is determined by the absolute NICD levels. Together with the failure of Prdm16 to activate Notch in the presence of DAPT (**Fig.3D**; **Fig.S4E**), this prompted us to analyze a potential physical interaction between NICD and Prdm16. Co-immunoprecipitation (Co-IP) studies in HEK293 cells with the known Prdm16 binding partner CTBP2 as positive control, demonstrated that Prdm16 and N1ICD resided in the same protein complex (**Fig.7D**), likely explaining the NICD-dependency of Prdm16’s modulatory effect on the endothelial transcriptional program.

**Figure 7.**
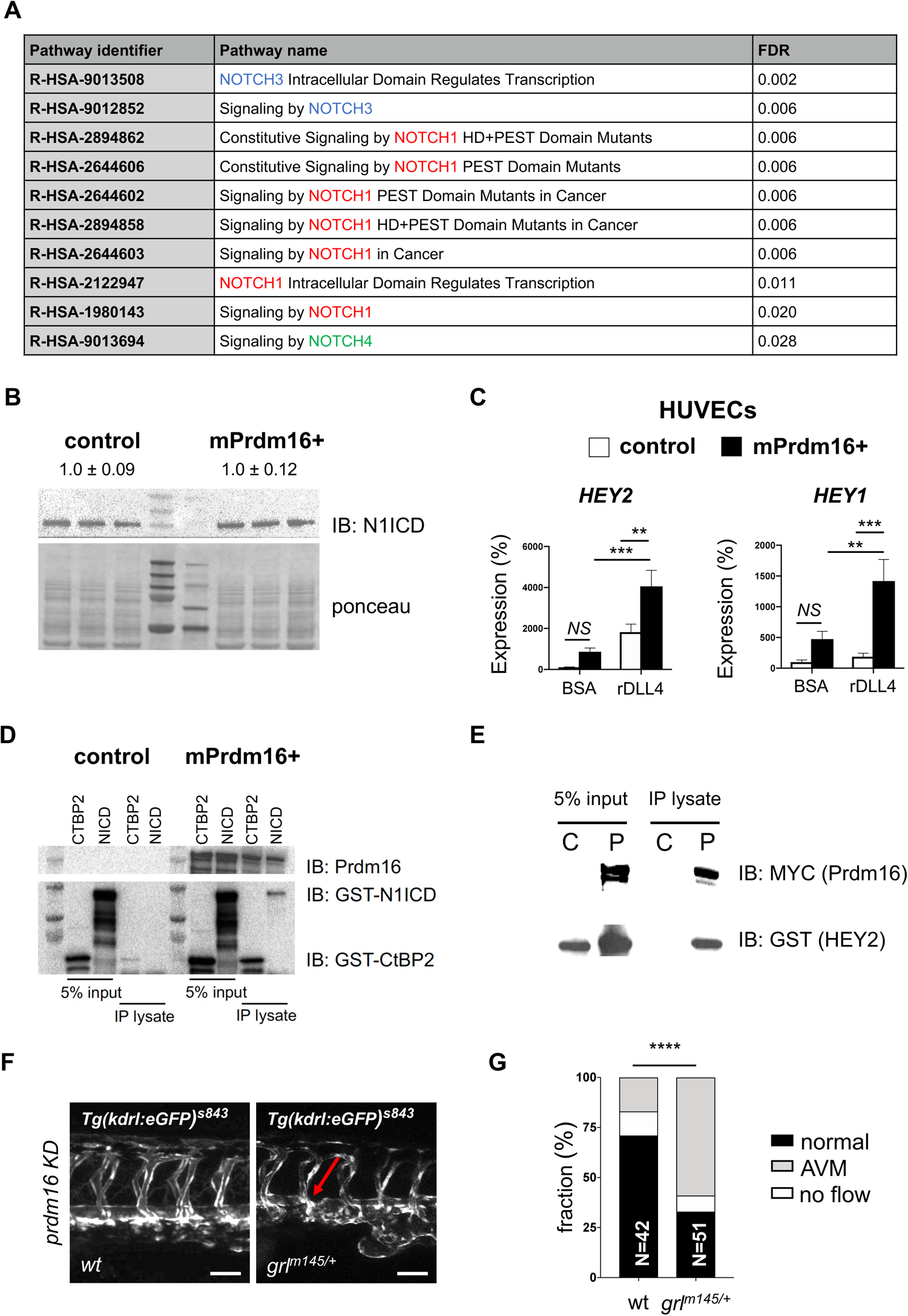
PRDM16 interacts with HEY2 to establish proper arterial development. **(A)** Reactome pathway analysis (https://reactome.org/PathwayBrowser/%23/) on genes upregulated in mPrdm16+ *versus* control BOECs, illustrating the involvement of NOTCH1 signaling downstream Prdm16-mediated biology. **(B)** Immunoblot for cleaved Notch1ICD (N1ICD) in control and Prdm16+ HUVECs, demonstrating that Prdm16 does not influence N1ICD levels. Ponceau red was used to normalize for protein loading. **(C)** qRT-PCR for *HEY1* and *HEY2* on control and mPrdm16+ HUVECs incubated on BSA or rDLL4-containing gelatin, demonstrating that Prdm16 amplifies the endothelial response to the DLL4 Notch ligand. **(D)** Co-immunoprecipitation of Myc-mPrdm16 and GST-N1ICD or GST-CtBP2 in control and mPrdm16+ HEK293 cells indicating that PRDM16 and N1ICD reside in the same protein complex. GST-CtBP2 was used as a positive control. **(E)** Co-immunoprecipitation of Myc-mPrdm16 and GST-HEY2 in control or mPrdm16+ HEK293 demonstrating that Prdm16 and HEY2 reside in the same protein complex. **(F)** Confocal imaging of wild-type (*wt*) and *grl^m145/+^* heterozygous embryos on the *Tg(kdrl:eGFP)^s843^* background with partial KD of *prdm16*. Incomplete KD of *prdm16* resulted in aortic coarctation (red arrow) in *grl^m145/+^*, but not *wt* embryos. **(G)** Quantification of AVMs in *wt* and *grl^m145/+^*embryos with partial *prdm16* KD. \*\**P*<0.01; \*\*\**P*<0.001; \*\*\*\**P*<0.0001; NS: not significant by 2-way ANOVA (C) or Chi square test (G). Scale bars: 50μm in F.

### Prdm16 functionally and physically interacts with HEY2

As Prdm16 typically drives lineage specification through its interaction with other DNA-binding TFs, we performed known motif enrichment analysis on genes up- or downregulated in mPrdm16+ versus control BOECs using HOMER software. The latter suggested the involvement of bHLH TFs in the repressive function of Prdm16, which was further confirmed by a similar analysis of our zebrafish dataset (**Fig.S10A**). As several of the consensus binding motifs of these bHLH TFs contained an E-box motif (CACGTG) - the known consensus binding sequence for HEY1/2, we speculated that Prdm16 might regulate AV lineage specification via its interaction with bHLH TFs, and more specifically HEY2. We therefore further investigated a potential interaction between PRDM16 and HEY TFs using ChIP-X Enrichment Analysis Version 3 (ChEA3), which integrates multiple resource types (co-expression, ChIP-seq, …) to generate a ranked list of TFs potentially controlling the expression of a specific set of DEGs.^55^ This analysis highlighted enrichment for both HEYL and HEY2 in the genes significantly upregulated in Prdm16+ *versus* control BOECs (**Fig.S10B**). Overall, these findings suggest a potential involvement of HEY2 in PRDM16-mediated AV lineage specification.

Additional co-IP studies in HEK293 cells demonstrated that PRDM16 indeed complexes with HEY2 (**Fig.7E**), in line with previous studies highlighting the functional and physical interaction between Prdm and bHLH TFs.^37,38^ To test the impact of this physical interaction on arterial development, we partially knocked-down *prdm16* or *hey2* alone or in combination. Although compound *prdm16*/*hey2* morphants displayed cardiac dysfunction, most embryos carried enough flow to reliably visualize and quantify AVMs (**Fig.S10C**). As expected, *prdm16*/*hey2* double morphants exhibited AV shunts and aortic coarctation defects while virtually all *hey2* or *prdm16* single morphants developed normally (**Fig.S10D-F; MoviesS13,S14**). To further investigate the interplay between Prdm16 and Hey2, we utilized the hey2 hypomorph zebrafish mutant, gridlock (*grl^m145^*).^9^ We crossed phenotypically normal *grl^m145/+^* fish with *Tg(kdrl:eGFP)^s843^*zebrafish and injected these embryos with a low dose of the *prdm16* Mo. Partial *prdm16* KD did not evoke vascular defects in wild-type *Tg(kdrl:eGFP)^s843^*embryos, but triggered aortic coarctation and AVMs in *Tg(kdrl:eGFP)^s843^;grl^m145/+^*embryos (**Fig.7F,G**; **MoviesS15,S16**). Titration of NOTCH activity by PRDM16 is therefore not confined to its potentiating effect on NICD-mediated signaling, but additionally involves a physical and functional interaction with the main arterializing TF downstream canonical NOTCH signaling, HEY2.

## Discussion

The molecular mechanisms that govern proper AV lineage specification to ensure the establishment of a defined network of arteries and veins are tightly controlled to prevent potentially life-threatening AVMs. Here, we demonstrate that the TF Prdm16 is exclusively expressed by arterial ECs throughout development, and that its vascular function extends beyond angiogenesis.^32^ We find that endothelial Prdm16 shapes the arterial vasculature and promotes arterial identity by suppressing the alternative venous cell fate. Our results indicate that PRDM16 amplifies canonical Notch signaling in ECs through its physical and functional interaction with both NICD and HEY2.

We first demonstrated that Prdm16 is a member of a select set of genes that hallmarks the arterial endothelial gene signature from zebrafish to human from the earliest developmental stages onwards. In zebrafish, KD of *prdm16* revealed previously undocumented vascular phenotypes, characterized by the occurrence of AVMs in the trunk. While others have previously knocked down *prdm16* in zebrafish, they did not observe AVMs, likely due to their localized nature, the presence of confounding cardiac defects resulting in reduced blood flow, and the need for DIS to visualize them. Importantly, these AVMs were still present in *prdm16* KD embryos when cardiac function was normalized, illustrating that they were not secondary to reduced cardiac performance or altered hemodynamics. Rather, our results suggest an EC-autonomous function for prdm16 during vascular development independent of its cardiac function, as EC-specific overexpression of mouse *Prdm16* protected against AVMs in *prdm16* morphants. Although targeted sequencing of isolated ECs from AVM lesions is needed to determine if somatic or inherited mutations in *PRDM16* truly contribute to the pathophysiology of human AVMs, these results indicate for the first time that Prdm16 might protect against AVMs.

*In vitro*, ectopic Prdm16 resulted in enhanced mRNA levels of known and newly identified arterial markers while simultaneously suppressing venous lineage identity. This was largely driven by boosting canonical NOTCH signaling in ECs, as Prdm16 was unable to fully arterialize BOECs and HUVECs in the presence of the canonical Notch inhibitor, DAPT. These observations indicate that Prdm16 belongs to a growing list of TFs that modulate endothelial Notch activity.^56,65–67^ Endothelial Prdm16 also determined arterial Notch levels *in vivo,* as KD of *prdm16* reduced canonical Notch signaling in the DA in control, but not in zebrafish embryos which overexpress mPrdm16 in their ECs. The latter were also protected against DAPT-induced loss of arterial Notch activity, further demonstrating the impact of endothelial Prdm16 on canonical Notch activity in the DA of the developing zebrafish. Consistent with a role for Prdm16 upstream of or in parallel with Notch, the vascular defects in *prdm16* morphants mirrored those observed in Notch-deficient zebrafish and mice.^3,4,7,10,44–46^ Concordantly, careful titration of *prdm16* and *notch* levels in zebrafish embryos revealed a synergistic effect between Prdm16 and Notch signaling, as combined subthreshold inhibition of *prdm16* and *notch* invariably led to arterial defects and AVMs in ∼50% of embryos in the absence of profound cardiac defects. AVMs were also evident in DAPT-but not DMSO-treated zebrafish embryos which specifically overexpressed the dominant negative PRDM16^K702Q^ variant in ECs, emphasizing the importance of endothelial Prdm16 for proper vascular development. Hence, our data identify a strong interaction between prdm16 and notch in ECs which parallels previous reports in non-vascular tissues.^20,34,35^ We further demonstrated that *prdm16* CRISPants phenocopied the increased susceptibility to DAPT-induced AVMs observed in *prdm16* morphants. The AVMs in DAPT-treated CRISPants occurred despite the lack of pronounced vascular defects under baseline (DMSO) conditions, most likely due to somatic mosaicism as a result of inefficient mutagenesis. Alternatively, this might be caused by redundancy between prdm16 and other prdm family members^57–59^ triggered by genetic compensation mechanisms readily observed in zebrafish mutants but not morphants.^60^ In support of the latter, others recently implicated Prdm3, the closest Prdm16 paralog, in maintaining arterial EC identity *in vitro*.^61^ Future experiments with defined *prdm16/prdm3* zebrafish mutants or EC-specific *Prdm3/Prdm16* KO mice could therefore reveal if both TFs co-determine arterial lineage specification and arterial development, in parallel with their well-documented redundant roles in adipose^58^ and craniofacial^57^ tissue.

Thompson *et al*. reported angiogenic defects in the postnatal retina of EC-specific *Prdm16* KO mice,^29^ while reduced angiogenesis was also previously observed in *prdm16* KD zebrafish.^32^ Although DIS is not suitable for analyzing angiogenesis in detail, we also noted impaired sprouting and reduced ISV lengths in 24 and 48 hpf-old *prdm16* morphant embryos. These angiogenic defects were however only observed with higher Mo concentrations when developmental delays became evident. Elucidating the precise role of *prdm16* during developmental angiogenesis therefore warrants further investigation, especially since we did not observe reduced sprouting in our *prdm16* CRISpants. Nonetheless, *prdm16* CRISPants challenged with DAPT displayed ISV hypersprouting reminiscent of those associated with higher doses of DAPT and reported in *notch* morphant and mutant zebrafish.^53,54^ Together with the occurrence of AVMs in both *prdm16* morphants and CRISPants, these results suggest a strong interaction between prdm16 and notch-mediated signaling during both angiogenesis and AV lineage specification.

We further observed that the reduced canonical Notch activity in *prdm16* morphants coincided with ectopic arterial expression of known venous markers, and a shift of the global endothelial transcriptome towards a more venous-like phenotype. These data highlight that Prdm16’s primary *in vivo* function is to suppress the venous transcriptional program, in line with its role as a repressor of alternative cell fates in cardiac and adipose tissue.^22,58^ Loss of *prdm16* did however not significantly lower the endothelial expression of classical notch pathway components associated with arterial lineage specification *in vivo* (e.g. *hey2*). Our findings indeed mirror those observed in *mib^ta52b^* mutant zebrafish embryos or embryos injected with a dominant-negative RBPJ_*k*_ construct, which both display preserved *hey2* expression within the developing (24 hpf) vasculature in the presence of ectopic mRNA levels of the venous marker *flt4* in their DA.^4^ *Vice versa*, notch1a-ICD overexpression represses *flt4* expression in the PCV, but fails to enhance arterial *hey2* mRNA levels, which is also not ectopically induced in the venous vasculature.^4^ It is possible that our bulk RNAseq strategy was not sensitive enough to detect relevant transcriptional changes in notch pathway genes which might have been revealed by single-cell RNAseq, due to the presence of both arterial and venous ECs in our samples. Additionally, compensatory mechanisms that safeguard sufficient expression of notch downstream targets when canonical notch is blocked, or spatiotemporal changes in notch activity and downstream target expression may exist. It has further been suggested that Hey2 can act upstream of Notch signaling^62^ and that *Hey2* expression can be regulated by Notch-independent pathways.^63,64^ Hence, Notch-independent mechanisms may control arterial *hey2* expression early on to warrant proper artery development, while Prdm16/Notch regulate Hey2 levels at later stages to maintain arterial identity. Additionally, these data may indicate that factors other than Notch and Prdm16 are needed to fully define the AV transcriptional profile of ECs during vascular development, which is further supported by the absence of eGFP+ signal in the venous vasculature of Notch reporter zebrafish embryos overexpressing mPrdm16 throughout their endothelium.

DAPT attenuated Notch induction in both mPrdm16+ BOECs and HUVECs, while this response was amplified in rDLL4-stimulated ECs. These results – together with our observations that combined loss of *prdm16* and *notch* triggered AVMs *in vivo –* illustrate that the absolute levels of NICD determine the molecular outcome of Prdm16-mediated lineage specification in ECs. mPrdm16+ HUVECs however did not display increased levels of cleaved NICD, nor did EC-specific mPrdm16 overexpression result in ectopic Notch activity in the PCV. These data indicate that Prdm16 co-determines arterial development and arterial lineage specification by amplifying Notch activity via mechanisms that complement the modulatory role of SOX^56,65,66^ and FOXC^67^ TFs on endothelial Notch activity. However, Prdm16 likely exerts its endothelial function via binding to other transcriptional mediators and TFs, as has been described in adipose,^17,68^ neuronal,^69,70^ and cardiac tissue.^20^ We found that Prdm16 enhanced canonical Notch activity in two separate and compatible ways. First, co-IP studies in HEK293 cells demonstrated direct binding of Prdm16 to NICD, congruent with our findings that *rbpj* KD or DAPT-treatment resulted in AVMs in otherwise phenotypically normal *prdm16* KD embryos. Binding of PRDM16 to NICD might augment the Notch output by enhancing NICD retention or transcriptional activity in the nucleus, without impacting NICD levels. Endo and colleagues demonstrated that the Drosophila homolog of Prdm16 *(hamlet)* modifies the chromatin at defined Notch target loci to determine the DNA occupancy of RBPJ_*k*_.^34^ As other chromatin modifying enzymes have been described as transcriptional co-repressors and co-activators of the NICD/RBPJ_*k*_ complex, it is tempting to speculate that Prdm16 functions as a co-activator. The Prdm16/NICD interplay could however be locus-dependent, as loss of *hamlet* in the sensory organ results in cell fate switches that are associated with both loss and gain of Notch activity.^34,71,72^ Additional studies are thus needed to further characterize PRDM16-NICD interactions under more physiological conditions, and indicate how the relative levels of endothelial NICD and PRDM16 modulate the molecular profile, function, and behavior of ECs.

Importantly, we demonstrated that PRDM16 also complexed with HEY2, the bHLH TF essential for arterial development and arterial EC specification in zebrafish,^4,7,8,9^ mice,^10^ and humans.^14^ The *in vivo* relevance of this interaction was confirmed by the occurrence of AVMs upon partial KD of *prdm16* in otherwise phenotypically normal *grl^m1^*^45^ heterozygous mutants, or in zebrafish embryos with combined subthreshold KD of *prdm16* and *hey2*. Additionally, the elevated *HEY2* mRNA levels in mPrdm16+ HUVECs/BOECs might be uniquely permissive for the physical and functional interaction between Prdm16 and Hey2 to drive these human ECs towards the arterial cell lineage, as both Prdm16 and Hey2 are typically near absent in cultured ECs.^33^ The PRDM16/HEY2 interaction described here echoes previously reported protein-protein complexes between PRDM and bHLH TFs in the ventral neural tube^36^ and during proper axon targeting.^37^ Future studies to fully define how PRDM16 modulates HEY2 function in vascular and non-vascular tissues could therefore shed further light on the complex interplay between PRDM and bHLH TFs.

In summary, this study identifies the functional interaction between Prdm16 and Notch in ECs, unravels the molecular mechanisms via which Prdm16 titrates arterial canonical Notch activity, and demonstrates that Prdm16-mediated signaling could represent a novel therapeutic target to treat AVMs.

