## Supplement for "Prdm16 amplifies Notch signaling and suppresses venous lineage specification to prevent arteriovenous malformations during vascular development"

#### Online Methods

##### EC isolation and propagation

***Isolation of human umbilical cord ECs and BOECs***: Umbilical cords were collected in phosphate-buffered saline (PBS) containing penicillin/streptomycin/amphotericin (PSA) and a gavage needle was inserted into both arteries. Cords were subsequently massaged and perfused with 20 mL ice-cold PBS to remove blood clots and check for perforations. After washing the cord, both sides were clamped off and 2.5 mL of a 0.2% collagenase type I solution was added to each umbilical artery through the gavage needles and incubated for 15 minutes at 37°C. Detached HUAECs were then flushed out with 20 mL PBS in a 50 mL falcon tube and filtered over a 40 μm cell strainer. HUVECs were then collected in a similar fashion and both HUAECs and HUVECs were incubated in EGM2-MV medium (CC-3202, Lonza) in 0.1% gelatin-coated flasks. Human BOECs were isolated as previously described. Briefly, peripheral blood was collected in heparin-containing syringes and diluted 1:1 with PBS supplemented with 1% PSA. Blood was then divided in 10 different 15 mL falcon tubes and carefully mixed with 2.5 mL histopaque (Ficoll Paque™ PLUS, GE Healthcare 17-440-02) and centrifuged for 30 minutes at 750 g after which the upper plasma layer was aspirated and discarded. The remaining buffy coats were pooled and collected in new 50 mL falcon tubes to which 45 mL of PBS supplemented with 1% PSA and 2% fetal bovine serum (FBS) was added. Cells were spun down, washed twice, resuspended in 18 mL EBM-2 medium (CC-3162, Lonza) and incubated in collagen type I-coated 6-well plates (657950, Greiner Bio-One). Cell medium was changed frequently and endothelial colonies typically appeared 11 to 28 days after plating. The use of human cells was approved by the Ethics Committee of University Hospitals Leuven (No B322200084019) and experiments were performed in accordance with the Committee's guidelines.

***Isolation of mouse arterial and venous ECs:*** Thoracic aorta and vena cava from *Tie2-GFP* mice[39] (expressing GFP in blood-vascular but not lymphatic ECs) were dissected out, pooled and placed in ice-cold MCDB131 medium (Gibco). Excess fat and contaminating tissue surrounding the vessels were removed and vessels were cut into small pieces prior to incubation in ice-cold PBS containing 2 mg/mL crude collagenase extract (Roche) at 37°C until fully dissociated. Digests were repeatedly shaken, filtered through a 100 μm nylon mesh (BD Biosciences) and spun down (600 g, 7 minutes). Arterial and venous single cell suspensions were then resuspended in PBS supplemented with 1% bovine serum albumin (BSA) and run through a FACS Aria™ device (Beckton Dickinson) to separate the endothelial ($GFP^+$) from the non-endothelial ($GFP^-$) cell fraction. EC and non-EC fractions were collected by FACS and directly collected in RNA lysis buffer. Purity was analyzed by quantitative reverse-transcription polymerase chain reaction (qRT-PCR) for endothelial, haematopoietic, and smooth muscle cell markers. Approximately 12 to 20 mice were used per isolation. Studies using mice were approved by the Ethical Committee of the animal facilities at KU Leuven (ECD 003/2012) and experiments were performed in accordance with the Committee's guidelines.

***Isolation of zebrafish ECs***: FACS sorting of zebrafish ECs was performed on single cell suspensions of 24 hpf, 48 hpf, 72 hpf and 6 dpf *Tg(kdr-eGFP)*$^{s843}$ embryos obtained by trypsinization (0.25% trypsine; Gibco) at 28°C. Vascular ($eGFP^+$) and non-vascular ($eGFP^-$) cell fractions were sorted with high purity in RNA extraction buffer supplemented with β-mercaptoethanol using a FACS Aria™ device.

***Propagation of ECs:*** HUVECs and BOECs were passaged when ~80% confluent in a 1:3 ratio. For all experiments, early (≤6) passage HUVECs or BOECs were used from at least 3 different donors.

**Lentiviral production and transduction of ECs**

***Lentivirus production:*** HEK293 cells were plated ($5x10^6$ cells/10 cm dish) and transfected the next day with the plasmid expressing the gene of interest and 2 helper plasmids (psPax2 and PMD2G) using Fugene transfection reagent. Briefly, 400 μL OPTIMEM (31985062, Gibco) was mixed with 1 μg PMD2G, 3 μg psPax2 and 4 μg plasmid of interest. Twenty-four μL of Fugene (E2311, Promega) was added and the mixture was incubated for 20 minutes at room temperature (RT) and gently applied to the cells. Medium was replaced the following day and lentiviral particle-containing supernatant was collected 36 hours later. Viruses were concentrated by centrifugation, semi-quantified and used directly to transduce cells or stored at -80°C until further use.

***Lentiviral transduction:*** Cells were plated at a density of 12,500 cells/$cm^2$ in a gelatin-coated 24-well plate the day prior to transduction. The next day, cells were incubated in fresh medium and transduced with pRRL2-mPrdm16-PGK-Cherry or pRRL2-empty-PGK-Cherry lentiviral particles 6 hours later. Medium was changed one day after transducing the cells and again at day 3 and RNA lysates were collected 6 days after transduction using TRIzol™ reagent (15596026, Invitrogen). For Western blot analysis, a similar approach scaled to a 10 cm-petri dish was used and cells were collected in Laemmli buffer supplemented with proteinase/phosphatase inhibitors. To study the effect ectopic Prdm16 upon simultaneous inhibition or activation of the canonical

Notch pathway, cells were either incubated with 50 (or 5) µM DAPT or vehicle (DMSO) or cultured on recombinant DLL4 (1µg/mL; R&D Biosystems) or BSA-containing gelatin, respectively.

**Luciferase Assays**

Cells were transduced as described above with pRRL2-empty-PGK-Cherry or pRRL2-mPrdm16-PGK-Cherry lentiviral particles, combined with a RBPJκ-luciferase reporter virus (Cignal Lenti RBP-Jk Reporter (luc) Kit: CLS-014L; SABiosciences; Qiagen) and a lentiviral Renilla control to normalize for transduction efficiency (Cignal™ Lenti Renilla Control (luc): CLS-RCL; SABiosciences; Qiagen). Medium was changed the following day and again at day 3 and cells were collected in 50 µL 1x luciferase lysis buffer 6 days after transduction. Firefly and Renilla luciferase activity were measured from 20 µL of lysate according to the manufacturers' protocol using the Dual-Luciferase® Reporter Assay System (E1910) from Promega on a Microplate Luminometer LB 96V (EG&G Berthold).

To score for *HEY1* and *HEY2* promoter activity, cells were co-transfected with pRRL2-mPrdm16-PGK-Cherry, pTK-renilla and pGL3-HEY1:luc or pGL3-HEY2:luc (kind gifts from Dr. Manfred Gessler, Würzburg, Germany) and promoter activity was measured 24 hours later. A pRRL2-empty-PGK-Cherry vector was used as control.

**EC transcriptional profiling**

***Microarray analysis of mouse arterial and venous ECs:*** Pooled RNA of ECs from 8-12 week-old mice was extracted (RNeasy minikit, Qiagen). The quality and purity of the RNA samples was analyzed by the VIB Nucleomics Core facility with a Bioanalyser 2100 and validated by PCR analysis prior to further proceedings. Five hundred pg of RNA from 5 selected isolations per tissue was amplified, labeled with biotin and hybridized on a mouse genome-wide microarray (Affymetrics Mo Gene1.0 ST array). The qualitative and statistical analysis of the microarray output was performed by the Nucleomics Core. In general, genes were considered differentially expressed between arterial and venous ECs if they displayed at least 2-fold enrichment in either of these two fractions ([log-value] ≥ 1) with a corrected *P*-value < 0.001. Such stringent approach was chosen to obtain 'workable' amounts of genes. To further define a conserved AV signature, we conducted a meta-analysis on this AV gene signature and a previously established list of human umbilical cord arterial and venous-specific genes[23] using a Fischer *t*-test on the combined data sets. To map the probe sets of both microarray platforms (*i.e.*, Human Genome U133 Plus 2.0 Array and the GeneChip Mouse Gene 1.0 ST Array), we first mapped the probe sets of the Human U133 Plus 2.0 array to the Mouse 430 2 array. This was done by mapping of homologous/orthologous probe sets with the package annotation Tools of BioConductor (annotation files *HG-U133_Plus_2.na32.annot.csv* and *Mouse430_2.na32.annot.csv* as downloaded from the Affymetrix website and HomoloGene database homologene.data build 66 as downloaded from NCBI). Secondly, we mapped the Mouse 430 2 probe sets to the probe sets on the Mouse Gene 1.0 ST Array, based on the mapping table as provided by Affymetrix. In case a probe set of one array maps to several probe sets of the other array, they are mentioned multiple times in the data table.

***Microarray analysis of mPrdm16- or Cherry-overexpressing BOECs:*** BOECs were transduced with pRRL2-empty-PGK-Cherry (control) or pRRL2-Prdm16-PGK-Cherry lentiviral particles as described above and RNA was collected 6 days after transduction. The quality and purity of the RNA samples was analyzed by the VIB Nucleomics Core facility with a Bioanalyser 2100 and validated by PCR analysis prior to further proceedings. Five hundred pg of RNA from 3 selected experiments was amplified, labeled with biotin and hybridized on a human genome-wide microarray (Affymetrix HG-U133 Plus 2.0 GeneChip Oligonucleotide Microarray). The qualitative and statistical analysis of the microarray output was performed by the Nucleomics Core. Genes were considered differentially expressed if they displayed at least 2-fold change compared to the control (Cherry) condition ([log-value] $\geq$ 1) with an FDR $\leq$ 0.05. Gene set enrichment and gene ontology analysis was performed through the UCSanDiego/Broad Institute GSEA software (https://www.gsea-msigdb.org/gsea/index.jsp). Differentially expressed genes were scanned for known motif enrichment using HOMER with default settings. Microarray data are accessible at the NCBI Gene Expression Omnibus website, accession number GSE131141.

**Expression analysis by staining or *in situ* hybridization**

***β-galactosidase staining:*** *Prdm16$^{Gt(OST67423)Lex}$* (also known as *Prdm16$^{Gt683Lex}$*) mice were generated by Lexicon Genetics and obtained via the Mouse Mutant Resource Repository Center (MMRRC). These mice carry an SA-Bgeo-PGK-Btk-SD stop cassette in the first intron of *Prdm16* of which the Bgeo cassette has been shown to faithfully reproduce the endogenous expression pattern of Prdm16. E9.5 or E10.5 embryos derived from *Prdm16$^{+/-}$* intercrosses, were processed as follows: embryos were briefly washed with PBS prior to fixation at 4°C with fixing solution for 20 minutes. Embryos were subsequently washed repeatedly with PBS and incubated with staining solution at 30°C overnight. Next day, embryos were washed three times with PBS and post-fixed with 4% PFA for 2 hours at RT. Tissues were washed multiple times, further processed for paraffin sectioning and nuclear fast red (NFR) staining. Briefly, slides were deparaffinated by subsequent washes with xylene, 100% EtOH, 96% EtOH, 70% EtOH and 50% EtOH and placed into the NFR solution for 5-10 minutes. Afterwards, slides were rinsed with MilliQ (MQ) water, dehydrated with ethanol (subsequently transferring the sections in 50% EtOH, 70% EtOH, 96% EtOH and three times in 100% EtOH), immersed twice in xylene and mounted with DPX. Embryos from *Prdm16*$^{+/+}$ mice were used as an internal negative control.

***Immunofluorescence staining for Prdm16:*** E14.5 or E17.5 FvB mouse embryos were dissected, rinsed in RNA Later (Ambion) and put in a 15 mL falcon tube to snap-freeze in liquid nitrogen. Unfixed cryopreserved embryos were sectioned (7 µm) on a cryostat (CM3000, Leica) prior to staining. Air-dried sections were incubated with 4% paraformaldehyde for 10 minutes at 4°C, washed with MQ water and an additional three times with TNT before being immersed in PBS-Tr (PBS with 0.1% Triton-X) for 30 minutes. Sections were blocked for 1 hour with Tris-NaCl-BMP buffer (TNB) containing 20% pre-immune donkey serum (PID; Sigma-Aldrich) and incubated overnight on a shaker with Sheep-anti-Prdm16 primary antibody (AF6295, R&D Systems) in 20% PID/TNB at 4°C. After washing three times with TNB-Tween buffer (TNT), sections were incubated with a secondary Donkey-anti-Sheep-Texas Red antibody (713-075-147, Jackson Immunoresearch Laboratories) and FITC-conjugated anti-αSMA (F3777, Sigma) in TNB for 2 hours at RT and once again washed three times with TNT before mounting with DAPI containing Prolong Gold. A similar procedure was followed to stain human umbilical cords and human arteries for PRDM16.

***Immunofluorescence staining for Endomucin:*** E14.5 embryos derived from *Prdm16*$^{+/-}$ intercrosses were collected in ice-cold PBS and fixated in 4% paraformaldehyde at 4°C overnight, washed 3X in PBS and embedded in paraffin before sectioning at 7µM. After deparaffination, sections were submerged in DAKO target retrieval solution pH6 (antigen retrieval) for 20min in a 100°C oven, followed by a slow cool down at room temperature for 30min. Sections were blocked for 1 hour with Tris-NaCl-BMP buffer (TNB) containing 20% pre-immune donkey serum (PID; Sigma-Aldrich) and incubated overnight on a shaker with Goat-anti-Endomucin primary antibody (AF4660, R&D Systems) in 20% PID/TNB at 4°C. After washing three times with TNB-Tween buffer (TNT), slides were incubated with secondary Donkey-anti-Goat Alexa-488 antibody (Invitrogen) 1/200 in TNB for 2 hours, and once again washed three times with TNT before mounting with DAPI containing Prolong Gold.

***Whole mount* in situ *hybridization:*** All buffers mentioned below are prepared with diethylpyrocarbonate (DEPC)-treated MQ water or DEPC-treated PBS. AB/TU zebrafish embryos were collected at appropriate developmental stages into a glass vial, filled with MQ water. After removing the water, embryos were fixed using 4% PFA for 1 hour on a shaker at RT. Embryos were washed once with 100% EtOH and fresh EtOH was added to the embryos for storage at -20°C. Embryos were first rehydrated by transferring them in 3 successive buffers with decreasing EtOH content. Ultimately, embryos were rinsed three times with PBS-T (PBS containing 0.1% Tween20) and permeabilized with 1x proteinase K (Roche) in PBS-T for 15' to open the cell membranes. The embryos were subsequently transferred into 0.1 M triethanolamine, 0.1 M triethanolamine (Sigma) with 1x acetic anhydride (Sigma) solution and 0.1 M triethanolamine with 2x acetic anhydride solution and washed two times with PBS-T before fixation with formaldehyde-PBS-T for 20 minutes. After four washes with PBS-T, the embryos were shortly incubated in settling buffer (50:50 mix of hybridization buffer and PBS-T) and briefly incubated in hybridization buffer for 10 minutes at 60°C. Afterwards, embryos were incubated overnight in hybridization solution containing the digoxygenin (DIG)-labeled probe of interest at 60°C. The next day, embryos were washed several times at RT with 2 x saline sodium citrate (SSC), 2 times with 0.2x SSC and two times with MAB (maleic acid buffer). Subsequently, 1 mL of MAB-BR (blocking reagent) was added for 30 minutes, prior to incubation for 4 hours with 1 mL of MAB-BR-serum with anti-DIG-AP antibody (1:2,000; Roche). Finally, embryos were incubated four times with MAB for 20 minutes at RT and subsequently overnight at 4°C. The next day, embryos were rinsed twice with alkaline phosphatase (AP) buffer before starting the color reaction by adding 1 mL BM purple AP substrate in the dark at RT. Depending on the intensity of the signal, the chromogenic reaction was stopped within hours to days by adding PBS-T. Finally, embryos were fixed with multiple aliquots of PFA fixative, bleached, rehydrated and stored in glycerol until further processing. Whole mount pictures were taken using a Zeiss Axio Imager Z1 microscope. Alternatively, embryos were paraffin-embedded, cross-sectioned (7 µm) on a microtome (Leica) and counterstained with NFR, as described above.

**RNA extraction, cDNA preparation and qRT-PCR**

To extract total RNA from cultured ECs, a TRIzol®-based protocol was used. First, 1 mL TRIzol® (Invitrogen) supplemented with 1% β-mercaptoethanol was added to the cells and stored at -80°C. After thawing, the samples were vortexed and 200 µL chloroform (Merck Chemicals) was added prior to incubation at RT for 10 minutes. Samples were then centrifuged at full speed for 15 minutes, after which the upper transparent layer was taken off and transferred into a new Eppendorf to which 500 µL isopropanol (Merck Chemicals), 200 µL 4 M LiCl (Merck Chemicals) and 1 µL glycogen (Invitrogen) were added. Samples were then again stored for at least 10 minutes at RT and after centrifugation at full speed for 10 minutes, the isopropanol layer was carefully discarded and 1 mL of cold 70% EtOH was added. Afterwards, the samples were centrifuged at top speed for 5 minutes and EtOH was carefully discarded. Again, 1 mL of cold 70% EtOH was added and samples were centrifuged at full speed for 10 minutes. After discarding the EtOH layer, samples were air-dried for at least 10 minutes and the remaining pellet was resuspended in RNase-free-water prior to storage at -80°C. For cDNA preparation from RNA samples, we used following protocol: 32 µL of RNA, to which we added 4 µL random hexamer primers and 4 µL 10 mM dNTP mix (Superscript III First Strand Kit) was denatured for 5 minutes at 65°C. Simultaneously, a solution of 8 µL 10x RT buffer, 16 µL $MgCl_2$ (25 mM), 8 µL DTT (0.1 M), 6 µL $H_2O$, 1 µL RNase OUT and 1 µL SSIII reverse transcriptase (Superscript III First Strand Kit) was prepared. Subsequently, 40 µL of the solution was added to each denatured RNA sample and the cDNA synthesis cycle was started (10 minutes at 25°C, 50 minutes at 50°C and 5 minutes at 85°C). The samples were cooled to 4°C and 1 µL RNaseH was added prior to incubation for 20 minutes at 37°C to break down residual RNA. For qRT-PCR, cDNA was subjected to PCR-based amplification and detected with SYBR green dye (Applied Biosystems). A standard threshold for DNA-based fluorescence detection (0.1) was set to standardize measurements and the $C_t$-value (defined by the number of cycles necessary to reach this threshold) was considered as the quantitative value of the gene expression level. Each measurement was done in triplicate. *GAPDH*, *$\beta$-actin* or *ef1$\alpha$* was used as housekeeping gene to standardise for the total amount of cDNA. Each PCR reaction was performed by adding 1 µL cDNA to 6 µL SYBR green, 4 µL MQ water and 1 µL primer mix (2.5 µM forward and reverse primer). The polymerase reaction was performed on a Real-Time PCR system (Step One Plus, Applied Biosystems): 2 minutes at 50°C, 10 minutes at 95°C and subsequently 40 rounds of amplification at 95°C for 50 seconds, each time followed by 4 seconds at 60°C. A list of primers is added in **Supplementary TableS6**.

**Western Blotting**

Proteins were extracted with Laemmli buffer (Promega) supplemented with protease inhibitors. For Western blotting, 10 to 40 µg of protein was loaded into each lane. Blot pictures were recorded with a Bio-Rad Chemidoc XRS+ molecular imager, equipped with Image Lab software (Bio-Rad laboratories). Antibodies targeting Prdm16 (AF6295; R&D Biosystems), Flrt2 (AF2877; R&D Biosystems), Notch1ICD (Val1744; Cell Signaling Technology), Dll4 (2589; Cell Signaling

Technology) and β-tubulin (loading control) were used. Alternatively, Ponceau staining was used as loading control. Band density analysis was performed using NIH Image software.

**Co-Immunoprecipitation**

In brief, $5.10^6$ HEK293 cells were plated one day prior to transfection. The medium was changed the following day and cells were transfected with different combinations of pRRL2-Myc-Prdm16-PGK-Cherry; pRRL2-Myc-empty-PGK-Cherry; pReceiverM04-GST-HEY2 and pReceiverM04-GST-GFP plasmids (GeneCopoeia). The medium was changed the next day and cells were collected in ice-cold PBS one day later, washed twice and resuspended in 1 mL ice-cold IP lysis buffer (50 mM Tris-HCl, pH 7.5; 150 mM NaCl; 0.5% Triton X-100; 10% glycerol; 1x complete proteinase inhibitor). Meanwhile, anti-Myc antibody was added to dynabeads which were blocked and washed beforehand and non-bound antibody was washed away. IP lysate was subsequently added to the dynabeads and incubated on a rotor for 4 hours at 4°C or stored at -80°C until used. Beads were washed 5 times and bound proteins were isolated by boiling the dynabeads resuspended in 1x Laemmli buffer for 10 minutes after which beads were magnetically removed.

***In vivo* zebrafish experiments**

***Zebrafish husbandry, genotyping and generation:*** Zebrafish husbandry and procedures performed in this study were approved by the Harvard Medical Area Standing Committee on Animals, the local Institutional Animal Care and Use Committee (2016N000276). Zebrafish embryos were incubated at 28.5°C in standard E3 medium.

***Zebrafish phenotyping:*** Confocal images were taken on a Fluoview FV1000 (Olympus) to visualize the vasculature or analyzed with ImageJ software to assess canonical Notch activity in the DA or whole embryo where indicated. Cardiac performance was measured as previously described[24]. Angiograms were generated by video microscopy of laterally positioned embryos using an Axioplan (Zeiss) upright microscope equipped with a FastCam PCI high-speed digital camera (Photron) and subsequent digital image subtraction in ImageJ using an in house-developed code. For select experiments, the total number of ISVs or ectopic sprouts visual through confocal images were counted after Z-stack projection with maximum intensity.

***Morpholino (Mo) knockdown and CRISPR/Cas9 KO:*** Antisense Mo oligomers (GeneTools LLC.) were designed to target the start codon or intron-exon boundaries of *prdm16*, *dll4*, *rbpj*, *hey2*. Lyophilized Mo's were resuspended in RNAse/DNAse free water to a final concentration of 1mM and further diluted to working dilutions prior to embryo injections. Sequences of Mo's used in this study are: *prdm16*$^{ATG}$ 5'-*TCTCGCCCTCGCCTTGGATCTCATC*-3; *prdm16*$^{splice}$ (or 'Mo2') 5'-*TCATTCGCTGTCTTCCCGTCTCCTG*-3; *dll4* 5'-*AAAGGTGAGCCAAGCTGCCATGCTT*-3'; *rbpj* 5'-*CAAACTTCCCTGTCACAACAGGCGC*-3; *hey2* 5'-*CGCGCAGGTACAGACACCAAAAACT*-3'; *nc* 5'-*GCCATGACATAAGATCCAAGCCAAC*-3'. One nl of working dilutions were injected into zebrafish embryos at the one-cell stage, at a concentration of 1 ng/nl (*dll4* KD and subthreshold *prdm16* KD), 0.25 ng/nl (*rbpj* KD and *hey2* KD). For the acute deletion of the entire *prdm16* locus, guide RNAs targeting the 5'-->3' sequences *CTAGCGCGCGCTGACTCGGG(TGG)* and *GGCGTCCAAGGAGTTCTGGG(AGG)* were purchased from IDT DNA and ribonucleotide complexes were made according to the manufacturer's instructions, prior to injections at the one-cell stage.

***DAPT and DBZ treatment:*** Zebrafish embryos were treated with vehicle (DMSO), 25 µM DAPT or 250 nM DBZ from 6 hpf onwards until imaging.

##### Statistical Analyses

Data were expressed as mean ± SEM and were analyzed by unpaired or paired (when applicable) Student's *t*-test when comparing two groups. When 3 or more categorical groups were present, 1-way (1 variable) or 2-way ANOVA (multiple variables) with post-hoc Tukey was used to estimate the significance of differences between groups. For *in vivo* phenotypic studies, the penetrance of the phenotype, the number of embryos exhibiting the different morphant phenotypes were counted and Chi-square analysis was used to determine whether this fraction differed between control and treatment groups. PRISM (PRISM8/9) software was used for statistical analyses and differences were considered significant when $P<0.05$. Statistics for microarray analysis are described in the corresponding microarray methodology sections above. A minimum of 3 biological replicates was performed for each experiment.


##### Acknowledgements

We thank David Traver (University of California, La Jolla, CA, USA) for the *prdm16* reporter zebrafish and Trista North (Boston Children's Hospital, Harvard Medical School, Boston, USA) for the Notch reporter zebrafish line. We would additionally like to thank P. Vandervoort, B. Hoekman and T. Vervoort for technical assistance. We also thank P. Carmeliet for the zebrafish WISH probes and the VIB Nucleomics Core for their help with the whole-genome transcriptomic data analysis.

**Author contributions**

Conceptualization, A.L., M.B.; Investigation, J.VW., S.C., C.M., A.L., and M.B; Resources, C.M., A.L. and M.B.; Writing – Original Draft, J.VW., C.M., A.L., and M.B.; Writing – Review & Editing, J.VW., S.C., A.KC., L.A., P.V., M.D., C.F., S.W., C.M., N.L., M.F., C.M., A.L., and M.B.; Visualization, J.VW., S.C., A.L. and M.B.; Formal Analysis, J.VW, S.C., A.KC., L.A., C.M., N.L., A.L., and M.B.; Supervision, M.F., C.M., A.L., and M.B.; Project Administration, M.B.; Funding Acquisition, J.VW., C.M., A.L., and M.B.

**Declaration of Interests**

The authors declare that they have no conflict of interest.

**Data and code availability**

The authors declare that the data supporting the findings of this study are available within the paper and its supplementary information. Bulk sequencing data (microarray and RNA-sequencing) reported in this manuscript will be submitted to the European Nucleotide Archive (ENA) and be made publicly available. Further information and requests for resources and reagents should be directed to and will be fulfilled by Manu Beerens.

**Figure S1. The transcription factor (TF) Prdm16 is specifically expressed in arterial endothelium. (A)** Strategy to identify Prdm16 as a potential regulator of arterial development and arterial endothelial specification. Using *ex vivo* transcriptional datasets from arterial and venous ECs from both human and mouse origin and a subsequent meta-analysis, we generated a list of 52 conserved (across species) arterial or venous-specific genes. This list contained 6 venous, and 3 arterial transcription factors (TFs), including *Prdm16*. **(B)** GTex expression dataset for *PRDM16* (https://www.gtexportal.org/home/) in human tissues, indicating very high expression levels for this TF in the aorta, the coronary and the tibial artery. **(C)** Publicly available single-cell RNA-sequencing datasets (Gtex portal) indicating that Prdm16 is highly expressed in blood endothelial cells (BECs) from different organs. **(D)** qRT-PCR analysis for *prdm16* mRNA levels in FACS-isolated endothelial (ECs) and non-EC fractions of *Tg(kdrl:eGFP)*$^{s843}$ zebrafish embryos at different developmental stages. Note that early during development, at 24 hpf, *prdm16* expression is markedly higher in ECs *versus* non-ECs. Data are normalized to a universal zebrafish (uniZF) RNA sample. dpf: days post-fertilization.

**Figure S2. Expression pattern of *prdm16* throughout zebrafish development revealed by WISH. (A-D')** Lateral (A-D) and dorsal (A'-D') view of zebrafish embryos after WISH for *prdm16* at different developmental stages, demonstrating expression in the hindbrain (HB), the telencephalon (Tel) and ventral diencephalon (VD) as early as 14 hpf. Expression in the dorsal aorta (DA) starts at 18 hpf, when the arterial ECs have just coalesced into one vessel and is more pronounced at 20 hpf, after which circulation starts. **(E-F')** Lateral (E,F) and dorsal (E',F') view of zebrafish embryos after WISH for *prdm16* at 22 (E,E') and 24 (F,F') hpf. Expression in the neuronal structures (HB,VD) and the DA remain evident at 22 and 24 hpf, while the intersomitic vessels (ISV) sprouting from the DA also express *prdm16*. Cells of the posterior cardinal vein, located just ventral of the DA, do not express *prdm16* mRNA. The non-vascular expression pattern of prdm16 was in concordance with previous reports. Tel: telencephalon; HB: hindbrain; VD: ventral diencephalon; PND: pronephric duct; DA: dorsal aorta; EP: epiphysis; OP: olfactory placode; PFM: pectoral fin muscle; ISV: intersomitic vessel. Scale bars: 200 μm.

**Figure S3. *Prdm16* KD-induced AVMs are independent of Prdm16's cardiac function.** (A) Confocal images of control and *prdm16* KD *Tg(kdrl:eGFP)*$^{s843}$ zebrafish embryos at 48 hpf (left panel) and quantification of ISV length (right panel), indicating perturbed angiogenesis in *prdm16* KD embryos. **(B)** Cardiac performance (nl/min) in *Tg(cmlc2:GFP)* and *Tg(cmlc2:PRDM16*$^{WT}$*)* zebrafish embryos after *prdm16* KD, indicating preserved cardiac function in *prdm16* KD *Tg(cmlc2:PRDM16*$^{WT}$*)* embryos. **(C)** DIS-generated angiograms of control and *prdm16* KD *Tg(cmlc2:PRDM16*$^{WT}$*)* embryos, illustrating the emergence of an AVM after *prdm16* KD in the absence of profound cardiac defects. **(D)** Quantification of the percentage of *Tg(cmlc2:PRDM16*$^{WT}$*)* which displayed an AVM after *prdm16* KD. **(E)** Quantification of the percentage of *Tg(kdrl:dsRed)* and *Tg(kdrl:dsRed;fli1a:mPrdm16)* zebrafish embryos with an AVM after *prdm16* KD. *$P<0.05$ by unpaired Student's *t*-test.; ***$P<0.001$ by Chi square test. Scale bars: 100 $\mu$m.

**Figure S4. Prdm16 induces canonical Notch activity in ECs. (A)** Heat map of Notch pathway components in control and mPrdm16+ BOECs, illustrating the positive effect of Prdm16 in the expression levels of Notch ligands, transcriptional co-activators, and Notch downstream transcription factors. **(B,C)** *Hey1* (B) and *Hey2* (C) promoter activity in control and mPrdm16+ HUVECs. **(D)** Graph demonstrating elevated canonical Notch activity in mPrdm16+ *versus* control BOECs. **(E)** TP1:luciferase measurements in control and mPrdm16+ HUVECs incubated with DAPT or vehicle (DMSO), demonstrating that Prdm16 induces canonical Notch activity in human ECs via NICD. **(F)** mRNA levels of *DLL4*, *HEY1*, *HEY2*, and *EFNB2* in control and mPrdm16+ BOECs treated with DMSO or DAPT, demonstrating that Prdm16 induces Notch signaling via its effect on the canonical, NICD-dependent Notch pathway. **(G,H)** mRNA levels of the arterial markers *GJA5* and *DKK2* in both control and mPrdm16+ HUVECs (G) and BOECs (H) in the absence or presence of DAPT, indicating that Prdm16 induces a broad arterial transcriptional program via canonical Notch signaling. *$P$<0.05; **$P$<0.01; ***$P$<0.001; ****$P$<0.0001 by paired Student's *t*-test (B,C,D) or 2-way ANOVA (E,F,G,H).

**Figure S5. Prdm16 represses the venous gene signature, while boosting canonical Notch signaling in zebrafish. (A-D)** Confocal images of 48 hpf *Tg(Tp1:eGFP)*$^{um14}$ (J,L) and *Tg((Tp1:eGFP)*$^{um14}$*;fli1a:mPrdm16)* injected with control or *prdm16* Mo, illustrating that EC-specific overexpression of mPrdm16 protects against loss of canonical Notch activity in the DA (emphasized in A'-D') after KD of endogenous *prdm16*. **(E)** Quantification of arterial *versus* total Notch activity in control *versus* *prdm16* KD *Tg(Tp1:eGFP)*$^{um14}$ and *Tg((Tp1:eGFP)*$^{um14}$*;fli1a:mPrdm16)* embryos. **(F)** mRNA levels of known venous markers in control *versus* mPrdm16+ BOECs, demonstrating that Prdm16 suppresses defined venous-specific markers in endothelial progenitor cells. **(G)** Volcano plot of DEGs of FACS-isolated ECs from control and *prdm16* KD *Tg(kdrl:eGFP)*$^{s843}$ zebrafish embryo. **(H)** Heat map of the DEGs of FACS-isolated ECs from control and *prdm16* KD *Tg(kdrl:eGFP)*$^{s843}$ zebrafish embryo. **(I)** Cluster analysis of our RNA samples of FACS-isolated ECs from control and *prdm16* KD embryos based on the genes that define the arterial EC population in the developing zebrafish embryo according to Metikala *et al*. **(J)** Heat map depicting the impact of *prdm16* KD on the arterial endothelial gene signature (Metikala *et al*.), demonstrating that *prdm16* both positively and negatively influences arterial-specific genes. **(K)** IF staining for Emcn in the jugular area of E14.5 WT and *Prdm16*$^{-/-}$ mouse embryos showing ectopic Emcn protein levels in the artery of the latter. $^{**}P<0.01$ by paired (F) or unpaired (K) Student's *t*-test (F). Scale bars: 100 μm.

**Figure S6. Cardiac function is preserved in zebrafish embryos with partial inhibition of both *prdm16* and *notch*. (A-C)** Heart rate (HR) in beats per minute (BPM) (A), stroke volume (SV) in nl (B) and cardiac output (CO) in nl/min (C) in *Tg(kdrl:eGFP)*$^{s843}$ zebrafish embryos at 48 hpf with subthreshold KD of *prdm16* and *dll4* (left panels), *prdm16* and *nicd* release (middle panels), or *prdm16* and *rbpj* (right panels).

**Figure S7. Canonical notch and prdm16 genetically interact during dorsal aorta (DA) formation in zebrafish. (A-D)** Confocal images of *Tg(kdrl:eGFP)*$^{s843}$ zebrafish injected with a subthreshold dose of either negative control (*nc;* A,B) or *prdm16* Mo (splice-site Mo – KD2; C-D') and treated with vehicle (DMSO; A,C) or the canonical Notch inhibitor DAPT (B,D,D'). While *prdm16* KD2 alone or treatment of control embryos with DAPT did not result in gross vascular defects, *prdm16* morphants treated with DAPT displayed aortic coarctation and occasionally lacked a segment of the DA (D, red arrow in D'). **(E)** DIS-generated angiogram of a DAPT-treated *prdm16* morphant demonstrating the presence of an AVM (black arrow). **(F)** Quantification of the vascular defects in control or *prdm16* KD2 embryos treated with DAPT or DMSO. **(G-J)** Confocal images of control (G,H) or *prdm16* KD (I,J,J') Tg*(kdrl:eGFP)*$^{s843}$ zebrafish embryos treated with DMSO (G,I) or the canonical Notch inhibitor DBZ (H,J,J'), illustrating the presence of aortic defects in DBZ-treated *prdm16* KD embryos (red arrow in J'), but not in their respective controls. **(K)** Angiogram of a DBZ-treated *prdm16* KD zebrafish embryo at 48 hpf, indicating the presence of a shunt between the DA and PCV, resulting in premature venous return of blood flow (black arrow). **(L)** Quantification of the occurrence of AVMs in control and *prdm16* KD embryos treated with vehicle (DMSO) or DBZ. **(M-T)** Confocal images of control and subthreshold prdm16 KD Tg*(Tp1:eGFP*$^{um14}$*)* zebrafish treated with DAPT or DMSO at 24 hpf (M-P) and 48 hpf (Q-T'). Partial KD of *prdm16* combined with DAPT-treatment resulted in complete loss of canonical Notch signaling at both 24 hpf (P) and 48 hpf (T) compared to controls in both non-endothelial and endothelial tissues (the latter demonstrated by loss of GFP signal in the DA; Q'-T'). $^{**}P<0.01$; $^{****}P<0.0001$ *versus* indicated condition by Chi-square test. Scale bars: 100 μm.

**Figure S8. DAPT-challenged *prdm16* CRISPants display vascular hypersprouting. (A)** Schematic overview and PCR-based validation of our acute *prdm16* KO strategy, which deletes the entire *prdm16* locus. **(B)** Quantification of the percentage of control or *prdm16* CRISPant embryos displaying vascular defects in the presence or absence of DAPT, emphasizing a synergistic effect between *prdm16* deletion and *notch* inhibition. **(C)** Quantification of the number of ISVs in DMSO- (white bars) or DAPT-treated (grey bars) control or *prdm16* CRISPant embryos, indicating a slight increase in the number of ISVs in DAPT-treated *prdm16* CRISPants compared to controls. **(D)** Quantification of the number of excess vascular branches sprouting from the ISVs in 48 hpf DMSO- or DAPT-treated control or *prdm16* CRISPant embryos, illustrating that *prdm16* CRISPants are sensitized to DAPT-induced vascular hypersprouting. **$P<0.01$; ***$P<0.001$; ****$P<0.0001$ *versus* indicated condition by Chi-square test (B) or 2-way-ANOVA (C,D).

**Figure S9. Prdm16 amplifies the Notch-ligand induced arteriovenous response in human ECs. (A)** mRNA levels of *DLL4*, *HEY1*, *HEY2*, and *EFNB2* in control (white bars) and mPrdm16+ (black bars) BOECs incubated on BSA- or rDLL4-containing gelatin as determined by qRT-PCR, demonstrating increased expression of Notch pathway components in rDLL4 stimulated mPrdm16+ BOECs compared to its relative controls. **(B)** qRT-PCR results indicating lower mRNA levels of known venous markers in rDLL4-stimulated mPrdm16+ HUVECs compared to non-stimulated mPrdm16+ or rDLL4-stimulated control HUVECs. **(C)** qRT-PCR results indicating lower mRNA levels of known venous markers in rDLL4-stimulated mPrdm16+ BOECs compared to non-stimulated mPrdm16+ or rDLL4-stimulated control BOECs. **(D)** Immunoblot for endogenous DLL4 and FLRT2 protein levels in control or mPrdm16+ HUVECs stimulated with rDLL4 or vehicle (BSA), demonstrating elevated DLL4 and reduced FLRT2 protein levels in rDLL4-stimulated mPrdm16+ HUVECs compared to its controls. **P*<0.05; ***P*<0.01; ****P*<0.001 as indicated by 2-way-ANOVA.

**Figure S10. Hey2 and Prdm16 cooperatively shape the arterial vasculature in zebrafish. (A)** Known motif enrichment analysis (HOMER software) on genes suppressed by Prdm16 in BOECs (upper panel) or zebrafish ECs (lower panel), highlighting the potential involvement of bHLH TFs (accentuated in yellow) in the repressive function of Prdm16 on gene transcription. **(B)** ChEA3 TF enrichment analysis (top 25) based on the genes upregulated in mPrdm16+ *versus* control BOECs, documenting enrichment for the E-box binding bHLH TFs HEYL and HEY2 (red arrows). **(C)** Cardiac performance as indicated by HR, SV, and CO in 48 hpf control zebrafish embryos or embryos with partial KD of *hey2*, *prdm16*, or both, indicating reduced cardiac performance in the latter. **(D)** Confocal images of control *Tg(kdrl:eGFP)*$^{s843}$ embryos and *Tg(kdrl:eGFP)*$^{s843}$ embryos with incomplete KD of *hey2*, *prdm16*, or both, illustrating aortic coarctation (white arrow) only in embryos with subthreshold inhibition of both *prdm16* and *hey2*. **(E)** DIS-generated angiograms from control embryos, or embryos with partial KD of *hey2*, *prdm16*, or *hey2* and *prdm16* at 48 hpf, pointing out an AVM in *hey2*/*prdm16* double morphants (black arrow), but not its respective controls. (F) Quantification of the vascular defects in control embryos and embryos with partial KD of *hey2*, *prdm16*, or both *hey2* and *prdm16*, demonstrating markedly more AVMs in embryos where both *hey2* and *prdm16* were inhibited. $^*P<0.05$; $^{**}P<0.01$; $^{***}P<0.001$ as indicated by 2-way-ANOVA. Scale bars: 50 μm.

**Fig.S1**

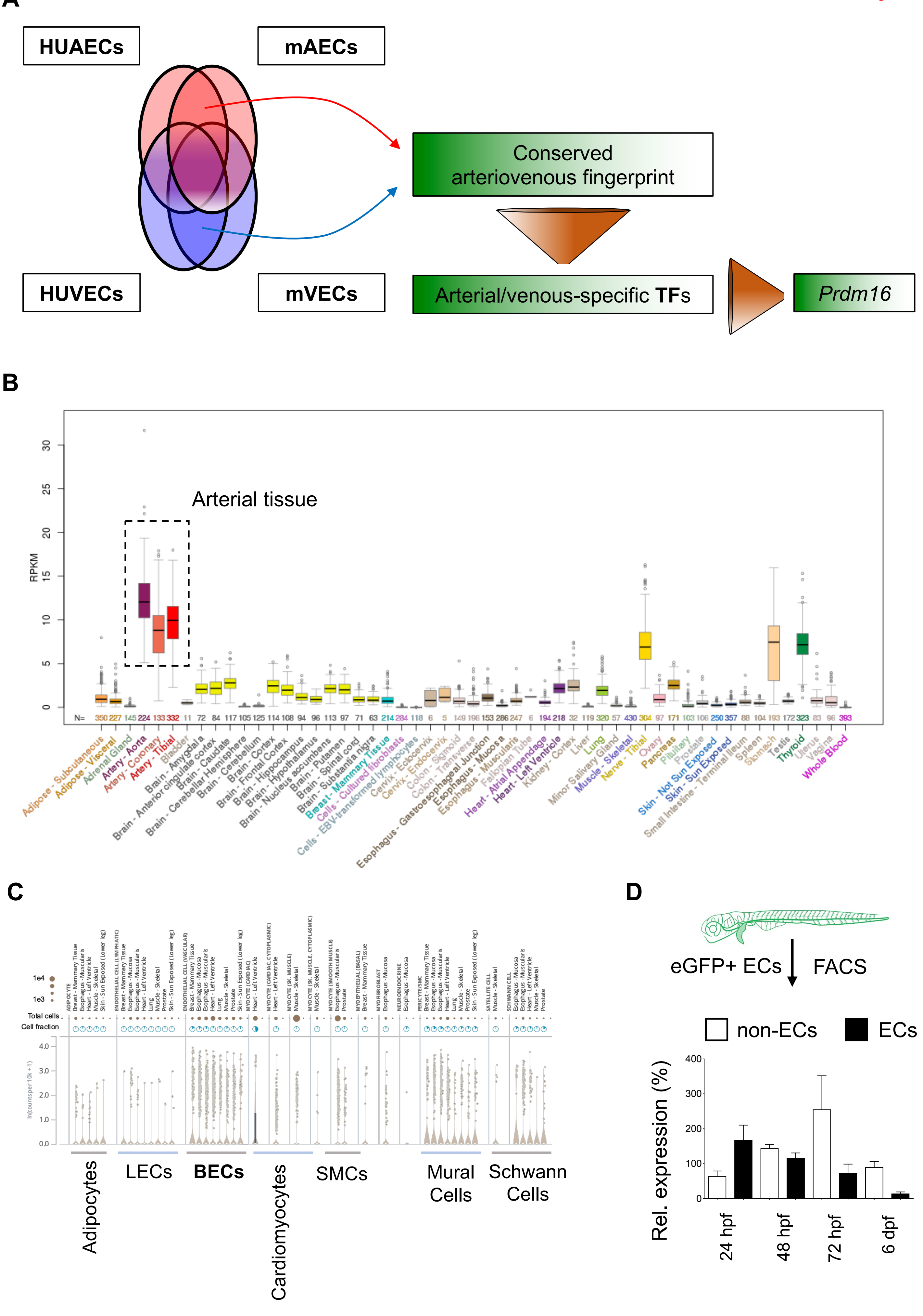
A
HUAECs
mAECs
HUVECs
mVECs
Conserved arteriovenous fingerprint
Arterial/venous-specific TFs
Prdm16
B
Arterial tissue
RPKM
N=
Adipose - Subcutaneous
Adipose - Visceral
Adrenal Gland
Artery - Aorta
Artery - Coronary
Artery - Tibial
Bladder
Brain - Amygdala
Brain - Anterior cingulate cortex
Brain - Caudate
Brain - Cerebellar Hemisphere
Brain - Cerebellum
Brain - Cortex
Brain - Frontal Cortex
Brain - Hippocampus
Brain - Hypothalamus
Brain - Nucleus accumbens
Brain - Putamen
Brain - Spinal cord
Brain - Substantia nigra
Breast - Mammary Tissue
Cells - Cultured fibroblasts
Cells - EBV-transformed lymphocytes
Cervix - Ectocervix
Cervix - Endocervix
Colon - Sigmoid
Colon - Transverse
Esophagus - Gastroesophageal Junction
Esophagus - Mucosa
Esophagus - Muscularis
Fallopian Tube
Heart - Atrial Appendage
Heart - Left Ventricle
Kidney - Cortex
Liver
Lung
Minor Salivary Gland
Muscle - Skeletal
Nerve - Tibial
Ovary
Pancreas
Pituitary
Prostate
Skin - Not Sun Exposed
Skin - Sun Exposed
Small Intestine - Terminal Ileum
Spleen
Stomach
Testis
Thyroid
Uterus
Vagina
Whole Blood
C
Total cells
Cell fraction
Adipocytes
LECs
BECs
Cardiomyocytes
SMCs
Mural Cells
Schwann Cells
D
eGFP+ ECs
FACS
non-ECs
ECs
Rel. expression (%)
24 hpf
48 hpf
72 hpf
6 dpf

**Fig.S2**

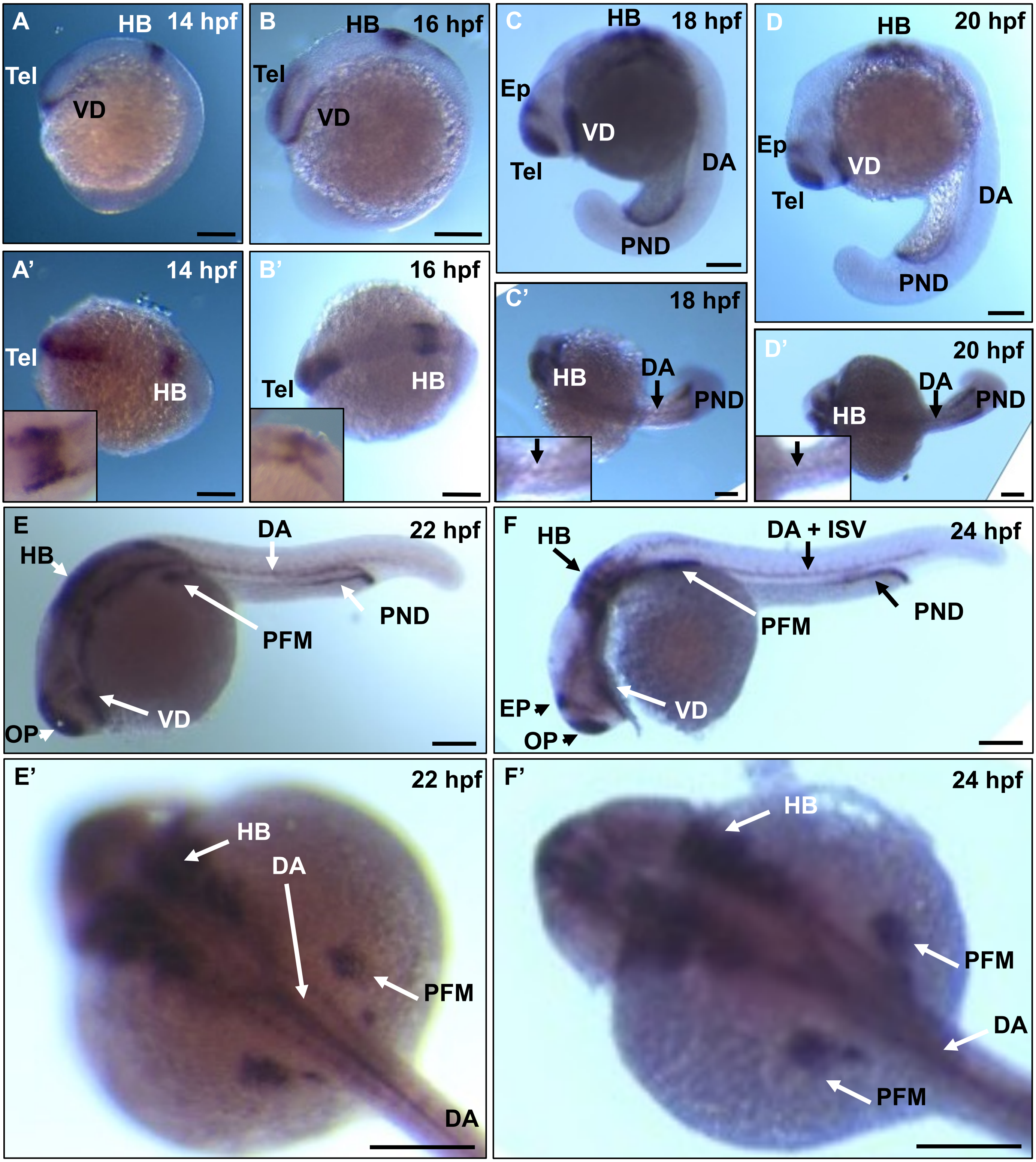
A
HB
14 hpf
Tel
VD
B
HB
16 hpf
Tel
VD
C
HB
18 hpf
Ep
VD
Tel
DA
PND
D
HB
20 hpf
Ep
VD
Tel
DA
PND
A'
14 hpf
Tel
HB
B'
16 hpf
Tel
HB
C'
18 hpf
HB
DA
PND
D'
20 hpf
DA
HB
PND
E
DA
22 hpf
HB
PFM
PND
VD
OP
F
HB
DA + ISV
24 hpf
PFM
PND
EP
VD
OP
E'
22 hpf
HB
DA
PFM
DA
F'
24 hpf
HB
PFM
DA
PFM

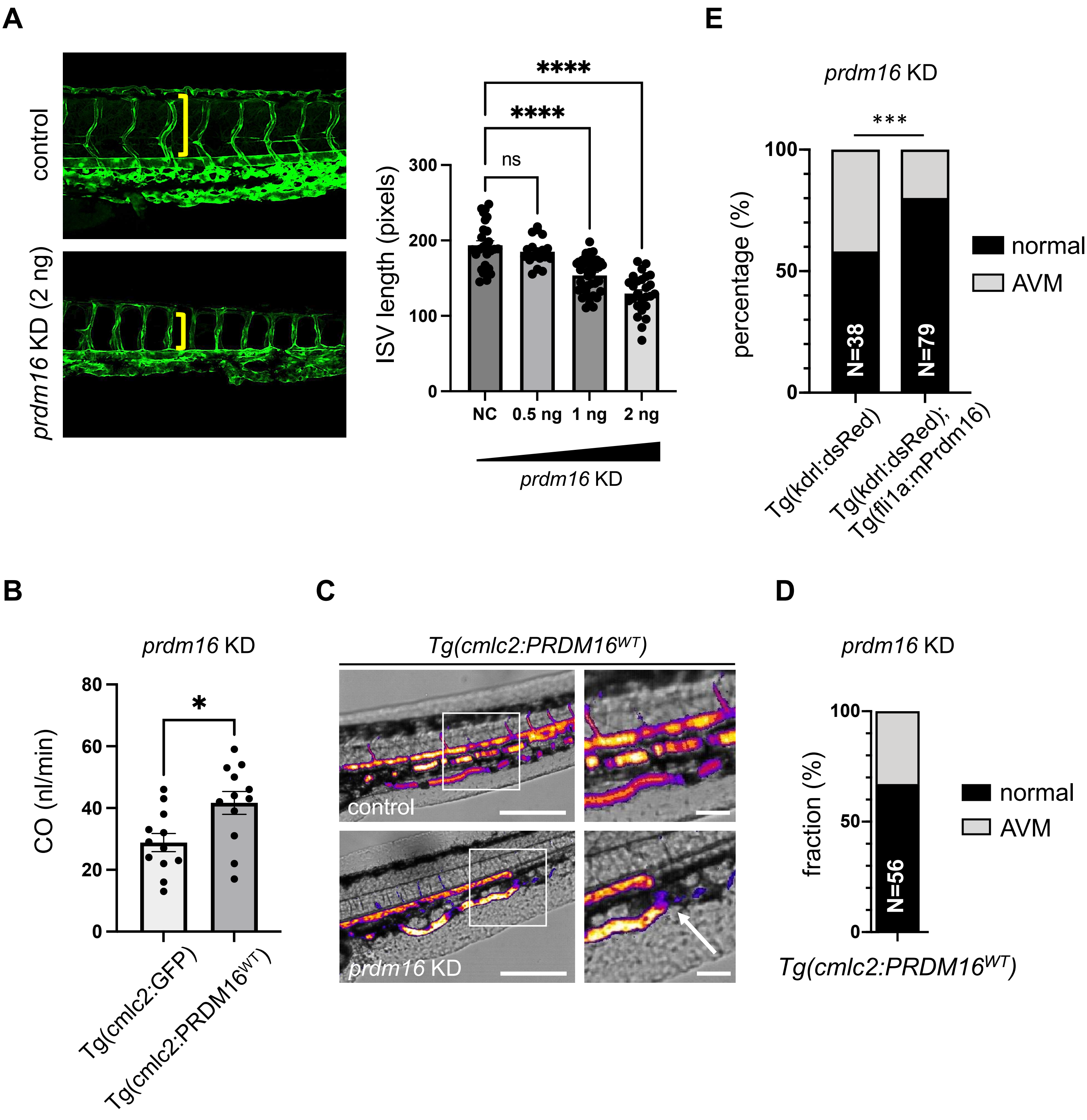
A
control
prdm16 KD (2 ng)
****
****
ns
ISV length (pixels)
300
200
100
0
NC
0.5 ng
1 ng
2 ng
prdm16 KD
E
prdm16 KD
***
100
50
0
percentage (%)
N=38
N=79
normal
AVM
Tg(kdrl:dsRed)
Tg(kdrl:dsRed);
Tg(fli1a:mPrdm16)
B
prdm16 KD
*
80
60
40
20
0
CO (nl/min)
Tg(cmlc2:GFP)
Tg(cmlc2:PRDM16^WT)
C
Tg(cmlc2:PRDM16^WT)
control
prdm16 KD
D
prdm16 KD
100
50
0
fraction (%)
N=56
normal
AVM
Tg(cmlc2:PRDM16^WT)

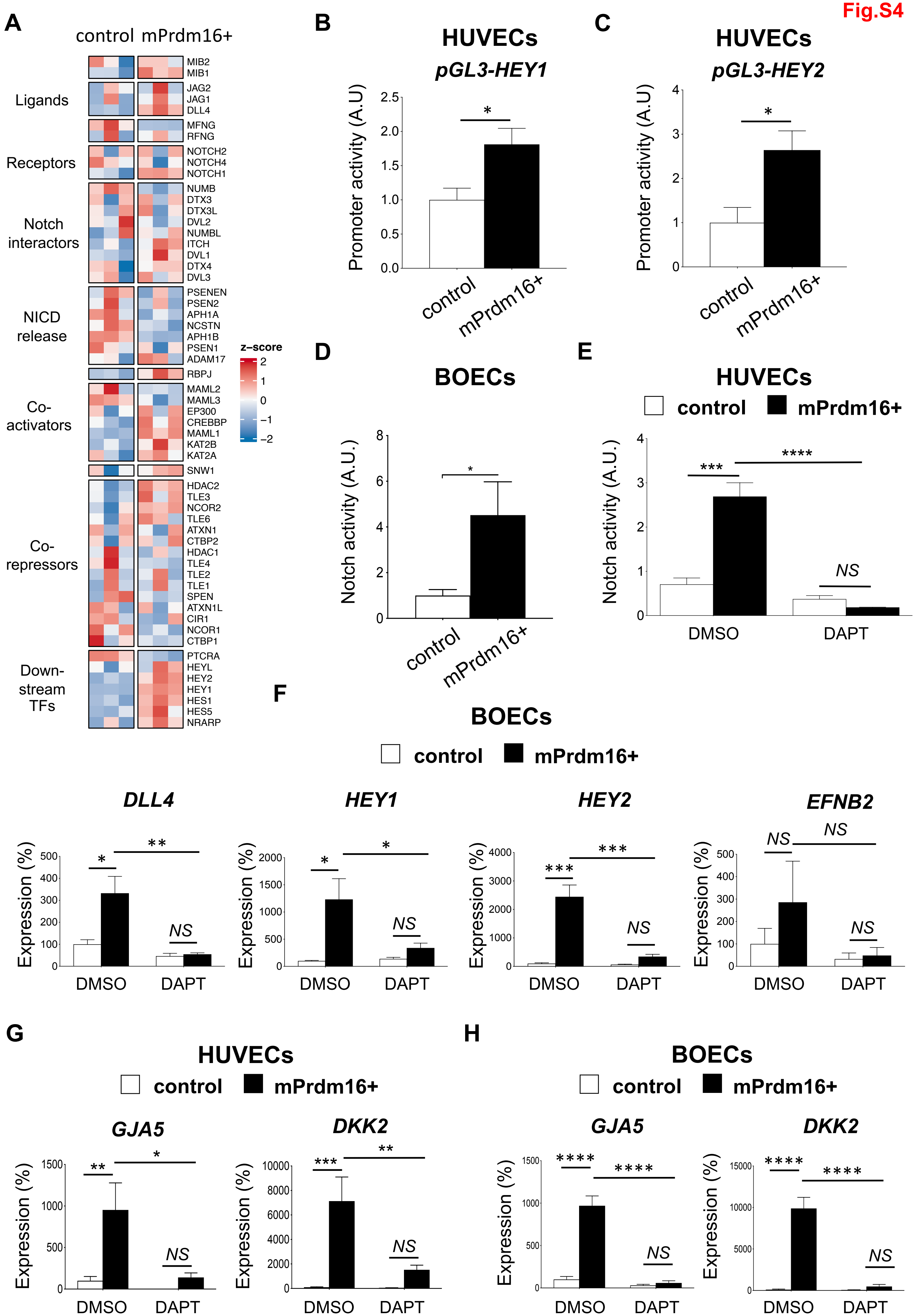
A
control mPrdm16+
Ligands
Receptors
Notch interactors
NICD release
Co-activators
Co-repressors
Down-stream TFs
MIB2
MIB1
JAG2
JAG1
DLL4
MFNG
RFNG
NOTCH2
NOTCH4
NOTCH1
NUMB
DTX3
DTX3L
DVL2
NUMBL
ITCH
DVL1
DTX4
DVL3
PSENEN
PSEN2
APH1A
NCSTN
APH1B
PSEN1
ADAM17
RBPJ
MAML2
MAML3
EP300
CREBBP
MAML1
KAT2B
KAT2A
SNW1
HDAC2
TLE3
NCOR2
TLE6
ATXN1
CTBP2
HDAC1
TLE4
TLE2
TLE1
SPEN
ATXN1L
CIR1
NCOR1
CTBP1
PTCRA
HEYL
HEY2
HEY1
HES1
HES5
NRARP
z-score
B
HUVECs
pGL3-HEY1
Promoter activity (A.U)
control
mPrdm16+
C
HUVECs
pGL3-HEY2
Promoter activity (A.U)
D
BOECs
Notch activity (A.U.)
E
HUVECs
control mPrdm16+
Notch activity (A.U.)
DMSO DAPT
NS
F
BOECs
control mPrdm16+
DLL4
HEY1
HEY2
EFNB2
Expression (%)
G
HUVECs
control mPrdm16+
GJA5
DKK2
H
BOECs
control mPrdm16+
GJA5
DKK2

Fig.S5

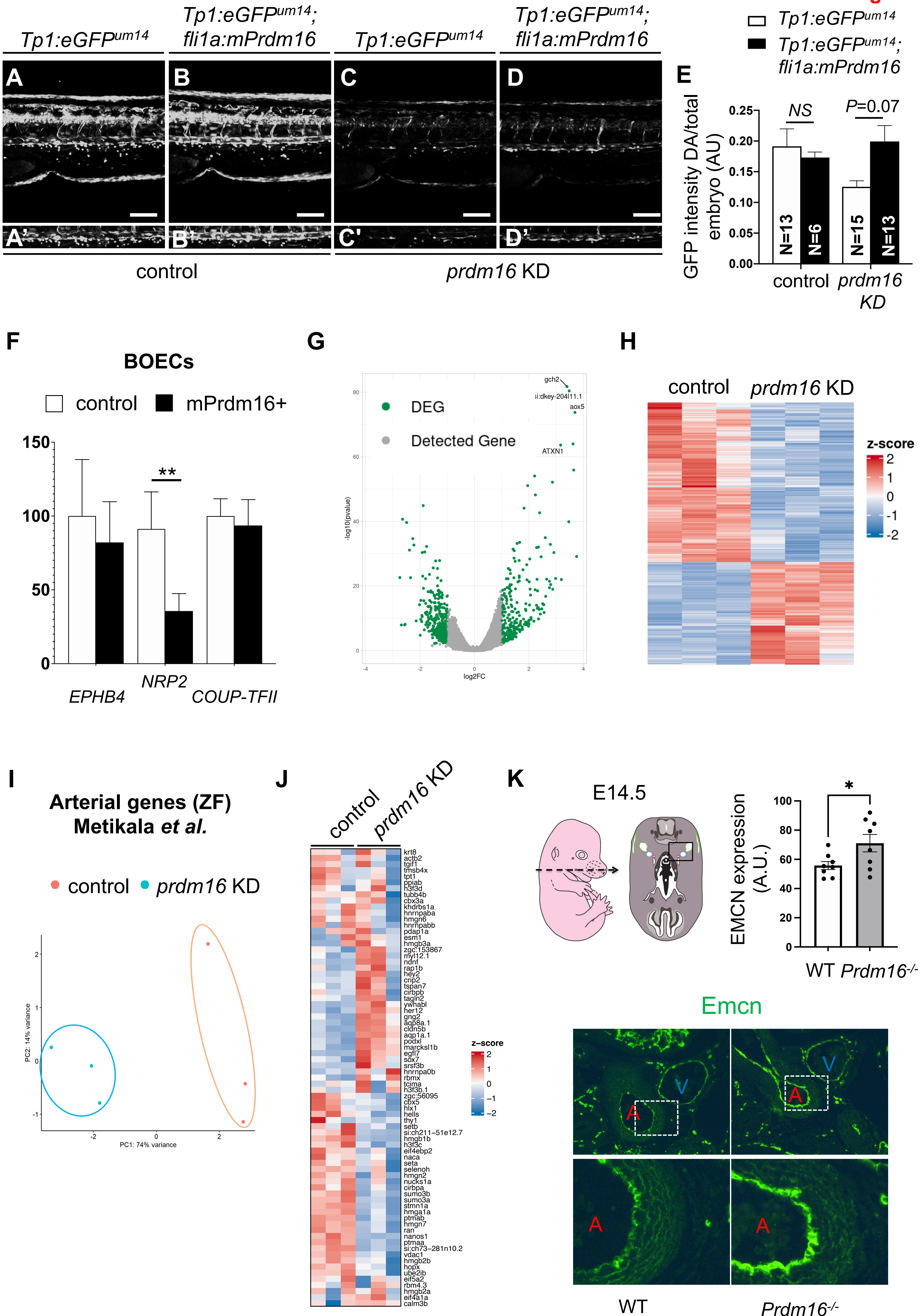

Tp1:eGFP^um14
Tp1:eGFP^um14; fli1a:mPrdm16
Tp1:eGFP^um14
Tp1:eGFP^um14; fli1a:mPrdm16
A
B
C
D
A'
B'
C'
D'
control
prdm16 KD
E
Tp1:eGFP^um14
Tp1:eGFP^um14; fli1a:mPrdm16
GFP intensity DA/total embryo (AU)
NS
P=0.07
N=13
N=6
N=15
N=13
0.25
0.20
0.15
0.10
0.05
0.00
control
prdm16 KD
F
BOECs
control
mPrdm16+
150
100
50
0
**
EPHB4
NRP2
COUP-TFII
G
DEG
Detected Gene
gch2
si:dkey-204l11.1
aox5
ATXN1
-log10(pvalue)
log2FC
H
control
prdm16 KD
z-score
2
1
0
-1
-2
I
Arterial genes (ZF)
Metikala et al.
control
prdm16 KD
PC2: 14% variance
PC1: 74% variance
J
control
prdm16 KD
krt8
actb2
tgif1
tmsb4x
tpt1
ppiab
h3f3d
tubb4b
cbx3a
khdrbs1a
hnrnpaba
hmgn6
hnrnpabb
pdap1a
esm1
hmgb3a
zgc:153867
myl12.1
ndnf
rap1b
hey2
crip2
tspan7
cirbpb
tagln2
ywhabl
her12
gng2
aqp8a.1
cldn5b
aqp1a.1
podxl
marcksl1b
egfl7
sox7
srsf3b
hnrnpa0b
rbmx
tcima
h3f3b.1
zgc:56095
cbx5
hlx1
hells
thy1
setb
si:ch211-51e12.7
hmgb1b
h3f3c
eif4ebp2
naca
seta
selenoh
hmgn2
nucks1a
cirbpa
sumo3b
sumo3a
stmn1a
hmga1a
ptmab
hmgn7
ran
nanos1
ptmaa
si:ch73-281n10.2
vdac1
hmgb2b
hopx
ube2ib
eif5a2
rbm4.3
hmgb2a
eif4a1a
calm3b
z-score
2
1
0
−1
−2
K
E14.5
EMCN expression (A.U.)
*
100
80
60
40
20
0
WT
Prdm16-/-
Emcn
V
A
WT
Prdm16-/-

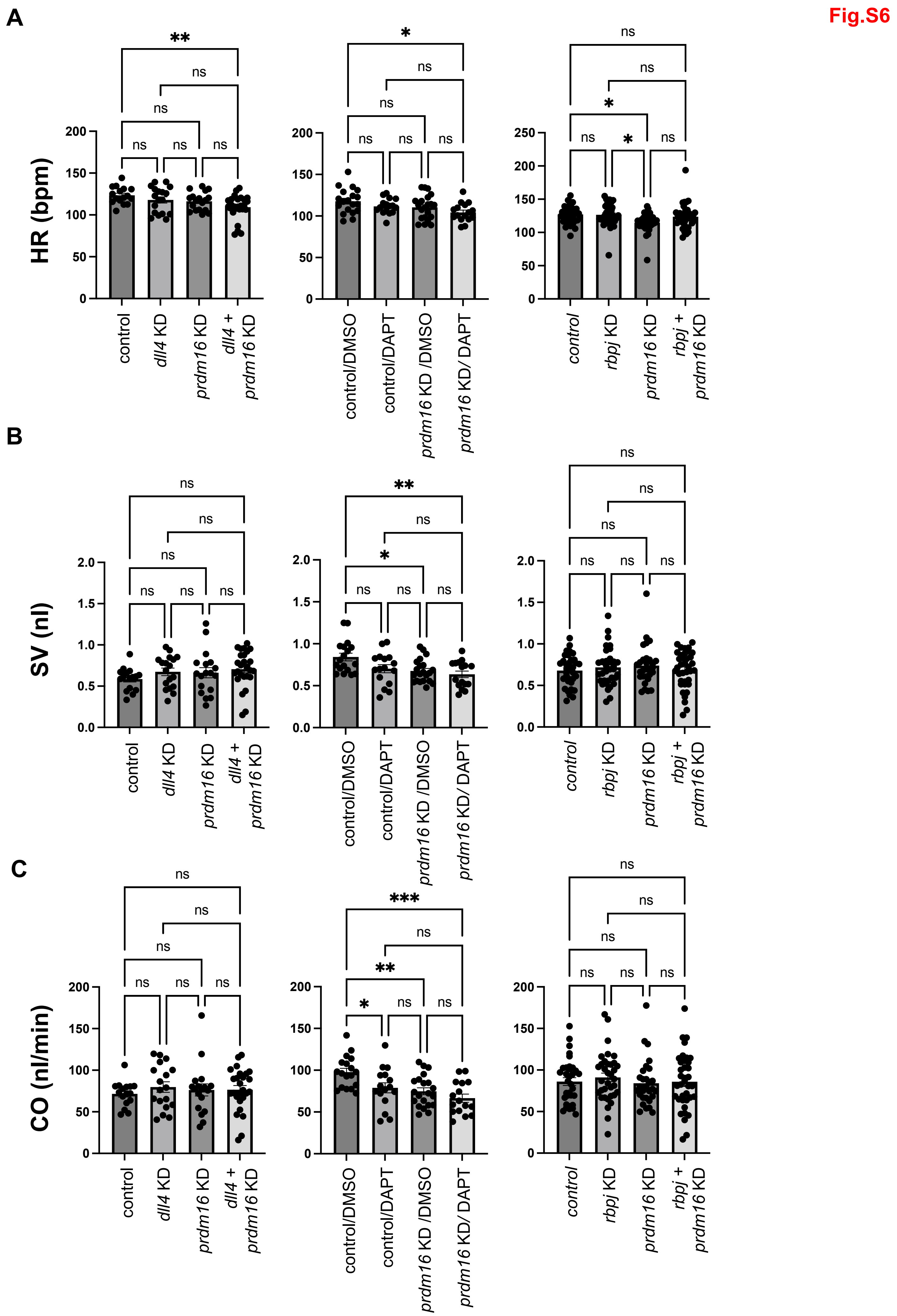

Fig.S6
A
B
C
HR (bpm)
SV (nl)
CO (nl/min)
**
*
***
ns
control
dll4 KD
prdm16 KD
dll4 + prdm16 KD
control/DMSO
control/DAPT
prdm16 KD /DMSO
prdm16 KD/ DAPT
rbpj KD
rbpj + prdm16 KD

Fig.S7

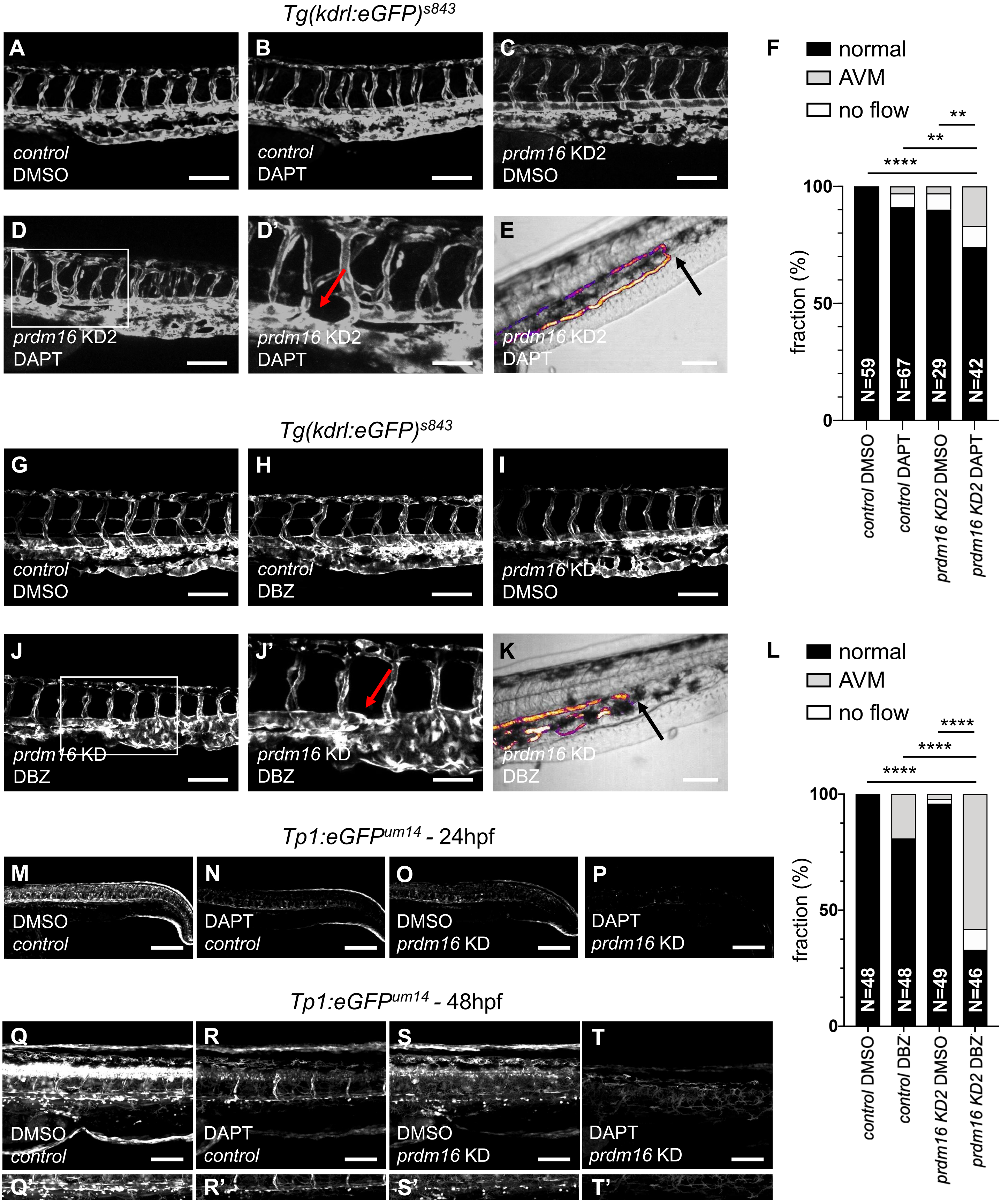
Tg(kdrl:eGFP)s843
A
control
DMSO
B
control
DAPT
C
prdm16 KD2
DMSO
D
prdm16 KD2
DAPT
D'
prdm16 KD2
DAPT
E
prdm16 KD2
DAPT
F
normal
AVM
no flow
**
**
****
fraction (%)
100
50
0
N=59
N=67
N=29
N=42
control DMSO
control DAPT
prdm16 KD2 DMSO
prdm16 KD2 DAPT
Tg(kdrl:eGFP)s843
G
control
DMSO
H
control
DBZ
I
prdm16 KD
DMSO
J
prdm16 KD
DBZ
J'
prdm16 KD
DBZ
K
prdm16 KD
DBZ
L
normal
AVM
no flow
****
****
****
fraction (%)
100
50
0
N=48
N=48
N=49
N=46
control DMSO
control DBZ
prdm16 KD2 DMSO
prdm16 KD2 DBZ
Tp1:eGFPum14 - 24hpf
M
DMSO
control
N
DAPT
control
O
DMSO
prdm16 KD
P
DAPT
prdm16 KD
Tp1:eGFPum14 - 48hpf
Q
DMSO
control
R
DAPT
control
S
DMSO
prdm16 KD
T
DAPT
prdm16 KD
Q'
R'
S'
T'

Fig.S8

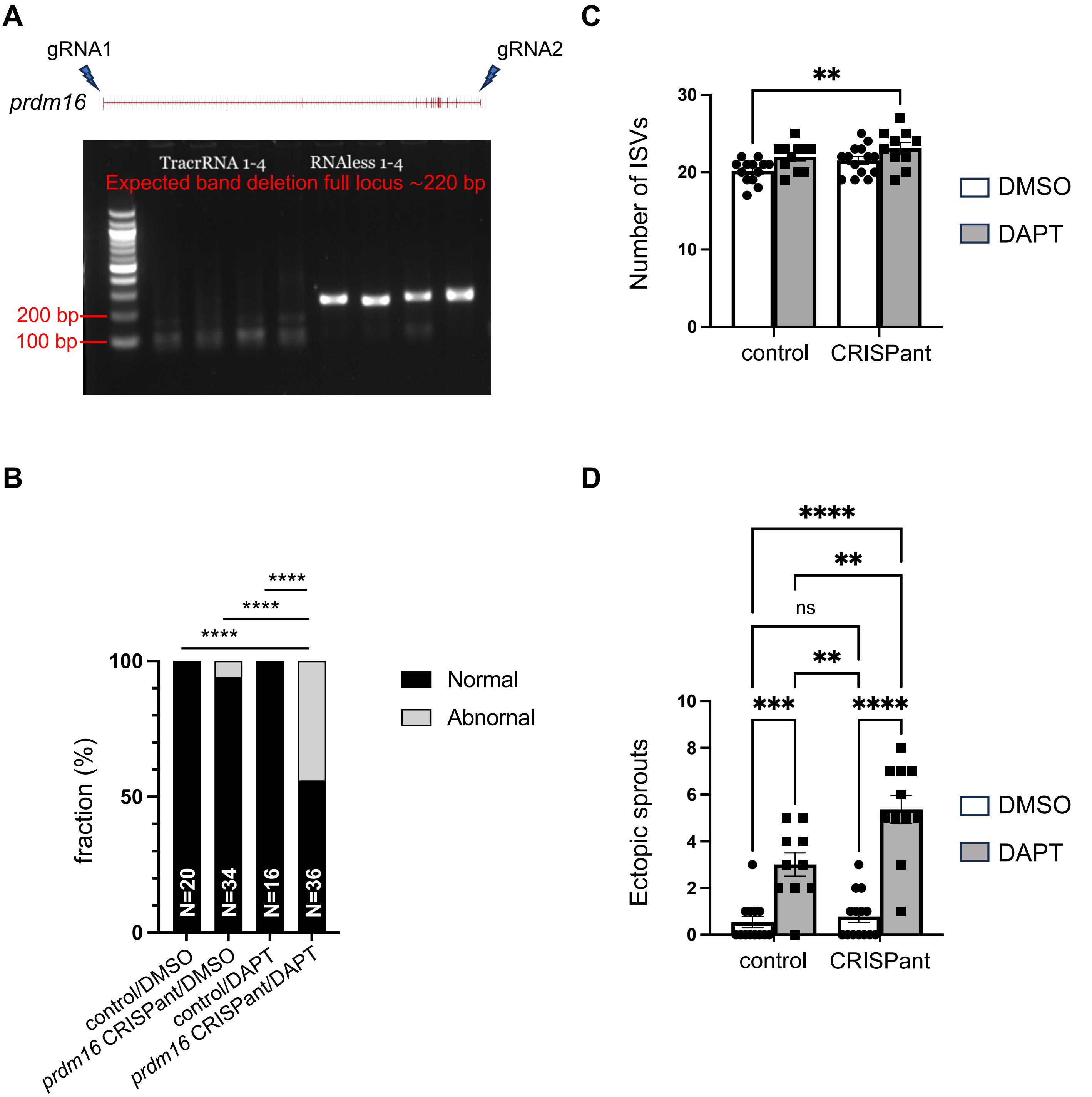
A
gRNA1
gRNA2
prdm16
TracrRNA 1-4
RNAless 1-4
Expected band deletion full locus ~220 bp
200 bp
100 bp
B
fraction (%)
100
50
0
****
****
****
N=20
N=34
N=16
N=36
control/DMSO
prdm16 CRISPant/DMSO
control/DAPT
prdm16 CRISPant/DAPT
Normal
Abnornal
C
Number of ISVs
30
20
10
0
**
control
CRISPant
DMSO
DAPT
D
Ectopic sprouts
10
8
6
4
2
0
****
**
ns
**
***
****
control
CRISPant
DMSO
DAPT

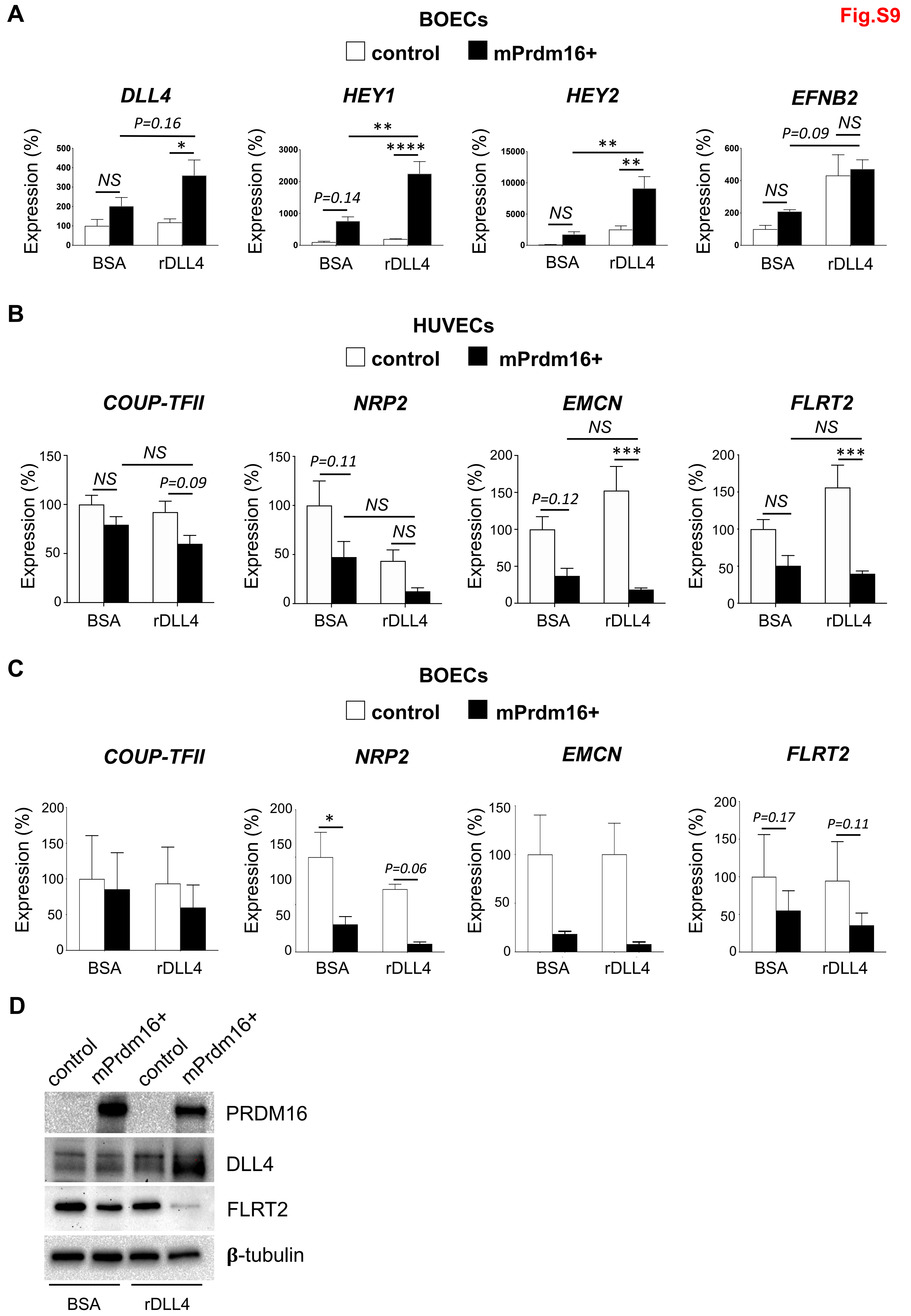
Fig.S9
A
BOECs
control
mPrdm16+
DLL4
HEY1
HEY2
EFNB2
Expression (%)
BSA
rDLL4
P=0.16
P=0.14
P=0.09
NS
B
HUVECs
COUP-TFII
NRP2
EMCN
FLRT2
P=0.09
P=0.11
P=0.12
C
P=0.06
P=0.17
D
PRDM16
DLL4
FLRT2
β-tubulin

**Fig.S10**

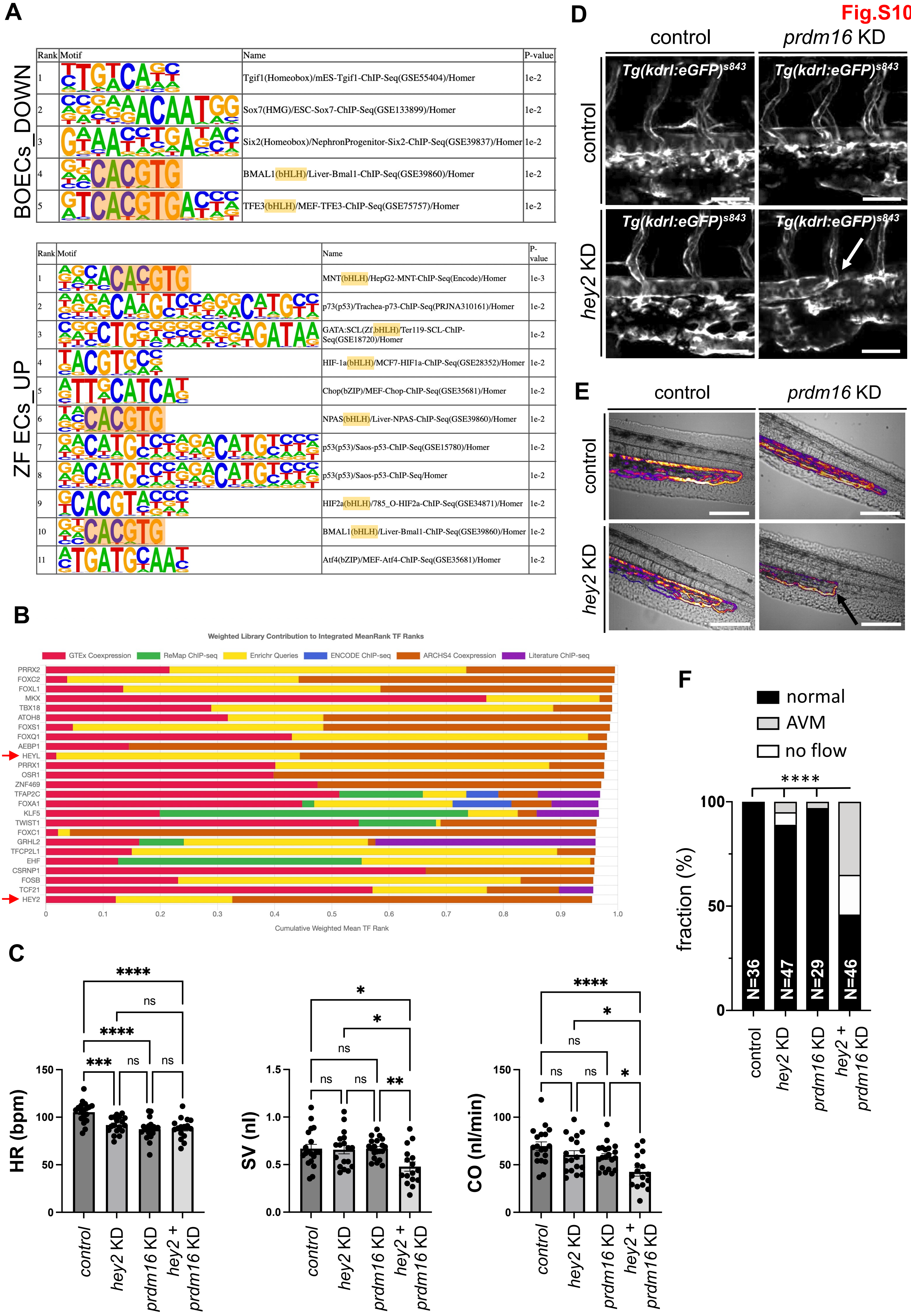
A
BOECs_DOWN
Rank | Motif | Name | P-value
1 | Tgif1(Homeobox)/mES-Tgif1-ChIP-Seq(GSE55404)/Homer | 1e-2
2 | Sox7(HMG)/ESC-Sox7-ChIP-Seq(GSE133899)/Homer | 1e-2
3 | Six2(Homeobox)/NephronProgenitor-Six2-ChIP-Seq(GSE39837)/Homer | 1e-2
4 | BMAL1(bHLH)/Liver-Bmal1-ChIP-Seq(GSE39860)/Homer | 1e-2
5 | TFE3(bHLH)/MEF-TFE3-ChIP-Seq(GSE75757)/Homer | 1e-2
ZF ECs_UP
Rank | Motif | Name | P-value
1 | MNT(bHLH)/HepG2-MNT-ChIP-Seq(Encode)/Homer | 1e-3
2 | p73(p53)/Trachea-p73-ChIP-Seq(PRJNA310161)/Homer | 1e-2
3 | GATA:SCL(Zf,bHLH)/Ter119-SCL-ChIP-Seq(GSE18720)/Homer | 1e-2
4 | HIF-1a(bHLH)/MCF7-HIF1a-ChIP-Seq(GSE28352)/Homer | 1e-2
5 | Chop(bZIP)/MEF-Chop-ChIP-Seq(GSE35681)/Homer | 1e-2
6 | NPAS(bHLH)/Liver-NPAS-ChIP-Seq(GSE39860)/Homer | 1e-2
7 | p53(p53)/Saos-p53-ChIP-Seq(GSE15780)/Homer | 1e-2
8 | p53(p53)/Saos-p53-ChIP-Seq/Homer | 1e-2
9 | HIF2a(bHLH)/785_O-HIF2a-ChIP-Seq(GSE34871)/Homer | 1e-2
10 | BMAL1(bHLH)/Liver-Bmal1-ChIP-Seq(GSE39860)/Homer | 1e-2
11 | Atf4(bZIP)/MEF-Atf4-ChIP-Seq(GSE35681)/Homer | 1e-2
B
Weighted Library Contribution to Integrated MeanRank TF Ranks
GTEx Coexpression
ReMap ChIP-seq
Enrichr Queries
ENCODE ChIP-seq
ARCHS4 Coexpression
Literature ChIP-seq
PRRX2
FOXC2
FOXL1
MKX
TBX18
ATOH8
FOXS1
FOXQ1
AEBP1
HEYL
PRRX1
OSR1
ZNF469
TFAP2C
FOXA1
KLF5
TWIST1
FOXC1
GRHL2
TFCP2L1
EHF
CSRNP1
FOSB
TCF21
HEY2
0 0.1 0.2 0.3 0.4 0.5 0.6 0.7 0.8 0.9 1.0
Cumulative Weighted Mean TF Rank
C
HR (bpm)
SV (nl)
CO (nl/min)
****
ns
****
***
ns
ns
*
*
ns
ns
ns
**
****
*
ns
ns
ns
*
control
hey2 KD
prdm16 KD
hey2 + prdm16 KD
D
control
prdm16 KD
control
hey2 KD
Tg(kdrl:eGFP)s843
E
control
prdm16 KD
control
hey2 KD
F
normal
AVM
no flow
****
fraction (%)
100
50
0
N=36
N=47
N=29
N=46
control
hey2 KD
prdm16 KD
hey2 + prdm16 KD

TableS1

#### Supplementary Table S1. List of conserved genes enriched in arterial or venous ECs within the vasculature

| ARTERIAL (N=22) | |
|---|---|
| *Aim1* | *Gja5* |
| *Antxr1* | ***Hey2*** |
| *Atp2a3* | *Ins-Igf2* |
| *Bcap29* | *Lox* |
| *Camk4* | *Nox4* |
| *Ccdc3* | ***Prdm16*** |
| *Cp* | *Ptprj* |
| *Dkk2* | *Rps6ka5* |
| *Edn1* | *Slc4a4* |
| *Efna5* | *Thbs1* |
| *Ehd4* | ***Tox*** |

| VENOUS (N=30) | |
|---|---|
| *Abcg2* | *Lyve1* |
| *Bex2* | ***Meox2*** |
| *Cdh11* | *Mest* |
| *Chn2* | *Nostrin* |
| *Ctsk* | ***Coup-TFII*** |
| *Dcn* | *Nrp2* |
| *Emcn* | *Pitpnc1* |
| *Enpp2* | *Pkhd1l1* |
| *Entpd1* | ***Prox1*** |
| *Flrt2* | *Pthlh* |
| *Fmnl2* | *Rgs5* |
| *Gpm6a* | *Samd5* |
| *Itgam* | *Sntb2* |
| *Lrg1* | ***Znf462*** |
| *Ltbp1* | ***Znf521*** |

### Supplementary Table S2. Gene Ontology (GO) analysis for genes with conserved arterial or venous-restricted expression pattern

| Gene Set Name | # Genes in Gene Set (K) | # Genes in Overlap (k) | k/K | FDR q-value |
|---|---|---|---|---|
| GOBP_CIRCULATORY_SYSTEM_DEVELOPMENT | 1224 | 15 | 0.0123 | 8.39E-8 |
| GOBP_CELL_MOTILITY | 1813 | 17 | 0.0094 | 8.9E-8 |
| GOBP_RESPONSE_TO_GROWTH_FACTOR | 764 | 12 | 0.0157 | 2.69E-7 |
| GOBP_VASCULATURE_DEVELOPMENT | 832 | 12 | 0.0144 | 5.32E-7 |
| GOBP_LOCOMOTION | 1335 | 14 | 0.0105 | 7.06E-7 |
| GOBP_BLOOD_VESSEL_MORPHOGENESIS | 709 | 11 | 0.0155 | 9.52E-7 |
| GOBP_TISSUE_MIGRATION | 382 | 9 | 0.0236 | 9.52E-7 |
| GOBP_REGULATION_OF_EPITHELIAL_CELL_MIGRATION | 296 | 8 | 0.0270 | 2.43E-6 |
| GOBP_RESPONSE_TO_TRANSFORMING_GROWTH_FACTOR_BETA | 297 | 8 | 0.0269 | 2.43E-6 |
| GOBP_AMEBOIDAL_TYPE_CELL_MIGRATION | 500 | 9 | 0.0180 | 6.73E-6 |
| GOBP_REGULATION_OF_LOCOMOTION | 1142 | 12 | 0.0105 | 6.73E-6 |
| GOBP_REGULATION_OF_ENDOTHELIAL_CELL_MIGRATION | 232 | 7 | 0.0302 | 8.81E-6 |
| GOBP_TUBE_MORPHOGENESIS | 953 | 11 | 0.0115 | 9.66E-6 |
| GOBP_REGULATION_OF_MULTICELLULAR_ORGANISMAL_DEVELOPMENT | 1484 | 13 | 0.0088 | 9.66E-6 |
| GOBP_CELL_MORPHOGENESIS | 964 | 11 | 0.0114 | 9.66E-6 |
| GOBP_CELL_JUNCTION_ORGANIZATION | 758 | 10 | 0.0132 | 1.13E-5 |
| GOBP_MUSCLE_TISSUE_DEVELOPMENT | 434 | 8 | 0.0184 | 2.33E-5 |
| GOBP_HEART_DEVELOPMENT | 618 | 9 | 0.0146 | 2.33E-5 |
| GOBP_ENDOTHELIAL_CELL_MIGRATION | 286 | 7 | 0.0245 | 2.33E-5 |
| GOBP_TUBE_DEVELOPMENT | 1169 | 11 | 0.0094 | 5.08E-5 |
| GOBP_RESPONSE_TO_ENDOGENOUS_STIMULUS | 1777 | 13 | 0.0073 | 5.41E-5 |
| GOBP_GENERATION_OF_NEURONS | 1485 | 12 | 0.0081 | 5.89E-5 |
| GOBP_ARTERY_DEVELOPMENT | 111 | 5 | 0.0450 | 7.86E-5 |
| GOBP_CELL_SUBSTRATE_ADHESION | 356 | 7 | 0.0197 | 7.86E-5 |
| GOBP_NEGATIVE_REGULATION_OF_MULTICELLULAR_ORGANISMAL_PROCESS | 1249 | 11 | 0.0088 | 7.86E-5 |
| GOBP_POSITIVE_REGULATION_OF_CELL_POPULATION_PROLIFERATION | 1286 | 11 | 0.0086 | 1.01E-4 |
| GOBP_ENZYME_LINKED_RECEPTOR_PROTEIN_SIGNALING_PATHWAY | 1022 | 10 | 0.0098 | 1.06E-4 |
| GOBP_TRANSFORMING_GROWTH_FACTOR_BETA_RECEPTOR_SIGNALING_PATHWAY | 233 | 6 | 0.0258 | 1.11E-4 |
| GOBP_CELLULAR_COMPONENT_MORPHOGENESIS | 789 | 9 | 0.0114 | 1.14E-4 |
| GOBP_CELL_MATRIX_ADHESION | 239 | 6 | 0.0251 | 1.18E-4 |
| GOBP_PROTEIN_PHOSPHORYLATION | 1330 | 11 | 0.0083 | 1.18E-4 |
| GOBP_POSITIVE_REGULATION_OF_DEVELOPMENTAL_PROCESS | 1370 | 11 | 0.0080 | 1.46E-4 |
| GOBP_POSITIVE_REGULATION_OF_MULTICELLULAR_ORGANISMAL_PROCESS | 1684 | 12 | 0.0071 | 1.46E-4 |
| GOBP_CIRCULATORY_SYSTEM_PROCESS | 602 | 8 | 0.0133 | 1.46E-4 |
| GOBP_NEUROGENESIS | 1715 | 12 | 0.0070 | 1.71E-4 |
| GOBP_HEART_MORPHOGENESIS | 265 | 6 | 0.0226 | 1.78E-4 |
| GOBP_STRIATED_MUSCLE_TISSUE_DEVELOPMENT | 265 | 6 | 0.0226 | 1.78E-4 |
| GOBP_HEART_VALVE_FORMATION | 16 | 3 | 0.1875 | 1.9E-4 |
| GOBP_CELL_JUNCTION_ASSEMBLY | 447 | 7 | 0.0157 | 2.29E-4 |
| GOBP_CELL_FATE_COMMITMENT | 286 | 6 | 0.0210 | 2.57E-4 |
| GOBP_CELL_PART_MORPHOGENESIS | 680 | 8 | 0.0118 | 2.96E-4 |
| GOBP_TAXIS | 470 | 7 | 0.0149 | 2.96E-4 |
| GOBP_REGULATION_OF_CELLULAR_RESPONSE_TO_TRANSFORMING_GROWTH_FACTOR_BETA_STIMULUS | 164 | 5 | 0.0305 | 3.13E-4 |
| GOBP_CELL_ADHESION | 1531 | 11 | 0.0072 | 3.28E-4 |
| GOBP_POSITIVE_REGULATION_OF_TRANSCRIPTION_BY_RNA_POLYMERASE_II | 1241 | 10 | 0.0081 | 3.66E-4 |
| GOBP_ANATOMICAL_STRUCTURE_FORMATION_INVOLVED_IN_MORPHOGENESIS | 1259 | 10 | 0.0079 | 4.07E-4 |
| GOBP_POSITIVE_REGULATION_OF_EPITHELIAL_CELL_MIGRATION | 177 | 5 | 0.0282 | 4.17E-4 |
| GOBP_PHOSPHORYLATION | 1596 | 11 | 0.0069 | 4.49E-4 |
| GOBP_ARTERY_MORPHOGENESIS | 81 | 4 | 0.0494 | 4.73E-4 |
| GOBP_OUTFLOW_TRACT_MORPHOGENESIS | 82 | 4 | 0.0488 | 4.86E-4 |

### Supplementary Table S3.

| Probe Set ID | Gene Symbol | Log-ratio |
|---|---|---|
| 203789_s_at | *SEMA3C* | 6,0437293999909 |
| 205190_at | *PLS1* | 5,22401858763448 |
| 206023_at | *NMU* | 5,22151097233681 |
| 206924_at | *IL11* | 4,68662270149357 |
| 222162_s_at | *ADAMTS1* | 4,52294128336308 |
| 224209_s_at | *GDA* | 4,39372023101228 |
| 222486_s_at | *ADAMTS1* | 4,23755056820467 |
| 206315_at | *CRLF1* | 4,21629423584944 |
| 206172_at | *IL13RA2* | 4,01675022510488 |
| 1556499_s_at | *COL1A1* | 3,94329846511255 |
| 202037_s_at | *SFRP1* | 3,83690250100065 |
| 228186_s_at | *RSPO3* | 3,80047257319591 |
| 215870_s_at | *PLA2G5* | 3,76370275121388 |
| 219529_at | *CLIC3* | 3,75980008772369 |
| 1553413_at | *FLJ13744* | 3,7443689522623 |
| 202409_at | *IGF2 /// INS-IGF2* | 3,58216736485246 |
| 214321_at | *NOV* | 3,35119305028312 |
| 206091_at | *MATN3* | 3,2637576453234 |
| 231766_s_at | *COL12A1* | 3,2467522134792 |
| 229147_at | *RASSF6* | 3,21600072349649 |
| 201427_s_at | *SEPP1* | 3,17590454124338 |
| 220468_at | *ARL14* | 3,15784668937622 |
| 226803_at | *CHMP4C* | 3,11704628020672 |
| 204637_at | *CGA* | 3,08094463175001 |
| 225548_at | *SHROOM3* | 3,07589700748369 |
| 224650_at | *MAL2* | 3,04389813841765 |
| 232164_s_at | *EPPK1* | 2,9982542020504 |
| 210372_s_at | *TPD52L1* | 2,95676354388968 |
| 213280_at | *RAP1GAP2* | 2,88707668340002 |
| 203108_at | *GPRC5A* | 2,86373891150044 |
| 212444_at | --- | 2,85373026518313 |
| 224516_s_at | *CXXC5* | 2,85247683867703 |
| 203786_s_at | *TPD52L1* | 2,83998166350043 |
| 203088_at | *FBLN5* | 2,79588126434779 |
| 205081_at | *CRIP1* | 2,78754430078725 |
| 226701_at | *GJA5* | 2,7855002977454 |
| 218404_at | *SNX10* | 2,77541028325866 |
| 239218_at | *PDE1C* | 2,73957086066001 |
| 244455_at | *KCNT2* | 2,73905348423647 |
| 228400_at | *SHROOM3* | 2,73314116476238 |
| 218332_at | *BEX1* | 2,72627499047362 |
| 234103_at | *KCNT2* | 2,65128326226068 |
| 229872_s_at | *LOC100132999* | 2,64398506445126 |
| 204268_at | *S100A2* | 2,62397757550335 |
| 233955_x_at | *CXXC5* | 2,61371956412411 |
| 206683_at | *ZNF165* | 2,60246857424034 |
| 236344_at | *PDE1C* | 2,60023692720396 |
| 221731_x_at | *VCAN* | 2,59541397712453 |
| 228423_at | *MAP9* | 2,59248859560428 |
| 222921_s_at | *HEY2* | 2,56675178752357 |
| 201839_s_at | *EPCAM* | 2,56036204598664 |
| 202890_at | *MAP7* | 2,55888720860849 |
| 212670_at | *ELN* | 2,54238603451272 |
| 230746_s_at | *LOC100288985* | 2,5377383816172 |
| 204485_s_at | *TOM1L1* | 2,50570315257178 |
| 209291_at | *ID4* | 2,43839931179474 |
| 204597_x_at | *STC1* | 2,4236016718966 |
| 201422_at | *IFI30* | 2,40282134529388 |
| 204620_s_at | *VCAN* | 2,39549870984118 |
| 228450_at | *PLEKHA7* | 2,37646067007187 |
| 227099_s_at | *C11orf96* | 2,37152699848548 |
| 229485_x_at | *SHISA3* | 2,36812749434374 |
| 225325_at | *MFSD6* | 2,36621515374093 |
| 204790_at | *SMAD7* | 2,3450904145901 |
| 211756_at | *PTHLH* | 2,34203510661309 |
| 228900_at | *SPECC1* | 2,31606001261042 |
| 232060_at | *ROR1* | 2,31035118071534 |
| 227478_at | *SETBP1* | 2,28957229773957 |
| 227372_s_at | *BAIAP2L1* | 2,28898265580399 |
| 200606_at | *DSP* | 2,26700749883868 |
| 44783_s_at | *HEY1* | 2,25996554101627 |
| 203817_at | *GUCY1B3* | 2,25734361406301 |
| 204595_s_at | *STC1* | 2,25314350679847 |
| 232165_at | *EPPK1* | 2,24637462163419 |
| 238481_at | *MGP* | 2,24179487789693 |
| 230563_at | *RASGEF1A* | 2,23446628179923 |
| 213800_at | *CFH* | 2,22931216467858 |
| 210511_s_at | *INHBA* | 2,22617404834073 |
| 227180_at | *ELOVL7* | 2,2233716684058 |
| 227140_at | *INHBA* | 2,21658156255373 |
| 219743_at | *HEY2* | 2,21204937309636 |
| 213618_at | *ARAP2* | 2,19761831728086 |
| 228739_at | *CYS1* | 2,17597840760864 |
| 228407_at | *SCUBE3* | 2,17575655568873 |
| 235775_at | *TMTC2* | 2,1738226895735 |
| 229963_at | *BEX5* | 2,16471783962849 |
| 218532_s_at | *FAM134B* | 2,15177195371168 |
| 204379_s_at | *FGFR3* | 2,14976068389766 |
| 203886_s_at | *FBLN2* | 2,13055206708623 |
| 207076_s_at | *ASS1* | 2,12224750017496 |
| 209343_at | *EFHD1* | 2,1150874108785 |
| 205206_at | *KAL1* | 2,11465014502298 |
| 217901_at | *DSG2* | 2,11212827984334 |
| 209695_at | *PTP4A3* | 2,10927281909759 |
| 209821_at | *IL33* | 2,10071405035096 |
| 218839_at | *HEY1* | 2,07715280935582 |
| 226568_at | *FAM102B* | 2,07259445081075 |
| 205257_s_at | *AMPH* | 2,05468499451034 |
| 225597_at | *SLC45A4* | 2,05121782881994 |
| 231879_at | *COL12A1* | 2,04684289107869 |
| 217551_at | *OR7E14P* | 2,04261914492913 |
| 228574_at | *TMTC2* | 2,03973679761089 |
| 220145_at | *MAP9* | 2,02820126458491 |
| 244461_at | *SPECC1* | 2,02432469343179 |
| 207069_s_at | *SMAD6* | 2,01000625202211 |
| 204151_x_at | *AKR1C1* | 2,0020644904357 |
| 219305_x_at | *FBXO2* | 1,97697104978289 |
| 227764_at | *LYPD6* | 1,94546602882747 |
| 222996_s_at | *CXXC5* | 1,94184031094537 |
| 204679_at | *KCNK1* | 1,92195427973791 |
| 227763_at | *LYPD6* | 1,92165910334657 |
| 40687_at | *GJA4* | 1,91698973682684 |
| 232615_at | --- | 1,90113211456885 |
| 64408_s_at | *CALML4* | 1,89908957912429 |
| 209160_at | *AKR1C3* | 1,88998460360399 |
| 210095_s_at | *IGFBP3* | 1,85209382253617 |
| 221879_at | *CALML4* | 1,8442610545246 |
| 202404_s_at | *COL1A2* | 1,8433906820215 |
| 242162_at | *WDR69* | 1,84289340013516 |
| 205805_s_at | *ROR1* | 1,83632109804013 |
| 205997_at | *ADAM28* | 1,82464122403379 |
| 225664_at | *COL12A1* | 1,82357461253805 |
| 213652_at | *PCSK5* | 1,81093278136482 |
| 1559394_a_at | --- | 1,80497454426827 |
| 223498_at | *SPECC1* | 1,79804084566283 |
| 219858_s_at | *MFSD6* | 1,78592535672427 |
| 202672_s_at | *ATF3* | 1,77462600160498 |
| 213106_at | *ATP8A1* | 1,77195884985198 |
| 244503_at | --- | 1,75840880396364 |
| 235165_at | *PARD6B* | 1,7555232014465 |
| 206382_s_at | *BDNF* | 1,75325493532947 |
| 226641_at | *ANKRD44* | 1,75242323232806 |
| 237301_at | --- | 1,717292898717 |
| 228062_at | *NAP1L5* | 1,71501192804771 |
| 203951_at | *CNN1* | 1,71492993724971 |
| 226847_at | *FST* | 1,71272605608834 |
| 231382_at | *FGF18* | 1,71126351377066 |
| 210715_s_at | *SPINT2* | 1,70736431227057 |
| 232687_at | --- | 1,67415335198543 |
| 1570155_at | --- | 1,66661204294078 |
| 203476_at | *TPBG* | 1,66454675346041 |
| 203895_at | *PLCB4* | 1,66114068782078 |
| 230493_at | *SHISA2* | 1,65932314612718 |
| 225136_at | *PLEKHA2* | 1,6516391704527 |
| 228850_s_at | *SLIT2* | 1,65145274212064 |
| 203592_s_at | *FSTL3* | 1,64947279734857 |
| 206574_s_at | *PTP4A3* | 1,6472954104296 |
| 204904_at | *GJA4* | 1,64411768758284 |
| 1555963_x_at | *B3GNT7* | 1,64040430002324 |
| 205559_s_at | *PCSK5* | 1,63850687557512 |

### Supplementary Table S3 (continued)

| | | |
|---|---|---|
| 200704_at | *LITAF* | 1,63115594674005 |
| 221841_s_at | *KLF4* | 1,63107053367741 |
| 202035_s_at | *SFRP1* | 1,63068882441903 |
| 213817_at | *IRAK3* | 1,62484536068267 |
| 241418_at | *LOC344887* | 1,62024875743745 |
| 205624_at | *CPA3* | 1,61666322897343 |
| 212873_at | *HMHA1* | 1,61161429616213 |
| 209699_x_at | *AKR1C2* | 1,60655641580992 |
| 213201_s_at | *TNNT1* | 1,60424805987121 |
| 205933_at | *SETBP1* | 1,60218500915685 |
| 230130_at | --- | 1,58126214108179 |
| 1552309_a_at | *NEXN* | 1,57549265518461 |
| 226103_at | *NEXN* | 1,56937088436824 |
| 238029_s_at | *SLC16A14* | 1,56879894855642 |
| 223681_s_at | *INADL* | 1,56643276938134 |
| 202901_x_at | *CTSS* | 1,5640863918127 |
| 225598_at | *SLC45A4* | 1,56083986486041 |
| 230793_at | *LRRC16A* | 1,53842989378623 |
| 219474_at | *C3orf52* | 1,53830445756622 |
| 228063_s_at | *NAP1L5* | 1,53719298471305 |
| 226258_at | *AMN1* | 1,5359276889826 |
| 238480_at | *TTC39C* | 1,53276579819906 |
| 211653_x_at | *AKR1C2* | 1,53259611848333 |
| 206178_at | *PLA2G5* | 1,52922185941962 |
| 204422_s_at | *FGF2* | 1,52817808636986 |
| 203397_s_at | *GALNT3* | 1,50797836220732 |
| 212353_at | *SULF1* | 1,50613331598573 |
| 219410_at | *TMEM45A* | 1,50390919704206 |
| 227058_at | *C13orf33* | 1,50194567798858 |
| 231332_at | --- | 1,49405259281984 |
| 211555_s_at | *GUCY1B3* | 1,48008005734526 |
| 202036_s_at | *SFRP1* | 1,47355178592341 |
| 231397_at | *LPPR5* | 1,47257045293958 |
| 1560475_at | *LOC100129455* | 1,47139114412989 |
| 218499_at | *MST4* | 1,46695791396583 |
| 209198_s_at | *SYT11* | 1,46115860820089 |
| 202826_at | *SPINT1* | 1,45887585128518 |
| 202720_at | *TES* | 1,45858362158986 |
| 205027_s_at | *MAP3K8* | 1,45280374005515 |
| 216594_x_at | *AKR1C1* | 1,45128419782761 |
| 223467_at | *RASD1* | 1,45079558574032 |
| 208131_s_at | *PTGIS* | 1,44058819624788 |
| 215717_s_at | *FBN2* | 1,4362394884224 |
| 204421_s_at | *FGF2* | 1,43426510976849 |
| 203973_s_at | *CEBPD* | 1,42544026417742 |
| 225283_at | *ARRDC4* | 1,42511148126204 |
| 204939_s_at | *PLN* | 1,42295278805752 |
| 242722_at | *LMO7* | 1,41991174127032 |
| 238441_at | *PRKAA2* | 1,4137959264497 |
| 226498_at | *FLT1* | 1,41312886404166 |
| 218718_at | *PDGFC* | 1,41297969905727 |
| 220180_at | *CCDC68* | 1,41170123963274 |
| 205239_at | *AREG* | 1,41161224248261 |
| 231906_at | *HOXD8* | 1,41145315132539 |
| 216869_at | *PDE1C* | 1,40765173094679 |
| 221261_x_at | *MAGED4 /// MAGED4B* | 1,40601547279749 |
| 209197_at | *SYT11* | 1,39989629142344 |
| 202403_s_at | *COL1A2* | 1,39786023583135 |
| 204584_at | *L1CAM* | 1,39744195695343 |
| 218631_at | *AVPI1* | 1,39731951042372 |
| 225975_at | *PCDH18* | 1,39443266128755 |
| 210355_at | *PTHLH* | 1,39011484526744 |
| 214719_at | *SLC46A3* | 1,3884530229409 |
| 228471_at | *ANKRD44* | 1,38815154414413 |
| 205729_at | *OSMR* | 1,38608371145944 |
| 1569362_at | --- | 1,3854814755998 |
| 1567575_at | --- | 1,38546042902438 |
| 225540_at | *MAP2* | 1,38425341304891 |
| 222719_s_at | *PDGFC* | 1,37177523865547 |
| 218180_s_at | *EPS8L2* | 1,37109560848652 |
| 220238_s_at | *KLHL7* | 1,35869677233041 |
| 227347_x_at | *HES4* | 1,35289398442871 |
| 227984_at | *LMF1* | 1,34703833594236 |
| 39248_at | *AQP3* | 1,3414047523504 |

| | | |
|---|---|---|
| 238919_at | --- | 1,34138638303494 |
| 226129_at | *FAM83H* | 1,33766681963499 |
| 207345_at | *FST* | 1,33088383300439 |
| 240655_at | --- | 1,32603604594769 |
| 235743_at | --- | 1,32437757958722 |
| 209897_s_at | *SLIT2* | 1,31688495777695 |
| 240815_at | --- | 1,31341704979732 |
| 206300_s_at | *PTHLH* | 1,3125667163762 |
| 203896_s_at | *PLCB4* | 1,30900369422726 |
| 219513_s_at | *SH2D3A* | 1,30716643310616 |
| 210365_at | *RUNX1* | 1,29914186236785 |
| 204948_s_at | *FST* | 1,29896555001809 |
| 203765_at | *GCA* | 1,29488628414951 |
| 227425_at | *REPS2* | 1,2939099789793 |
| 223313_s_at | *MAGED4 /// MAGED4B* | 1,29361683450663 |
| 242592_at | *GPR137C* | 1,29255826819502 |
| 230263_s_at | *DOCK5* | 1,2871407512382 |
| 212458_at | *SPRED2* | 1,28590714797483 |
| 239433_at | *LRRC8E* | 1,28523965641412 |
| 226552_at | *IER5L* | 1,28482440121654 |
| 217371_s_at | *IL15* | 1,28177535764805 |
| 224894_at | *YAP1* | 1,28067591850801 |
| 203502_at | *BPGM* | 1,28045106712855 |
| 221529_s_at | *PLVAP* | 1,28016079729247 |
| 201951_at | *ALCAM* | 1,27891265588615 |
| 227396_at | *PTPRJ* | 1,27676442344661 |
| 203184_at | *FBN2* | 1,27673077320683 |
| 237435_at | --- | 1,27299611569676 |
| 205547_s_at | *TAGLN* | 1,27007994342916 |
| 210015_s_at | *MAP2* | 1,26889797907056 |
| 205347_s_at | *TMSB15A* | 1,26860393856157 |
| 227235_at | *GUCY1A3* | 1,26825747400716 |
| 232617_at | *CTSS* | 1,26770196094556 |
| 202668_at | *EFNB2* | 1,2672192347484 |
| 218094_s_at | *DBNDD2 /// SYS1 /// SYS1-DBNDD2* | 1,26515770889517 |
| 1552730_at | *DHH* | 1,26245522895882 |
| 226912_at | *ZDHHC23* | 1,25485256944054 |
| 228933_at | *NHS* | 1,25222553287927 |
| 235295_at | --- | 1,24785770227774 |
| 220239_at | *KLHL7* | 1,24444997894583 |
| 202719_s_at | *TES* | 1,23339739309903 |
| 240173_at | --- | 1,23104338276037 |
| 219371_s_at | *KLF2* | 1,22944888230658 |
| 221648_s_at | *LOC100507192* | 1,2277067877276 |
| 203571_s_at | *C10orf116* | 1,22302076663077 |
| 229292_at | *EPB41L5* | 1,22037524526101 |
| 214587_at | *COL8A1* | 1,21628477802512 |
| 202902_s_at | *CTSS* | 1,21621010160918 |
| 202856_s_at | *SLC16A3* | 1,2123521622323 |
| 241470_x_at | --- | 1,20926218075273 |
| 201041_s_at | *DUSP1* | 1,20529485205423 |
| 238865_at | *PABPC4L* | 1,19788246878757 |
| 228051_at | *KIAA1244* | 1,19722920357872 |
| 226497_s_at | *FLT1* | 1,19667439829071 |
| 228877_at | *RGL3* | 1,18911569218785 |
| 205560_at | *PCSK5* | 1,18541138299403 |
| 207777_s_at | *SP140* | 1,18344904253659 |
| 224895_at | *YAP1* | 1,18293098796678 |
| 1553519_at | *C21orf94* | 1,18185924691068 |
| 221577_x_at | *GDF15* | 1,18112415854271 |
| 210002_at | *GATA6* | 1,18110720743387 |
| 239973_at | --- | 1,18066582653083 |
| 219863_at | *HERC5* | 1,17984680809036 |
| 241790_at | --- | 1,1787416265767 |
| 1559360_at | --- | 1,17689460949067 |
| 224952_at | *TANC2* | 1,17520584630956 |
| 206987_x_at | *FGF18* | 1,1734842889741 |
| 35666_at | *SEMA3F* | 1,17290627424217 |
| 213342_at | *YAP1* | 1,17126252040403 |
| 1569872_a_at | *LMF1* | 1,17074590328645 |
| 227107_at | --- | 1,16927244135503 |
| 210335_at | *RASSF9* | 1,16892679537721 |
| 242273_at | --- | 1,16647931747563 |

### Supplementary Table S3 (continued)

| | | |
|---|---|---|
| 241763_s_at | --- | 1,164663322194 |
| 228151_at | --- | 1,16407664869185 |
| 201565_s_at | ID2 | 1,16259196196948 |
| 203126_at | IMPA2 | 1,15962398388415 |
| 223374_s_at | B3GALNT1 | 1,15934054356378 |
| 207303_at | PDE1C | 1,15741327242352 |
| 227892_at | PRKAA2 | 1,15725741869748 |
| 227371_at | BAIAP2L1 | 1,15417079386819 |
| 228523_at | NANOS1 | 1,15383861347929 |
| 219377_at | FAM59A | 1,14883776747832 |
| 203395_s_at | HES1 | 1,14668517889227 |
| 211379_x_at | B3GALNT1 | 1,1453488560117 |
| 239028_at | LYPD6 | 1,14122690155769 |
| 204715_at | PANX1 | 1,13997217623929 |
| 215498_s_at | MAP2K3 | 1,13745557356997 |
| 213493_at | SNED1 | 1,13523827808447 |
| 230207_s_at | DOCK5 | 1,1321223348103 |
| 239367_at | BDNF | 1,13132985850448 |
| 229310_at | KLHL29 | 1,13037901021091 |
| 213112_s_at | SQSTM1 | 1,1302522960711 |
| 223378_at | GLIS2 | 1,1299784633564 |
| 223525_at | DLL4 | 1,12825335588383 |
| 224851_at | CDK6 | 1,1252379132885 |
| 210287_s_at | FLT1 | 1,12509984164314 |
| 205207_at | IL6 | 1,12311588988254 |
| 205780_at | BIK | 1,11774534556943 |
| 209185_s_at | IRS2 | 1,11592400332637 |
| 232596_at | DIAPH3 | 1,11416376037245 |
| 235626_at | CAMK1D | 1,11389129399881 |
| 1557415_s_at | LETM2 | 1,11260387795958 |
| 219842_at | ARL15 | 1,11127710604669 |
| 222062_at | IL27RA | 1,10842485650135 |
| 201189_s_at | ITPR3 | 1,10782580688588 |
| 204875_s_at | GMDS | 1,10751264924317 |
| 244845_at | --- | 1,10520032159144 |
| 219573_at | LRRC16A | 1,10482747384966 |
| 1556221_a_at | --- | 1,10407373511476 |
| 236947_at | --- | 1,10370314530597 |
| 231513_at | --- | 1,10215429099993 |
| 218326_s_at | LGR4 | 1,10093226394244 |
| 206448_at | ZNF365 | 1,09960935376579 |
| 202669_s_at | EFNB2 | 1,097167639678 |
| 212057_at | KIAA0182 | 1,09334838111106 |
| 203741_s_at | ADCY7 | 1,09300175234672 |
| 227014_at | ASPHD2 | 1,08806073965916 |
| 226313_at | C10orf35 | 1,08682356204822 |
| 208156_x_at | EPPK1 | 1,08318304145217 |
| 206191_at | ENTPD3 | 1,07816843791093 |
| 226559_at | IER5L | 1,07662649099769 |
| 219932_at | SLC27A6 | 1,07613277430435 |
| 229437_at | MIR155HG | 1,07463400896875 |
| 225060_at | LRP11 | 1,07429205004028 |
| 204501_at | NOV | 1,07403591999113 |
| 206765_at | KCNJ2 | 1,07380503250575 |
| 231856_at | KIAA1244 | 1,07319416279175 |
| 205251_at | PER2 | 1,07211391653402 |
| 222450_at | PMEPA1 | 1,06943094239946 |
| 242963_at | SGMS2 | 1,0683262932032 |
| 235046_at | INPP4B | 1,06650515128907 |
| 211057_at | ROR1 | 1,06559735186803 |
| 221904_at | FAM131A | 1,06024629879698 |
| 212354_at | SULF1 | 1,05912727503174 |
| 206400_at | LGALS7 /// LGALS7B | 1,05753439698542 |
| 222890_at | CCDC113 | 1,0569517898674 |
| 203836_s_at | MAP3K5 | 1,05605080728225 |
| 221942_s_at | GUCY1A3 | 1,05519093975861 |
| 211029_x_at | FGF18 | 1,05426742679913 |
| 206417_at | CNGA1 | 1,05414587302639 |
| 210220_at | FZD2 | 1,05211493680679 |
| 203845_at | KAT2B | 1,04790192773297 |
| 200706_s_at | LITAF | 1,04710172523668 |
| 205542_at | STEAP1 | 1,04598564877054 |
| 218376_s_at | MICAL1 | 1,04258200407638 |
| 225867_at | VASN | 1,04253652785592 |

| | | |
|---|---|---|
| 205691_at | SYNGR3 | 1,04244960182686 |
| 205463_s_at | PDGFA | 1,03921599452774 |
| 1555724_s_at | TAGLN | 1,03844924179716 |
| 204602_at | DKK1 | 1,03829502557181 |
| 228724_at | TTLL7 | 1,03824868505957 |
| 227337_at | ANKRD37 | 1,0364706288428 |
| 227384_s_at | LOC727820 | 1,03580833780531 |
| 203837_at | MAP3K5 | 1,03471413550889 |
| 203788_s_at | SEMA3C | 1,03452784586 |
| 240349_at | PRKAA2 | 1,03364299929001 |
| 204938_s_at | PLN | 1,03353848356876 |
| 229830_at | PDGFA | 1,0333390284017 |
| 242541_at | ABCA9 | 1,03100650355222 |
| 243000_at | CDK6 | 1,03009080652525 |
| 209909_s_at | TGFB2 | 1,02838429479328 |
| 229228_at | CREB5 | 1,02743408058123 |
| 236114_at | --- | 1,02620115756836 |
| 210044_s_at | LYL1 | 1,02592185035654 |
| 201952_at | ALCAM | 1,02447822964285 |
| 203085_s_at | TGFB1 | 1,02444653545872 |
| 230206_at | DOCK5 | 1,0216822072303 |
| 229011_at | --- | 1,02155105886185 |
| 210102_at | VWA5A | 1,01906022719071 |
| 230100_x_at | PAK1 | 1,01881042895358 |
| 206910_x_at | CFHR2 | 1,01009938577111 |
| 233417_at | --- | 1,00922328637384 |
| 205297_s_at | CD79B | 1,006726579124 |
| 201170_s_at | BHLHE40 | 1,0048419127324 |
| 227034_at | ANKRD57 | 1,00475894893188 |
| 202855_s_at | SLC16A3 | 1,00411492674708 |
| 203961_at | NEBL | 1,00343179695171 |
| 213424_at | KIAA0895 | 1,00285409294761 |
| 206675_s_at | SKIL | 1,00191640423979 |
| 226608_at | C16orf87 | 1,00170265573231 |
| 229097_at | DIAPH3 | 1,00146788271845 |
| 203835_at | LRRC32 | 1,00047323412054 |
| 1557749_at | EHBP1L1 | 1,0001517511488 |
| 233375_at | EFCAB2 | -1,0021561502709 |
| 229569_at | --- | -1,00224005863062 |
| 231798_at | NOG | -1,0023027657756 |
| 229667_s_at | HOXB8 | -1,00257516516463 |
| 211809_x_at | COL13A1 | -1,00978290758253 |
| 236798_at | --- | -1,01161835790728 |
| 229635_at | --- | -1,01535092783927 |
| 216466_at | NAV3 | -1,0157312291342 |
| 227856_at | C4orf32 | -1,01576800572881 |
| 229178_at | PRTG | -1,01983430971979 |
| 213715_s_at | KANK3 | -1,02049055188899 |
| 1557626_at | --- | -1,02128651492485 |
| 244007_at | ZNF462 | -1,02185766008188 |
| 202877_s_at | CD93 | -1,02455626465202 |
| 204088_at | P2RX4 | -1,02855107694911 |
| 208092_s_at | FAM49A | -1,02911074746624 |
| 210993_s_at | SMAD1 | -1,02919273261842 |
| 209733_at | MID2 | -1,0318911595877 |
| 212665_at | TIPARP | -1,03437808652043 |
| 203434_s_at | MME | -1,04022818898673 |
| 228819_at | TSPAN18 | -1,0403857809669 |
| 241060_x_at | --- | -1,04473596298869 |
| 237411_at | ADAMTS6 | -1,04607606610209 |
| 205698_s_at | MAP2K6 | -1,04958707731892 |
| 212551_at | CAP2 | -1,05152996212815 |
| 241561_at | --- | -1,05183086735778 |
| 1559716_at | INO80C | -1,05370534644992 |
| 205122_at | TMEFF1 | -1,0544198371194 |
| 237034_at | --- | -1,05556072676256 |
| 239719_at | CD109 | -1,05601703550023 |
| 226448_at | FAM89A | -1,05905486664868 |
| 210754_s_at | LYN | -1,06152407276063 |
| 226545_at | CD109 | -1,06250140267968 |
| 229695_at | --- | -1,06340615421195 |
| 213316_at | KIAA1462 | -1,06349253416602 |
| 1554717_a_at | PDE4D | -1,06371120601664 |
| 205479_s_at | PLAU | -1,06442967577418 |
| 228027_at | GPRASP2 | -1,06790874352394 |

### Supplementary Table S3 (continued)

| | | |
|---|---|---|
| 1560396_at | *KLHL6* | -1,06841449957477 |
| 204341_at | *TRIM16* | -1,07280014501307 |
| 223796_at | *CNTNAP3* | -1,07424530278427 |
| 230132_at | --- | -1,07807614418627 |
| 204929_s_at | *VAMP5* | -1,07980651383618 |
| 241679_at | --- | -1,08042710346944 |
| 213256_at | *MARCH3* | -1,08370824978245 |
| 212183_at | *NUDT4* | -1,0842257610461 |
| 222900_at | *NRIP3* | -1,08539442759807 |
| 222303_at | --- | -1,08542478203112 |
| 206481_s_at | *LDB2* | -1,08628580997257 |
| 210842_at | *NRP2* | -1,08733352463373 |
| 219412_at | *RAB38* | -1,09282744812035 |
| 221019_s_at | *COLEC12* | -1,09299909646828 |
| 212554_at | *CAP2* | -1,09307836437403 |
| 239288_at | *TNIK* | -1,09346100267651 |
| 1560031_at | *FRMD4A* | -1,09420640892057 |
| 240254_at | --- | -1,09483445604729 |
| 210138_at | *RGS20* | -1,09624700125675 |
| 210174_at | *NR5A2* | -1,0963601965017 |
| 236089_at | --- | -1,10093260000176 |
| 229331_at | *SPATA18* | -1,10217727048144 |
| 211071_s_at | *MLLT11* | -1,10297582452119 |
| 228101_at | *APBA1* | -1,1055535852444 |
| 244353_s_at | *SLC2A12* | -1,10780292568056 |
| 207808_s_at | *PROS1* | -1,10835688604679 |
| 202555_s_at | *MYLK* | -1,11118696750254 |
| 220753_s_at | *CRYL1* | -1,11235482881292 |
| 228444_at | --- | -1,11263476830763 |
| 240407_at | *LOC100126784* | -1,11304881550526 |
| 227862_at | *TRNP1* | -1,11317018717835 |
| 219400_at | *CNTNAP1* | -1,12268200138839 |
| 214770_at | *MSR1* | -1,12613113122471 |
| 204588_s_at | *SLC7A7* | -1,1308088117051 |
| 204011_at | *SPRY2* | -1,13322007096193 |
| 229759_s_at | *VEPH1* | -1,13542619269894 |
| 224911_s_at | *DCBLD2* | -1,14617433575811 |
| 212298_at | *NRP1* | -1,14653197261858 |
| 211434_s_at | *CCRL2* | -1,14697988994086 |
| 215069_at | *NMT2* | -1,14700347838187 |
| 237016_at | *TMEM217* | -1,15404884728123 |
| 231841_s_at | *KIAA1462* | -1,15530874048118 |
| 206584_at | *LY96* | -1,15711088015735 |
| 225388_at | *TSPAN5* | -1,15878432456236 |
| 225168_at | *FRMD4A* | -1,15942236429615 |
| 219926_at | *POPDC3* | -1,16093099994575 |
| 204404_at | *SLC12A2* | -1,16128494350346 |
| 228977_at | *LOC729680* | -1,16470663978859 |
| 202974_at | *MPP1* | -1,1655574602796 |
| 203585_at | *ZNF185* | -1,16638825607045 |
| 1552487_a_at | *BNC1* | -1,16834383008509 |
| 204780_s_at | *FAS* | -1,1702962844701 |
| 226853_at | *BMP2K* | -1,1740300967848 |
| 239849_at | --- | -1,17514814713627 |
| 228167_at | *KLHL6* | -1,17532052186557 |
| 236304_at | --- | -1,17948632325845 |
| 218729_at | *LXN* | -1,18107260968429 |
| 1555270_a_at | *WFS1* | -1,18215073033408 |
| 227123_at | *RAB3B* | -1,18254597376279 |
| 222043_at | *CLU* | -1,18581249520646 |
| 238169_at | --- | -1,18593057183727 |
| 214438_at | *HLX* | -1,18884829910144 |
| 211679_x_at | *GABBR2* | -1,18933781315909 |
| 209568_s_at | *RGL1* | -1,18987166894618 |
| 201328_at | *ETS2* | -1,19708965928854 |
| 216598_s_at | *CCL2* | -1,19958630639969 |
| 227307_at | *TSPAN18* | -1,20071425167067 |
| 1554485_s_at | *TMEM37* | -1,20342971299417 |
| 243586_at | --- | -1,20356268194244 |
| 232473_at | *PRPF18* | -1,2056193689288 |
| 205123_s_at | *TMEFF1* | -1,20725667566922 |
| 244317_at | *KIAA1324L* | -1,20765286862718 |
| 202626_s_at | *LYN* | -1,20799366455647 |
| 225167_at | *FRMD4A* | -1,21380248176261 |
| 227522_at | *CMBL* | -1,21664130578319 |
| 203562_at | *FEZ1* | -1,22712705384161 |
| 236442_at | *DPF3* | -1,23906706764465 |
| 222885_at | *EMCN* | -1,23966505993931 |
| 203934_at | *KDR* | -1,24190280320907 |
| 205005_s_at | *NMT2* | -1,24197848536929 |
| 237194_at | --- | -1,24339321038639 |
| 227401_at | *IL17D* | -1,24537902738487 |
| 207016_s_at | *ALDH1A2* | -1,24586579913592 |
| 1556331_a_at | --- | -1,24907777952897 |
| 205006_s_at | *NMT2* | -1,25180999326363 |
| 240259_at | *LOC100506718* | -1,25569970220257 |
| 215719_x_at | *FAS* | -1,25950886400547 |
| 209683_at | *FAM49A* | -1,2600755814687 |
| 232080_at | *HECW2* | -1,26023858285852 |
| 225835_at | *SLC12A2* | -1,26168813775116 |
| 219557_s_at | *NRIP3* | -1,26254469943089 |
| 217077_s_at | *GABBR2* | -1,26318955489032 |
| 226575_at | *ZNF462* | -1,26321596561391 |
| 222756_s_at | *ARRB1* | -1,26751420051482 |
| 226955_at | *AFAP1L1* | -1,26911949693157 |
| 238532_at | *DPF3* | -1,27263457038835 |
| 1555929_s_at | --- | -1,27394128423757 |
| 213865_at | *DCBLD2* | -1,27399836228289 |
| 216250_s_at | *LPXN* | -1,27632090390998 |
| 213624_at | *SMPDL3A* | -1,27832356231356 |
| 204058_at | *ME1* | -1,27870781399989 |
| 226028_at | *ROBO4* | -1,28210520769378 |
| 219582_at | *OGFRL1* | -1,2827466928261 |
| 224694_at | *ANTXR1* | -1,28797444371774 |
| 242301_at | *CBLN2* | -1,29021048146154 |
| 212660_at | *PHF15* | -1,2920525636416 |
| 219049_at | *CSGALNACT1* | -1,29538115694333 |
| 225387_at | *TSPAN5* | -1,2968056454213 |
| 219956_at | *GALNT6* | -1,30554114064103 |
| 209652_s_at | *PGF* | -1,30903940915386 |
| 215506_s_at | *DIRAS3* | -1,31454706090299 |
| 212543_at | *AIM1* | -1,32143318678829 |
| 204059_s_at | *ME1* | -1,32254119425067 |
| 205801_s_at | *RASGRP3* | -1,32939791370968 |
| 202908_at | *WFS1* | -1,33029294477844 |
| 49111_at | --- | -1,33314126569304 |
| 244065_at | *LOC643792* | -1,33652106355154 |
| 221861_at | --- | -1,33685700942014 |
| 1555786_s_at | *C14orf34* | -1,33760042460871 |
| 200974_at | *ACTA2* | -1,34767878135522 |
| 205924_at | *RAB3B* | -1,34996175879756 |
| 242509_at | --- | -1,35322470697273 |
| 218660_at | *DYSF* | -1,3569987216126 |
| 220450_at | --- | -1,35791285526176 |
| 205472_s_at | *DACH1* | -1,36196920709057 |
| 210510_s_at | *NRP1* | -1,36250395366689 |
| 206638_at | *HTR2B* | -1,36315775340086 |
| 202625_at | *LYN* | -1,3653789377076 |
| 204781_s_at | *FAS* | -1,37063838546168 |
| 228794_at | *XIRP2* | -1,37492804779713 |
| 214156_at | *MYRIP* | -1,37641725777339 |
| 1552658_a_at | *NAV3* | -1,38136991863341 |
| 209890_at | *TSPAN5* | -1,38587182459602 |
| 235301_at | *KIAA1324L* | -1,38708655920587 |
| 225566_at | *NRP2* | -1,38795216172148 |
| 214632_at | *NRP2* | -1,38833292549885 |
| 215743_at | *NMT2* | -1,38870881334701 |
| 222862_s_at | *AK5* | -1,39081428369128 |
| 236361_at | *GALNTL2* | -1,39204434390031 |
| 231945_at | *FILIP1* | -1,39464796921841 |
| 211844_s_at | *NRP2* | -1,40172364632698 |
| 219436_s_at | *EMCN* | -1,40204375750621 |
| 1556325_at | *FILIP1* | -1,40290729468444 |
| 43511_s_at | --- | -1,40503988232216 |
| 1553798_a_at | *FBXL13* | -1,41344531267555 |
| 239519_at | --- | -1,41380641273288 |
| 205825_at | *PCSK1* | -1,41979022028501 |
| 209543_s_at | *CD34* | -1,41996102169905 |
| 213385_at | *CHN2* | -1,42504236357963 |
| 229354_at | *AHRR* | -1,42988524841155 |

### Supplementary Table S3 (continued)

| | | |
|---|---|---|
| 213992_at | *COL4A6* | -1,43006873229725 |
| 209785_s_at | *PLA2G4C* | -1,44530995880319 |
| 225163_at | *FRMD4A* | -1,45226537572092 |
| 238469_at | *OGFRL1* | -1,45260738008504 |
| 223423_at | *GPR160* | -1,45512644164696 |
| 228568_at | *GCOM1* | -1,45543369241803 |
| 225842_at | *PHLDA1* | -1,46078624785029 |
| 1555928_at | --- | -1,46330143213144 |
| 227874_at | *EMCN* | -1,46874846678941 |
| 204220_at | *GMFG* | -1,47624987331697 |
| 1562367_at | *C15orf54* | -1,47665451423135 |
| 228885_at | *MAMDC2* | -1,47685563108168 |
| 221031_s_at | *APOLD1* | -1,47879905135629 |
| 225810_at | *MTMR10* | -1,48052592668371 |
| 230748_at | *SLC16A6* | -1,48163718603664 |
| 236335_at | --- | -1,48466866530599 |
| 230425_at | *EPHB1* | -1,49410896482337 |
| 202947_s_at | *GYPC* | -1,49685863753355 |
| 226810_at | *OGFRL1* | -1,50894356005804 |
| 222108_at | *AMIGO2* | -1,52552048169533 |
| 235183_at | --- | -1,53103667247033 |
| 228501_at | *GALNTL2* | -1,538128961209 |
| 232701_at | --- | -1,54505662882261 |
| 221371_at | *TNFSF18* | -1,55135225914311 |
| 224724_at | *SULF2* | -1,55230316296082 |
| 232383_at | *TFEC* | -1,55554515546065 |
| 205680_at | *MMP10* | -1,56974177021941 |
| 205471_s_at | *DACH1* | -1,58555136269806 |
| 224823_at | *MYLK* | -1,59437186696439 |
| 217996_at | *PHLDA1* | -1,59543055691667 |
| 206715_at | *TFEC* | -1,61317936729109 |
| 1570515_a_at | *FILIP1* | -1,62089190210735 |
| 228915_at | *DACH1* | -1,62728969684818 |
| 233555_s_at | *SULF2* | -1,63793321888041 |
| 207426_s_at | *TNFSF4* | -1,64140695101795 |
| 211919_s_at | *CXCR4* | -1,65255988639837 |
| 1552489_s_at | *MPP4* | -1,67468499112158 |
| 209990_s_at | *GABBR2* | -1,67663411919333 |
| 219367_s_at | --- | -1,68897000739593 |
| 230061_at | *TM4SF18* | -1,70565498938722 |
| 209201_x_at | *CXCR4* | -1,72574118436593 |
| 1555366_at | *NSAP11* | -1,72671421736163 |
| 228489_at | *TM4SF18* | -1,76780791088656 |
| 222877_at | --- | -1,77114333903588 |
| 217028_at | *CXCR4* | -1,78463681240894 |
| 236383_at | --- | -1,78936554743521 |
| 223510_at | *NRP2* | -1,79206729693627 |
| 204823_at | *NAV3* | -1,83057909154309 |
| 212951_at | *GPR116* | -1,84474103179383 |
| 212950_at | *GPR116* | -1,8640052258768 |
| 219884_at | *LHX6* | -1,87105326265266 |
| 217997_at | *PHLDA1* | -1,91627692119996 |
| 227458_at | *CD274* | -1,92874162329614 |
| 202888_s_at | *ANPEP* | -1,97610752920857 |
| 217999_s_at | *PHLDA1* | -1,98445407868903 |
| 211343_s_at | *COL13A1* | -2,00661391591662 |
| 206211_at | *SELE* | -2,03946743213277 |
| 234279_at | --- | -2,07872099253941 |
| 219059_s_at | *LYVE1* | -2,11877052851679 |
| 204475_at | *MMP1* | -2,15227302383435 |
| 222856_at | *APLN* | -2,1552723681185 |
| 230040_at | *ADAMTS18* | -2,56617927102486 |
| 220037_s_at | *LYVE1* | -2,58172723218289 |
| 237356_at | --- | -2,86209429092656 |

TableS4

### Supplementary Table S4. Gene Ontology (GO) for DEGs from Prdm16+ *versus* control BOECs

| Gene Set Name | # Genes in Gene Set (K) | # Genes in Overlap (k) | k/K | FDR q-value |
|---|---|---|---|---|
| GOBP_ANIMAL_ORGAN_MORPHOGENESIS | 1042 | 20 | 0.0192 | 1.11E-10 |
| GOBP_OSSIFICATION | 443 | 12 | 0.0271 | 3.39E-7 |
| GOBP_CONNECTIVE_TISSUE_DEVELOPMENT | 286 | 10 | 0.0350 | 5.61E-7 |
| GOBP_EPITHELIUM_DEVELOPMENT | 1244 | 17 | 0.0137 | 5.61E-7 |
| GOBP_NEGATIVE_REGULATION_OF_MULTICELLULAR_ORGANISMAL_PROCESS | 1249 | 17 | 0.0136 | 5.61E-7 |
| GOBP_TRABECULA_FORMATION | 26 | 5 | 0.1923 | 1.97E-6 |
| GOBP_CARDIAC_CHAMBER_DEVELOPMENT | 168 | 8 | 0.0476 | 1.97E-6 |
| GOBP_CIRCULATORY_SYSTEM_DEVELOPMENT | 1224 | 16 | 0.0131 | 2.2E-6 |
| GOBP_HEART_MORPHOGENESIS | 265 | 9 | 0.0340 | 2.75E-6 |
| GOBP_HEART_DEVELOPMENT | 618 | 12 | 0.0194 | 2.91E-6 |
| GOBP_CARDIAC_CHAMBER_MORPHOGENESIS | 127 | 7 | 0.0551 | 4.5E-6 |
| GOBP_CARDIAC_VENTRICLE_DEVELOPMENT | 127 | 7 | 0.0551 | 4.5E-6 |
| GOBP_CELL_ADHESION | 1531 | 17 | 0.0111 | 4.54E-6 |
| GOBP_SKELETAL_SYSTEM_DEVELOPMENT | 529 | 11 | 0.0208 | 4.55E-6 |
| GOBP_CARTILAGE_DEVELOPMENT | 206 | 8 | 0.0388 | 4.55E-6 |
| GOBP_SEMI_LUNAR_VALVE_DEVELOPMENT | 47 | 5 | 0.1064 | 1.79E-5 |
| GOBP_TRABECULA_MORPHOGENESIS | 47 | 5 | 0.1064 | 1.79E-5 |
| GOBP_REGULATION_OF_GROWTH | 619 | 11 | 0.0178 | 1.86E-5 |
| GOBP_PULMONARY_VALVE_MORPHOGENESIS | 19 | 4 | 0.2105 | 2.22E-5 |
| GOBP_BIOMINERAL_TISSUE_DEVELOPMENT | 174 | 7 | 0.0402 | 2.38E-5 |
| GOBP_CARDIAC_SEPTUM_DEVELOPMENT | 108 | 6 | 0.0556 | 3.07E-5 |
| GOBP_NEGATIVE_REGULATION_OF_DEVELOPMENTAL_PROCESS | 1004 | 13 | 0.0129 | 3.33E-5 |
| GOBP_PULMONARY_VALVE_DEVELOPMENT | 23 | 4 | 0.1739 | 4.17E-5 |
| GOBP_HEART_VALVE_MORPHOGENESIS | 61 | 5 | 0.0820 | 4.81E-5 |
| GOBP_CHONDROCYTE_PROLIFERATION | 25 | 4 | 0.1600 | 5.47E-5 |
| GOBP_GROWTH | 926 | 12 | 0.0130 | 9.09E-5 |
| GOBP_ATRIOVENTRICULAR_VALVE_DEVELOPMENT | 29 | 4 | 0.1379 | 9.19E-5 |
| GOBP_CARDIAC_SEPTUM_MORPHOGENESIS | 72 | 5 | 0.0694 | 9.19E-5 |
| GOBP_CARDIAC_VENTRICLE_MORPHOGENESIS | 72 | 5 | 0.0694 | 9.19E-5 |
| GOBP_HEART_VALVE_DEVELOPMENT | 73 | 5 | 0.0685 | 9.53E-5 |
| GOBP_VENTRICULAR_SEPTUM_DEVELOPMENT | 74 | 5 | 0.0676 | 9.87E-5 |
| GOBP_NEGATIVE_REGULATION_OF_CELL_POPULATION_PROLIFERATION | 957 | 12 | 0.0125 | 1.05E-4 |
| GOBP_REGULATION_OF_CARDIAC_MUSCLE_CONTRACTION | 78 | 5 | 0.0641 | 1.21E-4 |
| GOBP_TUBE_DEVELOPMENT | 1169 | 13 | 0.0111 | 1.21E-4 |
| GOBP_TISSUE_MORPHOGENESIS | 635 | 10 | 0.0157 | 1.23E-4 |
| GOBP_OUTFLOW_TRACT_MORPHOGENESIS | 82 | 5 | 0.0610 | 1.42E-4 |
| GOBP_REGULATION_OF_CELL_DIFFERENTIATION | 1628 | 15 | 0.0092 | 1.47E-4 |
| GOBP_DEVELOPMENTAL_GROWTH | 660 | 10 | 0.0152 | 1.61E-4 |
| GOBP_VASCULATURE_DEVELOPMENT | 832 | 11 | 0.0132 | 1.62E-4 |
| GOBP_OSTEOBLAST_DIFFERENTIATION | 257 | 7 | 0.0272 | 1.66E-4 |
| GOBP_CARDIAC_CONDUCTION_SYSTEM_DEVELOPMENT | 37 | 4 | 0.1081 | 1.71E-4 |
| GOBP_AORTIC_VALVE_MORPHOGENESIS | 38 | 4 | 0.1053 | 1.86E-4 |
| GOBP_STRIATED_MUSCLE_TISSUE_DEVELOPMENT | 265 | 7 | 0.0264 | 1.9E-4 |
| GOBP_REGULATION_OF_MUSCLE_CONTRACTION | 170 | 6 | 0.0353 | 2.13E-4 |
| GOBP_REGULATION_OF_ACTIN_FILAMENT_BASED_MOVEMENT | 41 | 4 | 0.0976 | 2.37E-4 |
| GOBP_REGULATION_OF_STRIATED_MUSCLE_CONTRACTION | 99 | 5 | 0.0505 | 2.84E-4 |
| GOBP_NEGATIVE_REGULATION_OF_CELL_DIFFERENTIATION | 724 | 10 | 0.0138 | 2.95E-4 |
| GOBP_REGULATION_OF_SYSTEM_PROCESS | 561 | 9 | 0.0160 | 2.95E-4 |
| GOBP_MESONEPHROS_DEVELOPMENT | 103 | 5 | 0.0485 | 3.24E-4 |
| GOBP_POSITIVE_REGULATION_OF_SIGNALING | 1795 | 15 | 0.0084 | 3.64E-4 |
| GOBP_MUSCLE_TISSUE_DEVELOPMENT | 434 | 8 | 0.0184 | 3.88E-4 |
| GOBP_RESPONSE_TO_LAMINAR_FLUID_SHEAR_STRESS | 14 | 3 | 0.2143 | 4.00E-4 |
| GOBP_RESPONSE_TO_CYTOKINE | 948 | 11 | 0.0116 | 4.18E-4 |
| GOBP_ARTERY_DEVELOPMENT | 111 | 5 | 0.0450 | 4.26E-4 |
| GOBP_CIRCULATORY_SYSTEM_PROCESS | 602 | 9 | 0.0150 | 4.56E-4 |
| GOBP_BUNDLE_OF_HIS_CELL_TO_PURKINJE_MYOCYTE_COMMUNICATION | 15 | 3 | 0.2000 | 4.56E-4 |
| GOBP_HEART_TRABECULA_FORMATION | 15 | 3 | 0.2000 | 4.56E-4 |
| GOBP_REGULATION_OF_SMAD_PROTEIN_SIGNAL_TRANSDUCTION | 53 | 4 | 0.0755 | 5.15E-4 |
| GOBP_REGULATION_OF_HEART_CONTRACTION | 209 | 6 | 0.0287 | 5.15E-4 |
| GOBP_RENAL_SYSTEM_DEVELOPMENT | 325 | 7 | 0.0215 | 5.15E-4 |

| | | | | |
|---|---|---|---|---|
| GOBP_EPITHELIAL_CELL_CELL_ADHESION | 16 | 3 | 0.1875 | 5.15E-4 |
| GOBP_HEART_VALVE_FORMATION | 16 | 3 | 0.1875 | 5.15E-4 |
| GOBP_REGULATION_OF_DEVELOPMENTAL_GROWTH | 328 | 7 | 0.0213 | 5.29E-4 |
| GOBP_ACTIN_FILAMENT_BASED_PROCESS | 809 | 10 | 0.0124 | 5.83E-4 |
| GOBP_VENTRICULAR_CARDIAC_MUSCLE_TISSUE_DEVELOPMENT | 58 | 4 | 0.0690 | 6.71E-4 |
| GOBP_ENZYME_LINKED_RECEPTOR_PROTEIN_SIGNALING_PATHWAY | 1022 | 11 | 0.0108 | 6.85E-4 |
| GOBP_CELL_CELL_SIGNALING | 1713 | 14 | 0.0082 | 7.92E-4 |
| GOBP_REGULATION_OF_CELL_DEVELOPMENT | 845 | 10 | 0.0118 | 8.03E-4 |
| GOBP_SUPRAMOLECULAR_FIBER_ORGANIZATION | 847 | 10 | 0.0118 | 8.08E-4 |
| GOBP_CARDIAC_MUSCLE_CONTRACTION | 139 | 5 | 0.0360 | 9.84E-4 |
| GOBP_POSITIVE_REGULATION_OF_CELL_POPULATION_PROLIFERATION | 1286 | 12 | 0.0093 | 9.84E-4 |
| GOBP_NEGATIVE_REGULATION_OF_HORMONE_SECRETION | 67 | 4 | 0.0597 | 1.08E-3 |
| GOBP_NEGATIVE_REGULATION_OF_GROWTH | 247 | 6 | 0.0243 | 1.08E-3 |
| GOBP_REGULATION_OF_MUSCLE_SYSTEM_PROCESS | 247 | 6 | 0.0243 | 1.08E-3 |
| GOBP_REGULATION_OF_ACTIN_FILAMENT_BASED_PROCESS | 380 | 7 | 0.0184 | 1.16E-3 |
| GOBP_HEART_PROCESS | 256 | 6 | 0.0234 | 1.28E-3 |
| GOBP_KIDNEY_EPITHELIUM_DEVELOPMENT | 150 | 5 | 0.0333 | 1.28E-3 |
| GOBP_NEGATIVE_REGULATION_OF_SMAD_PROTEIN_SIGNAL_TRANSDUCTION | 23 | 3 | 0.1304 | 1.28E-3 |
| GOBP_REGULATION_OF_BLOOD_CIRCULATION | 259 | 6 | 0.0232 | 1.33E-3 |
| GOBP_EMBRYO_DEVELOPMENT | 1127 | 11 | 0.0098 | 1.41E-3 |
| GOBP_MESENCHYMAL_CELL_DIFFERENTIATION | 264 | 6 | 0.0227 | 1.44E-3 |
| GOBP_CARDIAC_MUSCLE_CELL_CONTRACTION | 76 | 4 | 0.0526 | 1.57E-3 |
| GOBP_ARTERY_MORPHOGENESIS | 81 | 4 | 0.0494 | 1.99E-3 |
| GOBP_NEGATIVE_REGULATION_OF_LOCOMOTION | 423 | 7 | 0.0165 | 2.05E-3 |
| GOBP_SECRETION | 967 | 10 | 0.0103 | 2.05E-3 |
| GOBP_NEGATIVE_REGULATION_OF_CELL_DEVELOPMENT | 286 | 6 | 0.0210 | 2.11E-3 |
| GOBP_PROSTATE_GLAND_MORPHOGENESIS | 28 | 3 | 0.1071 | 2.11E-3 |
| GOBP_NEGATIVE_REGULATION_OF_SECRETION | 172 | 5 | 0.0291 | 2.19E-3 |
| GOBP_SMAD_PROTEIN_SIGNAL_TRANSDUCTION | 85 | 4 | 0.0471 | 2.25E-3 |
| GOBP_REGULATION_OF_CARDIAC_MUSCLE_CELL_ACTION_POTENTIAL | 29 | 3 | 0.1034 | 2.27E-3 |
| GOBP_REGULATION_OF_TRANSMEMBRANE_RECEPTOR_PROTEIN_SERINE_THREONINE_KINASE_SIGNALING_PATHWAY | 299 | 6 | 0.0201 | 2.58E-3 |
| GOBP_EPITHELIAL_TO_MESENCHYMAL_TRANSITION | 180 | 5 | 0.0278 | 2.61E-3 |
| GOBP_MUSCLE_SYSTEM_PROCESS | 452 | 7 | 0.0155 | 2.83E-3 |
| GOBP_HEART_TRABECULA_MORPHOGENESIS | 32 | 3 | 0.0938 | 2.94E-3 |
| GOBP_STRIATED_MUSCLE_CONTRACTION | 186 | 5 | 0.0269 | 2.94E-3 |
| GOBP_REPRODUCTIVE_SYSTEM_DEVELOPMENT | 309 | 6 | 0.0194 | 2.94E-3 |
| GOBP_REGULATION_OF_MULTICELLULAR_ORGANISMAL_DEVELOPMENT | 1484 | 12 | 0.0081 | 2.94E-3 |
| GOBP_NEGATIVE_REGULATION_OF_CHONDROCYTE_PROLIFERATION | 5 | 2 | 0.4000 | 3.04E-3 |
| GOBP_ANATOMICAL_STRUCTURE_FORMATION_INVOLVED_IN_MORPHOGENESIS | 1259 | 11 | 0.0087 | 3.13E-3 |
| GOBP_RESPONSE_TO_FLUID_SHEAR_STRESS | 34 | 3 | 0.0882 | 3.33E-3 |

TableS5

### Supplementary Table S5. KEGG pathway analysis for DEGs from Prdm16+ *versus* control BOECs

| Gene Set Name | # Genes in Gene Set (K) | Description | # Genes in Overlap (k) | k/K | FDR q-value |
|---|---|---|---|---|---|
| **KEGG_MAPK_SIGNALING_PATHWAY** | 267 | MAPK signaling pathway | 13 | 0.0487 | 4.47E-6 |
| **KEGG_TGF_BETA_SIGNALING_PATHWAY** | 86 | TGF-beta signaling pathway | 8 | 0.0930 | 8.51E-6 |
| **KEGG_GAP_JUNCTION** | 90 | Gap junction | 7 | 0.0778 | 1.25E-4 |
| **KEGG_CYTOKINE_CYTOKINE_RECEPTOR_INTERACTION** | 265 | Cytokine-cytokine receptor interaction | 10 | 0.0377 | 4.54E-4 |
| **KEGG_GNRH_SIGNALING_PATHWAY** | 101 | GnRH signaling pathway | 6 | 0.0594 | 1.72E-3 |
| **KEGG_PATHWAYS_IN_CANCER** | 325 | Pathways in cancer | 10 | 0.0308 | 1.72E-3 |
| **KEGG_LONG_TERM_DEPRESSION** | 70 | Long-term depression | 5 | 0.0714 | 2.18E-3 |
| **KEGG_MELANOMA** | 71 | Melanoma | 5 | 0.0704 | 2.18E-3 |
| **KEGG_VASCULAR_SMOOTH_MUSCLE_CONTRACTION** | 115 | Vascular smooth muscle contraction | 6 | 0.0522 | 2.18E-3 |
| **KEGG_AXON_GUIDANCE** | 129 | Axon guidance | 6 | 0.0465 | 3.68E-3 |
| **KEGG_REGULATION_OF_ACTIN_CYTOSKELETON** | 213 | Regulation of actin cytoskeleton | 7 | 0.0329 | 8.19E-3 |
| **KEGG_JAK_STAT_SIGNALING_PATHWAY** | 155 | Jak-STAT signaling pathway | 6 | 0.0387 | 8.19E-3 |
| **KEGG_PURINE_METABOLISM** | 159 | Purine metabolism | 6 | 0.0377 | 8.64E-3 |
| **KEGG_CHRONIC_MYELOID_LEUKEMIA** | 73 | Chronic myeloid leukemia | 4 | 0.0548 | 1.71E-2 |
| **KEGG_PHOSPHATIDYLINOSITOL_SIGNALING_SYSTEM** | 76 | Phosphatidylinositol signaling system | 4 | 0.0526 | 1.86E-2 |
| **KEGG_FOCAL_ADHESION** | 199 | Focal adhesion | 6 | 0.0302 | 2.22E-2 |
| **KEGG_HYPERTROPHIC_CARDIOMYOPATHY_HCM** | 83 | Hypertrophic cardiomyopathy (HCM) | 4 | 0.0482 | 2.27E-2 |
| **KEGG_DILATED_CARDIOMYOPATHY** | 90 | Dilated cardiomyopathy | 4 | 0.0444 | 2.87E-2 |
| **KEGG_CIRCADIAN_RHYTHM_MAMMAL** | 13 | Circadian rhythm - mammal | 2 | 0.1538 | 3.01E-2 |
| **KEGG_NOTCH_SIGNALING_PATHWAY** | 47 | Notch signaling pathway | 3 | 0.0638 | 3.23E-2 |
| **KEGG_INTESTINAL_IMMUNE_NETWORK_FOR_IGA_PRODUCTION** | 48 | Intestinal immune network for IgA production | 3 | 0.0625 | 3.26E-2 |
| **KEGG_INOSITOL_PHOSPHATE_METABOLISM** | 54 | Inositol phosphate metabolism | 3 | 0.0556 | 4.34E-2 |
| **KEGG_STEROID_HORMONE_BIOSYNTHESIS** | 55 | Steroid hormone biosynthesis | 3 | 0.0545 | 4.37E-2 |
| **KEGG_CALCIUM_SIGNALING_PATHWAY** | 178 | Calcium signaling pathway | 5 | 0.0281 | 4.66E-2 |
| **KEGG_ARACHIDONIC_ACID_METABOLISM** | 58 | Arachidonic acid metabolism | 3 | 0.0517 | 4.66E-2 |
| **KEGG_ENDOCYTOSIS** | 181 | Endocytosis | 5 | 0.0276 | 4.66E-2 |

TableS6

#### Supplementary Table S6. List of primers for qRT-PCR and morpholino sequences

| Gene | Forward primer (5'-3') | Reverse primer (5'-3') |
|---|---|---|
| | ***Human*** | |
| *PRDM16* | CAAACGCTTCGAATGTGAAA | CGTGTAGGACTTGTGGCAGA |
| *DLL4* | ATGACCACTTCGGCCACTATG | GCCCGAAAGACAGATAGGCTG |
| *HEY1* | GAGAAGGCTGGTACCCAGTG | GCTCAGATAACGCGCAACTT |
| *HEY2* | CCCCTGCGAGGAGACGA | ATCTAATCACAGAGCTAGTACTTTGCCC |
| *EFNB2* | CTGCTGGATCAACCAGGAATTCC | GGTACTTCAGCAAGAGG |
| *COUP-TFII* | CGCCTCAAAAAGTGCCTCA | GCATCCTGCCCCTCTGC |
| *EPHB4* | GAGCTGTGTGGCAATCAAGA | GAACTGTCCGTCGTTTAGCC |
| *GJA5* | TACAACACAAGGCAGCAAGC | TCGTATCACACCGGAAATCA |
| *ATP2A3* | AGGCCAAGGGTGTGTATGAG | TGCAGTACACGGACATGGAT |
| *DKK2* | AGTACCCGCTGCAATAATGG | GAAATGACGAGCACAGCAAA |
| *FLRT2* | CCTCCTGACCAACAAGGGTA | GGTTATTCCGAGCAGTGAGC |
| *EMCN* | TTGTCACACCAACAACTGGAA | TGTTGAAACAGCATTTGGAAG |
| *GAPDH* | TGGTATCGTGGAAGGACTCATGAC | ATGCCAGTGAGCTTCCCGTTCAGC |
| *β-ACTIN* | TGGCACCACACCTTCTACAATG | TAGCAACGTACATGGCTGGG |
| | ***Mouse*** | |
| *Prdm16* | CACGGAAGAGCGTGAGTACA | GGCAGACCTGCGATATGAAT |
| *Coup-TFII* | CTGGTTAGTAGCCGCTCTGG | TCCCAGTCCTTGCCATTTAG |
| *Hey2* | GCGTCGGGATCGAATAAATA | AGCATGGGCATCAAAGTAGC |
| *β-actin* | GACGGCCAGGTCATCACTAT | CTTCTGCATCCTGTCAGCAA |
| | ***Zebrafish*** | |
| *prdm16* | AGCAGCCATCGCTTCACTAT | GGAGCATCCAGTTTCAGAGC |
| *ef1α* | TCACTGGTACTTCTCAGGCTGACT | TTCTTGGAGATACCAGCCTCAAA |
| | ***Morpholino sequences (5'-3')*** | |
| *prdm16* Mo | TCTCGCCCTCGCCTTGGATCTCATC | |
| *prdm16* Mo2 | TCATTCGCTGTCTTCCCGTCTCCTG | |
| *Dll4* Mo | AAAGGTGAGCCAAGCTGCCATGCTT | |
| *Rbpj* Mo | CAAACTTCCCTGTCACAACAGGCGC | |
| *Hey2* Mo | CGCGCAGGTACAGACACCAAAAACT | |
| nc | GCCATGACATAAGATCCAAGCCAAC | |
